# Genetic architectures of floral pigment and patterning in hybrid monkeyflowers

**DOI:** 10.1101/2022.04.29.490035

**Authors:** Arielle M. Cooley, Emily S. G. Simmons, Caroline Schlutius, Melia Matthews, CiCi Xingyu Zheng, Daniel Thomas, Patrick P. Edger, Adrian E. Platts, Amy M. LaFountain, Logan George, Aaron Williams, Douglas Hundley, Greg Conradi Smith, Yao-Wu Yuan, Alex D. Twyford, Joshua R. Puzey

**Affiliations:** Biology Dept, Whitman College, Walla Walla WA, USA; Biology Dept, William & Mary, Williamsburg VA, USA; Department of Horticulture, Michigan State University, East Lansing, MI, USA; Dept of Ecology & Evolutionary Biology, University of Connecticut, Storrs CT, USA; Math Dept, Whitman College, Walla Walla WA, USA; Royal Botanic Garden Edinburgh, 20a Inverleith Row, Edinburgh, Scotland, UK; Institute of Ecology and Evolution, School of Biological Sciences, University of Edinburgh, Edinburgh, UK; Department of Ecological Microbiology, Bayreuth Center of Ecology and Environmental Research, University of Bayreuth; Dept of Applied Science, School of Computing, Data Sciences, and Physics, William & Mary, Williamsburg VA, USA

**Author notes:** Corresponding author, 345 Boyer Ave., Walla Walla WA 99362 U.S.A.

**Keywords:** anthocyanins, carotenoid intensity, color patterning, digital image analysis, *Erythranthe*, floral pigmentation, genetic architecture, hybrid novelty, hybridization, *Mimulus*, monkeyflower, spatial patterning, reaction-diffusion, positional specification

## Abstract

Coloration in living organisms varies in hue, saturation, and pattern. Here we use genetic mapping to investigate all three elements in monkeyflowers *Mimulus luteus* var. *variegatus* and *M. cupreus*, whose flower petals differ in both yellow carotenoid and magenta anthocyanin pigmentation and whose hybrids exhibit anthocyanin pattern variation not seen in either parent. We report two QTLs associated with carotenoid intensity, and show that lighter yellow petals accumulate relatively high proportions of beta-carotene at the expense of downstream carotenoids. We propose that the derived loss of carotenoid saturation in *M. l. variegatus* is due to a function-reducing mutation in a *Beta-Carotene Hydroxylase* candidate gene, coupled with a reduction in the availability of carotenoid-storing chromoplasts influenced by the *ORANGE* candidate gene. We next identify five QTLs associated with the spatial patterning of anthocyanin pigment. Two QTLs, 9b and 23, each contain genes in the anthocyanin-activating subgroup of the *MYB* family. We hypothesize that *MYB5a/NEGAN*, at QTL-9b, activates spot formation in hybrids as part of a Turing-type reaction-diffusion system, while a MYB at QTL-23 activates solid, unpatterned anthocyanin pigment and may also be able to activate the *variegatus* allele of *MYB5a/NEGAN*. In addition to the apparently stochastic placement of most spots (consistent with a Turing mechanism), some pattern aspects – notably spots at the petal tips – appear to conform to a positional specification model in which spot location is predetermined. Collectively, this work begins to identify how combinations of yellow and red hues, shifts in carotenoid intensity, and two cryptic patterning systems all contribute to the complexity of color divergence between two close relatives.

## Introduction

Plant pigmentation has long served as a vehicle for investigating genomic, developmental, and evolutionary mechanisms (McClintock 1950; Sobel and Streisfeld 2013; Grünig *et al*. 2026). Pigmentation in flowers consists primarily of the yellow-to-orange carotenoids and the red-to-purple anthocyanins (Grotewold 2006). Both types of pigment frequently vary in hue and saturation, while anthocyanins are additionally known for their ability to create visual patterns on flowers and leaves (Davies *et al*. 2012).

The origin of patterns in nature has long mystified both biologists and mathematicians. One of the simplest explanations was proposed by Alan Turing (1952), and gave rise to a class of self-organizing reaction-diffusion equations commonly referred to as Turing systems. In developmental biology these models describe gene regulatory networks, which can contain as few as two interacting gene products and generate spatially repeated patterns such as the spots of the cheetah or the stripes of the zebra (Hunter 2023). In a Turing-type reaction-diffusion system, for example, small stochastic fluctuations in activator concentration are selectively amplified through a positive feedback loop, leading to spot formation at random locations within a patterned area (Gierer and Meinhardt 1972). The activator also activates its own repressor, which diffuses outward past the growing spot and creates a ring of inhibition that limits further spot growth. A competing explanation for pattern formation is positional specification, in which landmarks or coordinates within a tissue determine where a trait is produced (Wolpert 1969). A hallmark of positional specification is that trait locations are determined by positional cues, rather than emerging spontaneously through self-organization.

An attractive biological system for studying the evolution of plant pigmentation, including color patterning, is the monkeyflower genus *Mimulus* (synonym *Erythranthe* (Barker *et al*. 2012; Lowry *et al*. 2019)). Anthocyanin and carotenoid pigmentation vary across *Mimulus* (Wu *et al*. 2008; Sobel and Streisfeld 2013; Yuan 2019), although many of the complex color patterns are found in species that are rare, unstudied, or both. One exception is the *luteus* species group from Chile (Grant 1924; Watson and Von Bohlen 2000), which combines ease of growth in the greenhouse, a solid foundation of prior ecological and pigmentation research (Medel *et al*. 2007; Cooley *et al*. 2011; Esterio *et al*. 2013), ease of hybridization (Stanton *et al*. 2016), and an intriguing hybrid-specific petal pattern phenotype.

The *luteus* group is ancestrally characterized by yellow flowers displaying red dots of anthocyanin pigment in the nectar guide region (Grant 1924). More recently, both carotenoids and anthocyanins have evolved dramatically within the group (Fig. 1a). The result is a range of petal colors including yellow, orange, red, and magenta, depending on the abundance of yellow carotenoid pigments versus magenta anthocyanin pigment (Cooley *et al*. 2008). The group has ancient allopolyploid origins (Mukherjee and Vickery 1962; Vickery *et al*. 1968; Vickery 1995; Edger *et al*. 2017), but traits and markers segregate in a simple diploid fashion (Cooley and Willis 2009).

**Figure 1.**
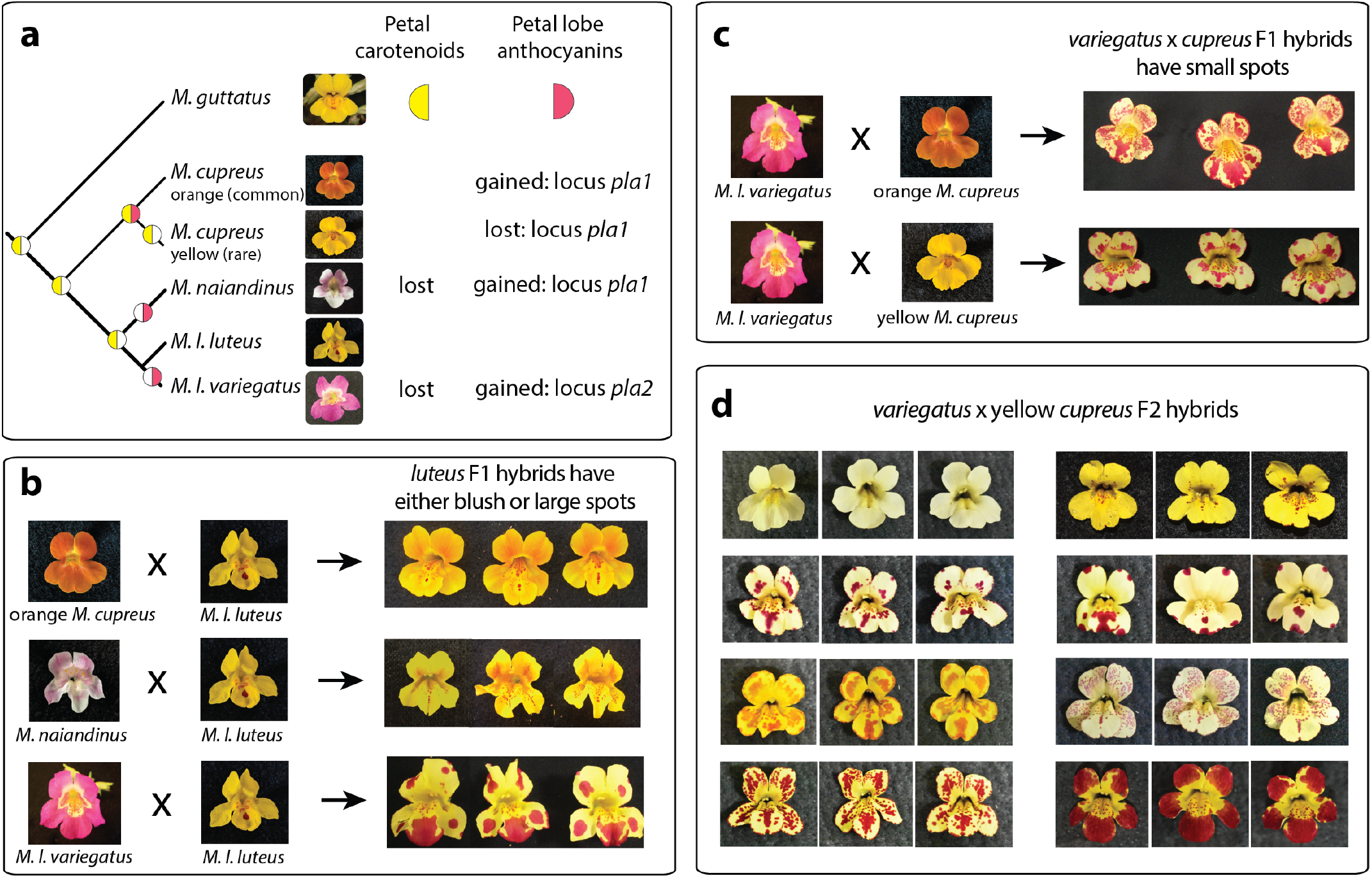
Flower color divergence yields complex spatial patterning in *Mimulus* hybrids. (a) Because the *luteus* group is nested within a large monophyletic grouping of yellow-flowered species, the ancestral state is inferred to consist of yellow, carotenoid-pigmented petals (indicated by yellow color in the left side of the circle) with no petal lobe anthocyanins (indicated by a lack of magenta color in the right side of the circle). Carotenoids are hypothesized to have been lost twice independently, while anthocyanins are hypothesized to have been gained three times and lost once (Cooley and Willis 2009). (b) Crosses within the *luteus* group yield either faint “blush” phenotypes or solid spots of anthocyanin pigmentation in the F1 hybrids. Crosses are shown with maternal parent first; no differences were observed between reciprocal crosses. Three flowers per plant are shown for each F1 hybrid to illustrate the degree of consistency across flowers of the same genotype. (c) *M. l. variegatus* crossed with either the orange or yellow morph of *M. cupreus* generates F1 hybrids with anthocyanin spots. (d) Three flowers per plant are shown for each of 8 different F2 hybrids derived from the *M. l. variegatus* x yellow-flowered *M. cupreus* cross. Photos courtesy Bella Rivera, Joshua Shin, Leah Samuels.

Petal carotenoids have been lost or greatly reduced in *M. naiandinus* and *M. l. variegatus*. Petal Lobe Anthocyanin (PLA), meanwhile, has been gained in *M. naiandinus, M. l. variegatus*, and the common orange morph of *M. cupreus*. In each case, PLA is dominant and segregates in a 3:1 ratio in an F2 population (Cooley and Willis 2009). PLA has been gained at least twice independently, evidenced by its mapping to two different genomic regions – *pla1* for *M. naiandinus* and the orange morph of *M. cupreus*, and the unlinked *pla2* for *M. l. variegatus* (Cooley and Willis 2009; Cooley *et al*. 2011). Candidate genes at both *pla1* and *pla2* consist of a tandem array of R2R3 MYB transcription factor genes belonging to Subgroup 6, which activates the anthocyanin biosynthetic pathway across angiosperms (Stracke *et al*. 2001).

Genetic mapping (Cooley *et al*. 2011) and functional genetic experiments (Zheng *et al*. 2021; Orr *et al*. 2025) show that expanded expression of *MYB5a/NEGAN*, at *pla2*, is responsible for the gain of PLA in *M. l. variegatus*. Crosses show that the *pla1* locus, containing *MYB* genes 1–3 and associated with the gain of petal anthocyanins in the orange morph of *M. cupreus*, is also responsible for secondary loss of PLA in a rare yellow morph of *M. cupreus* (Cooley and Willis 2009).

Hybridization reveals an unexpected effect of the separate evolution of petal anthocyanins: hybrids of the two solid-colored species *M. l. variegatus* x *M. cupreus* are characterized by a spatially complex distribution of petal-lobe anthocyanin spots not seen in either parent (Fig. 1). The spotted pattern is seen regardless of whether the cross involves the common orange morph of *M. cupreus* (high carotenoids and high anthocyanins), or the rare yellow morph (high carotenoids but no petal lobe anthocyanins).

Anthocyanin spots are characteristic of many monkeyflower species, but they are almost always restricted to the nectar guide region of the central petal (Grant 1924). In congeners *Mimulus guttatus* and *M. lewisii*, these spots are thought to be created by a Turing-type reaction-diffusion system, in which the activator of nectar guide spots is *MYB5a/NEGAN* (Ding *et al*. 2020). Like the nectar guide spots, PLA is activated by *MYB5a/NEGAN*, and *variegatus* x *cupreus* hybrids show a stochastic pattern in which flowers from the same plant have similar but not identical spot locations – a characteristic prediction of Turing-type patterning (Turing 1952). An argument against the Turing hypothesis is that PLA in both *M. l. variegatus* and orange *M. cupreus* appears to be solid rather than patterned. However, our previous mathematical modeling has identified variation in interaction strengths and diffusion rates that allows unpatterned parent species to produce patterns when their divergent Turing systems interact in the genome of an F1 or F2 hybrid (Simmons *et al*. 2023). We therefore hypothesized that the *MYB5a/NEGAN* nectar guide spotting system characterized in Ding *et al*. (2020) has expanded outward to activate anthocyanin in the petal lobes of *M. l. variegatus*, while *MYB1–3/PELAN* has done the same in the orange morph of *M. cupreus*.

Here we use newly developed digital tools for quantifying color patterning, and a new *M. l. luteus* reference genome, to genetically map the petal carotenoid and anthocyanin differences between *M. l. variegatus* and *M. cupreus*. We identify two carotenoid-intensity QTLs and five anthocyanin patterning QTLs, including one encompassing *MYB5a/NEGAN* and one that includes the *MYB1–3/PELAN* gene cluster. Overall, this work begins to reveal the components that collectively generate both species divergence and striking hybrid-specific complexity in monkeyflower coloration.

## Materials and Methods

### Plant materials and growth conditions

Two highly inbred lines of *Mimulus luteus* var. *variegatus* RC6 and *M. cupreus* LM43 (Table 1) were developed through thirteen and ten generations of single-seed descent respectively. Multiple individuals of each inbred line were grown in the Whitman College greenhouse under 14-hour days with daily misting from an automatic watering system. Temperatures ranged from approximately 10–15 °C overnight and 15–30 °C during the day. Pollen from *M. cupreus* LM43 plants was applied to *M. l. variegatus* stigmas to generate F1 hybrid seeds. F1 plants were raised in the Whitman College greenhouse and manually self-fertilized to generate F2 hybrid seeds. The cross to generate F1 plants was performed in only one direction because previous work indicated that cross direction does not influence color patterning traits in either an F1 or F2 hybrid population of *variegatus* x *cupreus* (Cooley and Willis 2009). F2 plants were grown in two locations: the Whitman College greenhouse and the William & Mary greenhouse. At the latter location, plants were grown under a 16-hour light regimen at 18–25 °C. *Mimulus l. luteus* line EY7 was grown at William & Mary and was used for the preparation of a reference genome.

**Table 1.** Seed sources.

| Taxon | Line ID | Inbreeding | Population | Location<sup>a</sup> |
| --- | --- | --- | --- | --- |
| *M. luteus* var. *luteus* | Mll-EY7 | 13 generations | El Yeso | 33.4°S, 70.0°W (2600 m) |
| *M. luteus* var. *variegatus* | Mlv-RC6 | 13 generations | Río Cipreses | 34.2°S, 70.3°W (1200 m) |
| *M. cupreus* yellow morph | Mcu-LM43 | 10 generations | Laguna del Maule | 36.0°S, 70.3°W (2300 m) |

<sup>a</sup> Locations from which seeds were collected is given as latitude, longitude (meters above sea level).

### Assembly of the M. l. luteus reference genome

To improve the assembly of the draft *M. l. luteus* genome (Edger *et al*. 2017), we generated new long-read and long-range genomic sequencing data. Using *Mimulus luteus* var. *luteus* line EY7, we first reduced levels of secondary compounds by maintaining plants in the dark for several days, then extracted high molecular weight DNA. The DNA was sequenced using the Sequel II platform to obtain approximately 35x genome-wide PacBio HiFi coverage. A HiC library was constructed and sequenced to approximately 100x, using 150 bp short reads generated via Illumina NovaSeq, to aid in scaffolding. The primary assembly was constructed using Hifiasm (https://github.com/chhylp123/hifiasm; Cheng *et al*. (2021)) with primarily default parameters (except -D 10) to assemble PacBio highly accurate long reads (HiFi reads) into a set of contigs with a low probability of chimerism between subgenomes.

Next, HiC reads were aligned to the primary assembly with bwa (mem) and processed with SALSA 2.2 (https://github.com/marbl/SALSA) to generate a scaffolded assembly. Analysis of synteny (COGE synmap) with the diploid *M. guttatus* suggested that read alignment included some misalignments between subgenomes, and this had been incorporated into the scaffolding. We therefore moved away from bwa to novoalign for HiC alignments with parameters (-r None -t 30) selected to reject any alignments with even a low chance of alignment ambiguity.

To aid in annotation, RNA was extracted from young floral and vegetative buds and sequenced using PacBio IsoSeq. The resulting assembly was annotated using two rounds of Maker with a transcriptome input generated using StringTie (https://ccb.jhu.edu/software/stringtie/) in reference-guided long-read mode based on a set of *M. l. luteus* IsoSeq reads, together with the *M. guttatus* v5.0 proteome from Phytozome 12 (https://phytozome.jgi.doe.gov/pz/portal.html) and a repeat library generated on the assembly using RepeatModeler (https://www.repeatmasker.org/RepeatModeler/). HMMs for Augustus (http://bioinf.uni-greifswald.de/augustus/) and SNAP (https://github.com/KorfLab/SNAP) were initially trained on *M. guttatus* gene models and then retrained on the first round *M. l. luteus* maker gene models, while geneMark used the standard eukaryotic gene models. Genome statistics were generated using the Assemblathon stats (https://github.com/KorfLab/Assemblathon/) code and assembly/annotation completeness was assessed using BUSCO v4 (https://busco.ezlab.org/) with Eudicot odb10 models in genome and proteome modes, respectively. Completeness assessment with the recently released BUSCO v5 code did not significantly change these results.

### Petal photography and qualitative trait assessment

Each *Mimulus* flower consists of five petals: two dorsal (upper), two lateral, and one ventral (lower) petal (Fig. 1). In many *Mimulus* species, including those studied here, the lower petal differs from the other four in that it has a series of anthocyanin spots leading from the petal lobe into the “nectar guide” region of the throat.

We examined both upper petals of each flower, considering them to be approximate (though mirror image) replicates of the same pattern. We also examined the lower petal, hypothesizing that the genes that pattern the nectar guide region might alter how color patterns are produced throughout this unique petal, compared to any of the other petals.

From each of 2–4 flowers per F2 plant, the two upper petals and the lower petal were cut at the base within 48 hours of flower opening. Petals were placed on a strip of white tape and photographed in a darkroom. A single 60-watt soft white (2700K) bulb was used to provide illumination. Each photo included a color standard and a ruler. Photographs were taken using an Olympus VG-120 digital camera (Whitman College) and a Nikon D3200 (William & Mary). Each photograph was cropped and rotated in Gimp v. 2.10.8 (https://www.gimp.org/) so that only the tape background and petals were visible. The resulting .jpg image was processed using the digital image analysis pipeline described below.

Using the photographs, nine qualitative traits were scored based on visual assessment by a single individual (Table 2). For these traits, a single score was assigned to each flower rather than to each petal. Petal Lobe Anthocyanin was scored as present if anthocyanin pigment was robustly visible in the flat, lobe portion of the petal, regardless of whether that pigment was patterned in spots or in a thin layer that we called blush (Table 2). To evaluate repeatability, a set of 100 images was later scored by a second individual for four representative traits. Images were scored identically for the binary traits of blush, globular spray, and fine spray in 85, 92, and 87 out of 100 images, respectively. The fourth tested trait, carotenoid value, had three levels and yielded identical scores in only 66 out of 100 cases, suggesting a lower degree of repeatability.

**Table 2.** Qualitative traits utilized in genetic mapping.

| Trait name | Description |
| --- | --- |
| Carotenoid Value | Darkness of yellow petal pigment, scored as low=0, medium=0.5, high=1. |
| Petal Lobe Anthocyanin (PLA) | Binary indicator of whether any anthocyanin pigment is present (1) or absent (0) on the petal lobes outside of the nectar guide region. |
| Blush | Binary indicator of whether any of the petals have (1) or do not have (0) a very thin layer of diffuse anthocyanin pigment, typically strongest at the base of the petal. |
| Tip Spot | Binary indicator of whether the two upper (dorsal) petals have (1) or do not have (0) a single spot at the tip of the petal. |
| Rim Spots | Amount of rim covered by spots, scored as low=0, medium=0.5, or high=1. |
| Huge Spot | Binary indicator of whether there is (1) or is not (0) a spot that covers nearly all of the petal. |
| Column | Binary indicator of whether the majority of the central column of the lower (ventral) petal is (1) or is not (0) anthocyanin pigmented. |
| Fine Spray | Binary indicator of whether there is (1) or is not (0) a spray of small spots, coming up from the throat on the two upper (dorsal) petals. |
| Globular spray | Binary indicator of whether there is (1) or is not (0) a spray of large spots coming up from the throat on the two upper (dorsal) petals. |

### Digital image analysis pipeline

To enhance downstream analysis of flower photos, full-color photos were first transformed to a 3-color space (l*a*b (Standard *et al*. 2007)), using k-means clustering with 3 centroids: red for anthocyanin-pigmented petal tissue, yellow for non-anthocyanin-pigmented petal tissue, and white for the background (Fig. 2). Using a custom script in combination with the MATLAB plotter, clustering was supervised with manual repositioning of color centroids to ensure optimum retention of detail of spot shapes. Color categorization was implemented in MATLAB version 2017b (MATLAB 2017). One trait – proportion red – was calculated directly in MATLAB for each petal, then averaged across petals and flowers to obtain a single upper petal value and a single lower petal value per plant.

**Figure 2.**
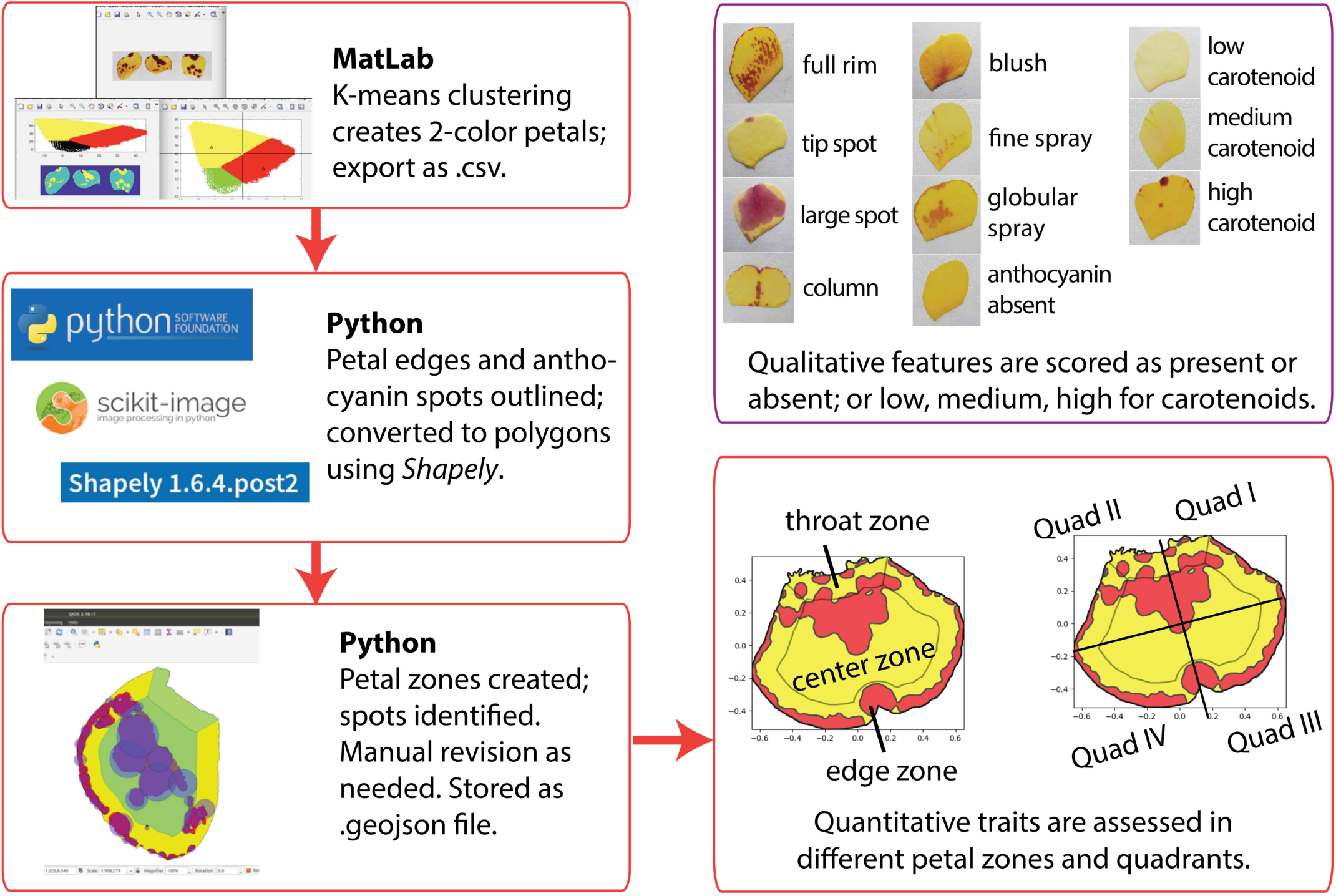
Digital image analysis workflow. See Tables 2 and 3 for the full list of qualitative and quantitative traits.

Further image analysis was undertaken using a custom pipeline (Fig. 2) implemented within the SciPy Ecosystem (Virtanen *et al*. 2020), with tools from the Scikit Image (Van der Walt *et al*. 2014) and Shapely (Gillies *et al*. 2007–) packages. In brief, spots and petals were vectorized to polygons from rasters using the Marching Squares (Cubes) algorithm (Lorensen and Cline 1987); holes in spots were detected with a custom algorithm; and all polygons were checked for validity and repaired if possible. Repairs to polygons were implemented by both automated methods based on Shapely validation tools, and a series of manual correction programs based on the Matplotlib pyplot framework. These manual curations also acted as a final quality control step for vector images. All petal polygons were assigned a unitless area of one, and spot sizes were scaled relative to this total area, to allow generalized comparisons among flower petals and images of different sizes. Several programs for hand-curation of spot and petal vectors were created and used to ensure final image quality.

**Table 3.** Quantitative traits utilized for genetic mapping. Each trait was measured in three petals per flower (two upper – averaged together to yield a single value – and one lower).

| Trait category | Trait name | Trait description |
| --- | --- | --- |
| Spot characteristics | SpotBiggest | Area of the largest detected spot |
|  | SpotSmallest | Area of the smallest detected spot |
|  | SpotMean | Mean area of all spots |
|  | SpotMedian | Median area of all spots |
|  | SpotNumber | Total number of spots on petal |
|  | PropRed | Proportion of petal covered in red pigment |
| Center zone | CenterNumWithin | Number of spots contained entirely in the center zone |
|  | CenterNumTouch | Number of spots touching the center zone |
|  | CenterNumMostly | Number of spots with a majority of their area in the center zone |
|  | CenterNumCentroids | Number of spots whose center of mass is located in the center zone |
|  | CenterProp1 | Proportion of total spot area in center zone |
|  | CenterProp2 | Proportion of center zone covered by spots |
|  | CenterDist1 | Mean distance between spot centers of mass and the center of all spots |
|  | CenterDist2 | Mean distance between spot centers of mass and the center of only the spots that are located in the center zone |
| Edge zone | EdgeNumWithin | Number of spots entirely contained in edge zone |
|  | EdgeNumTouch | Number of spots touching edge zone |
|  | EdgeNumTouch2 | Number of spots touching actual petal edge (not the edge zone) |
|  | EdgeNumMostly | Number of spots with a majority of their area in the edge zone |
|  | EdgeProp1 | Proportion of total spot area in edge |
|  | EdgeProp2 | Proportion of edge zone covered by spots |
|  | EdgeProp3 | Proportion of length of petal edge (not edge zone) covered in spots |
|  | EdgeDist1 | Mean distance between the petal rim and the closest edge of each spot |
|  | EdgeDist2 | Mean distance between the petal rim and each spot's center of mass |
| Throat | ThroatNumTouch | Number of spots touching throat zone |
|  | ThroatNumTouch2 | Number of spots touching the petal base |
|  | ThroatNumMostly | Number of spots mostly in throat zone |
|  | ThroatProp1 | Proportion of total spot area in throat |
|  | ThroatProp2 | Proportion of throat zone covered by spots |
| Petal regions: "X" indicates that each trait was measured for proximal and distal petal halves, and for Quadrants 1–4 | X Num | Number of spots with centers of mass in Region X |
|  | X Prop1 | Proportion of total spot area in Region X |
|  | X Prop2 | Proportion of Region X covered by spots |

The final set of vectors representing the petal, spots, and holes-within-spots was saved by the pipeline as GeoJSON files that are readable with standard GIS software. Following the conversion of photos to vectors, spots were counted and measured in relation to the size of their respective petals and their position within the petals. These measurements were then used as quantitative phenotypic traits in the QTL experiment.

Code for the Python image processing package is available on GitHub at: github.com/danchurch/mimulusSpeckling. It is also available at pypi.org: pypi.org/project/makeFlowerPolygons-dcthom/.

A total of 104 petal traits were computationally measured to assess the features listed in Table 3 for both upper petals and lower petals (52 traits each) for genetic mapping. A total of 353 plants were assessed, of which 310 were ultimately used for genetic mapping.

### Carotenoid extraction and analysis

Carotenoids were extracted from ventral petals of yellow-flowered *M. cupreus* and *M. cupreus* x *M. l. variegatus* F1 hybrids, as well as from a series of four F2 hybrids that display pigmentation ranging from pale to dark yellow. The petal area used for each extraction was standardized by closing a 1.5 mL Eppendorf tube over the petals to create a “punch”. Two punches per flower were placed in 200 µL of methanol and ground with a nylon pestle until the tissue became colorless.

The ground tissue was centrifuged at 16,000 g for 1 min to pellet debris, after which the supernatant was transferred to a clean Eppendorf tube. Carotenoids and flavonoids were then partitioned by adding 150 µL of methylene chloride and 150 µL of distilled water to each tube. The carotenoid-containing layer (i.e., lower phase) of selected samples was collected and dried under a gentle stream of nitrogen gas. The dried extracts were treated with ethanolic potassium hydroxide to remove fatty acid esters according to the methods of Schiedt (1995). Saponified carotenoids were dried under nitrogen gas, redissolved in 9:1 (v/v) hexanes/acetone and chromatographed according to the method outlined in LaFountain *et al*. (2015).

Carotenoids were identified based on their absorption spectra and relative order of elution as compared to a previously published analysis (LaFountain *et al*. 2015). The relative percentages of each carotenoid per sample were determined by integrating the area under each carotenoid peak (Waters Empower 2.0 Software), manually correcting for differences in their extinction coefficients (Britton 1995) and comparing each value to the total carotenoid content.

### QTL Mapping

Fresh leaf tissue (0.09–0.10 g per leaf) was collected from each of 353 *M. l. variegatus* RC6 x yellow *M. cupreus* LM43 F2 plants whose petals had been photographed. Tissue was snap frozen in liquid nitrogen. DNA was extracted using a Qiagen DNeasy Plant Mini Kit (Germantown, MD, USA), double eluted in 30–35 µL of warm dH20, and checked for concentration using a Qubit 4 Fluorometer (Invitrogen, Carlsbad, CA, U.S.A.). Genotyping by Sequencing (GBS) libraries were prepared following the protocol of Elshire *et al*. (2011). In brief, 100 ng of DNA was digested with the frequent-cutting enzyme ApeKI prior to the ligation of a unique forward adaptor and a standard common adaptor. Up to 95 samples and a water control were subsequently pooled prior to PCR amplification.

Three lanes of Illumina HiSeq 100 bp single-end sequencing were performed by the Duke University Center for Genomic and Computational Biology. Sequences were demultiplexed using Stacks version 2.1 (Catchen *et al*. 2011) and aligned to the *M. l. luteus* genome using bowtie2 and SNPs were called using GATK HaplotypeCaller. The resultant VCF file was filtered to remove sites with greater than 30% missing data and a minor allele frequency cutoff of 0.05. The file was then converted into a HapMap using Tassel ‘`run_pipeline.pl`’ to convert HapMap to CSV. The `run_pipeline.pl` script filters the SNP dataset to include locations that meet the following cutoffs: (1) the parents were genotyped and (2) the parental genotype is not heterozygous (https://bitbucket.org/tasseladmin/tassel-5-source/wiki/UserManual/GenosToABH/GenosToABHPlugin). A total of 4616 sites survived this filtering.

The resultant CSV file was used in R/qtl to create linkage groups and conduct QTL mapping (Broman *et al*. 2003). To create linkage groups, the data were further filtered to remove markers with more than 70 missing individuals and individuals with more than 500 missing marker calls. Markers with distorted segregation patterns (as determined by cutoff of p<0.01) were filtered and duplicate individuals, defined as those who shared greater than 80% identity at markers, were removed from the dataset. Using this filtered dataset, we estimated recombination fractions between alleles, formed linkage groups, and reordered markers using the newly formed linkage map. Linkage groups were formed using a maximum recombination fraction of 0.25, minimum LOD score of six, and markers were allowed to be reorganized to form new linkage groups. The physical *M. l. luteus* genome was not used to constrain the linkage group assembly. The median length of individual linkage groups was 102 cM with a total linkage map of 4926 cM. The *Mimulus guttatus* (diploid relative) linkage maps are 2011–2096 cM (Fishman *et al*. 2001).

We used R/qtl to perform QTL mapping for the carotenoid and anthocyanin traits shown in Tables 2 and 3. QTLs were identified using the R/qtl ‘scanone’ function, with the ‘em’ method and 5000 permutations. A ‘binary’ model was used for the manually scored, qualitative phenotypes while a ‘normal’ model was used for the computationally measured, quantitative phenotypes. For QTL analysis of quantitative traits, individuals that lacked anthocyanin on their petal lobes were excluded. Quantitative-trait analyses were performed separately on the upper and lower petals.

To generate the graphs shown in Figures 3–5, trait means were calculated for each genotype at the highest LOD score marker for that trait. Genotypes were compared in RStudio version 2026.01.0+392 using a General Linear Model with either a binomial distribution (for qualitative traits with two levels) or a normal distribution. Tukey post hoc tests were performed if significant differences among genotypes were reported.

**Figure 3.**
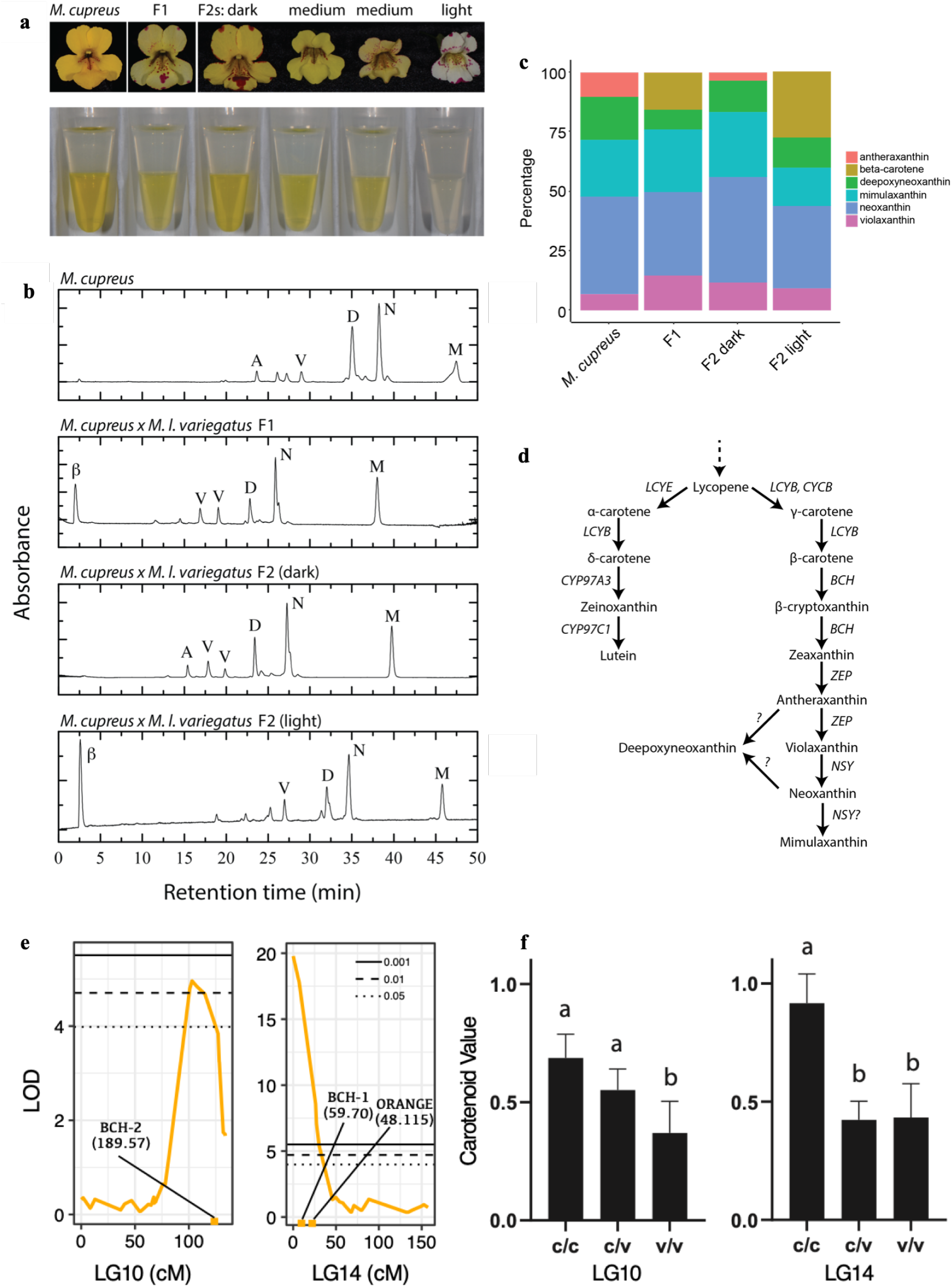
Intensity of yellow pigmentation corresponds with shifts in carotenoid composition. (a) The *M. cupreus* parent, *M. cupreus* x *M. l. variegatus* F1 hybrid, a dark F2 individual (third from left), and a pale-yellow individual (fifth from left) were used for liquid phase partitioning of flavonoids and carotenoids. (b) HPLC chromatograms of floral extracts. Abbreviations are as follows: A, antheraxanthin; B, beta-carotene; D, deepoxyneoxanthin; M, mimulaxanthin; N, neoxanthin; V, violaxanthin. (c) Relative percentages of carotenoids as determined by integration of HPLC peak areas. (d) A partial carotenoid pathway based on LaFountain *et al*. (2015) and Stanley and Yuan (2019). (e) LOD scores and candidate genes for QTL-10 and QTL-14. (f) Mean trait values with 95% confidence intervals across *cupreus* (c/c), heterozygous (c/v), and *variegatus* (v/v) genotypes. Letters show significance groupings based on Tukey post hoc tests.

### Identification of candidate genes

Following QTL mapping, we extracted the markers under each significant QTL peak. For QTL peaks where multiple traits were mapped, a single representative trait was used. The specific location for traits can be found in Supplemental File 2 (TraitLocations.txt). We then extended the QTL window 100k bases (based on bordering markers) in each direction and extracted all genes within that window. The protein sequences for these genes were searched against the Arabidopsis protein database using blastp. Next, using ChatGPT to search the BLAST output, we identified genes potentially related to anthocyanin biosynthesis, pattern formation or carotenoid biosynthesis. The raw BLASTP outputs, ChatGPT prompts, and filtered files are available in Supplemental File 2. We also extracted all putative MYBs within broader 1Mb of the anthocyanin peak on QTL-9b. Next, these Mybs were added to the MYB alignment from Yuan et al. (2014) and aligned them using Mafft. The alignment was trimmed, removing sites with greater than 70% missing data, and a tree was built using iqtree3.

### Recombinant Inbred Lines

Recombinant inbred lines (RILs) were developed from parental lines originally collected in Chile in 2004 (Table 1). *Mimulus luteus* var. *variegatus* and a yellow-flowered morph of *Mimulus cupreus* were self-pollinated for several generations after collection to establish homozygosity before use in crosses. The recombinant inbred line (RIL) breeding scheme began with a cross between the two parental lines to produce heterozygous F1 hybrids. These F1 plants were selfed to generate a diverse F2 population. To increase genome-wide recombination, F2 individuals were randomly intercrossed, resulting in a genetically mixed F3 generation. F3 individuals were designated as the first generation of each RIL and assigned unique identifiers. From the F3 generation onward, single-seed descent was carried out for at least five generations. At each generation, petal pigmentation pattern phenotypes were photographed and recorded. To investigate whether pattern subtypes are genetically separable, we selected a subset of 5th- and 6th-generation RIL lines with distinct phenotypes and performed targeted crosses. The individuals selected for crossing exhibited one of the following phenotypes: speckles, tip spots, blush, solid pigmented, or blotchy spots. Resulting F1s were selfed to produce F2 populations. F2 individuals were grown under standard conditions and photographed at flowering.

### Microscopy of the “blush” trait

Early-generation RIL plants producing petals with the blush phenotype were imaged using a HIROX KH-7700, a high-resolution digital microscopy system. Fresh petals were collected from plants and placed whole, without fixation or preparation, under the HIROX for imaging.

### Data availability

Supplementary File 1 contains a summary table of all QTLs and the traits that mapped to them, along with graphs for each statistically significant QTL. Supplementary File 2 lists the output from BLAST results and candidate gene searches within the QTLs. Supplementary File 3 contains a larger version of the MYB gene tree shown in Figure 4. Supplementary File 4 shows a variety of crosses between Recombinant Inbred Line (RIL) parents that differ for blush and spot traits. Supplementary File 5 shows genotypes of F2s with the most “variegatus-like” petal phenotypes (high proportion anthocyanin-pigmented). Sequence data will become available on GenBank upon publication. The Python image processing package is available on GitHub at <github.com/danchurch/mimulusSpeckling>. Raw sequencing data (fastq files) used in QTL mapping will be available in SRA upon publication. Seeds of inbred lines are available upon request.

**Figure 4.**
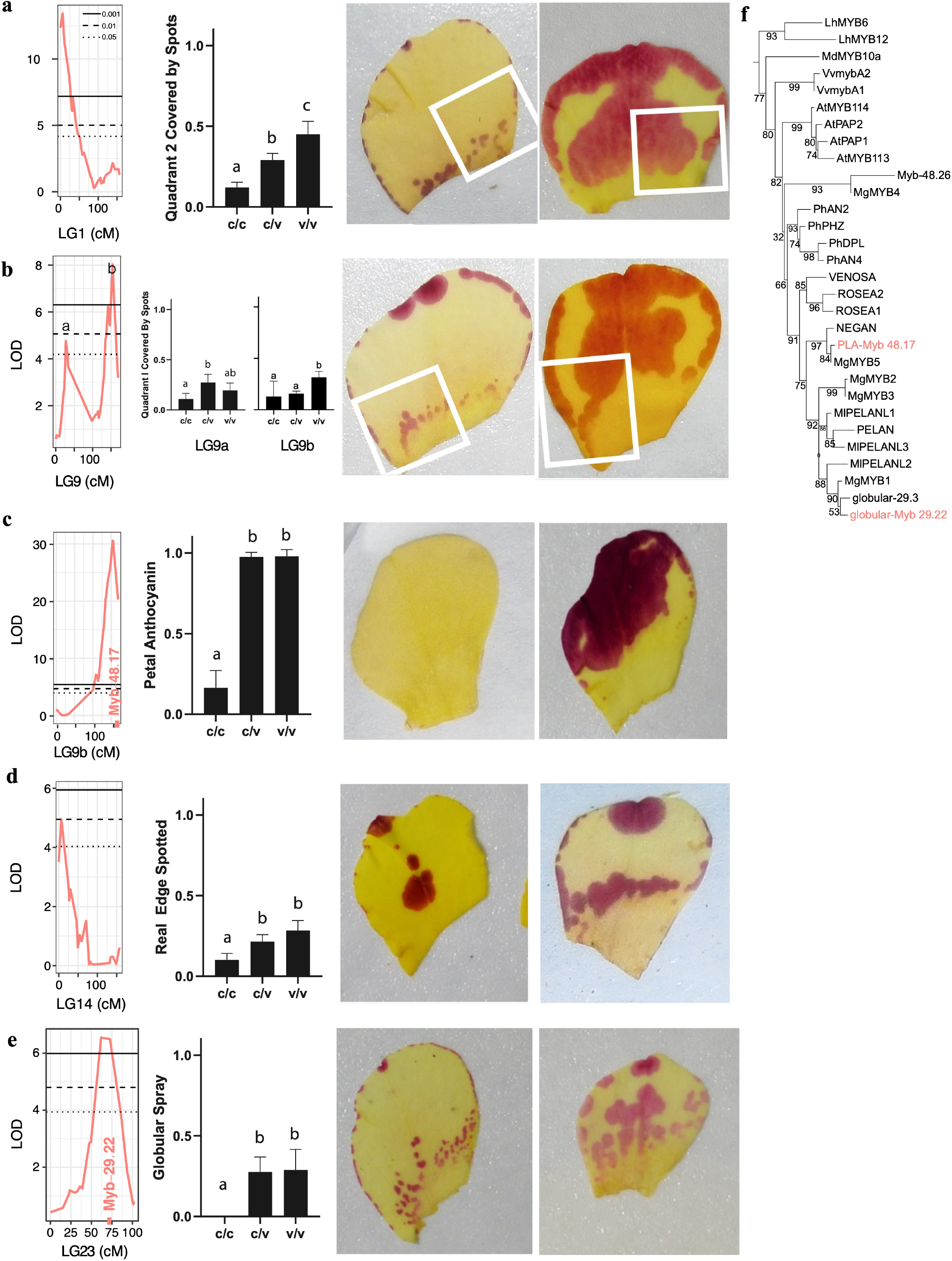
Quantitative anthocyanin traits mapped to five QTLs. Sample traits shown are: (a) quadrant 2 covered by spots; (b) Quadrant 1 covered by spots; (c) presence vs absence of Petal Lobe Anthocyanin; (d) real edge spotted; (e) presence versus absence of globular spray. Within each row, the first column shows LOD scores for the QTL(s) for the sample trait. The second column shows mean trait values by genotype (c=*cupreus* and v=*variegatus*), with 95% confidence interval error bars and significance groupings from Tukey post hoc tests indicated above the columns. The third and fourth columns show examples of a petal scored as having a low (left) versus high (right) value for that trait. QTL graphs for all other quantitative traits can be found in Supplemental File S111.(f) A maximum likelihood tree of R2R3 MYBs shows MYBs under QTL fall within NEGAN and PELAN MYB phylogenetic groups.

## Results and Discussion

### Chromosome-level assembly of the M. l. luteus reference genome

The newly assembled *Mimulus luteus* var. *luteus* genome size of 599 Mb, with a scaffold L50 of 32 and N50 of 6.4 Mb, compares favorably with estimates for the genome size of 640–680 Mb made by flow cytometry and a previous assembly of 410 Mb that had a scaffold N50 of 0.28 Mb (Edger *et al*. 2017). Notably, this revised assembly appears to incorporate both more duplicated polyploid gene space and pericentromeric sequence space missing from the previous *luteus* assembly and the assembly of the diploid relative *M. guttatus* (JGI version 5.0, Phytozome-13: MguttatusTOL551v5.0) respectively.

Consistent with a more complete polyploid gene space, the maker annotation gene count increased to 53,411 genes from 46,855 in the previous assembly and was nearly twice the *M. guttatus* diploid protein-coding gene count (2×26,718(guttatus): 53,436). The similarity of these two values may be somewhat coincidental since the short-read *M. guttatus* assembly was likely depleted in gene space relative to the long-read *M. l. luteus* assembly, whilst diploidization will have begun to reduce the gene count in *M. l. luteus*, bringing these two values together. Missing BUSCO genes remained around 2.5%, similar to the level in the previous assembly (2.6%). The *luteus* assembly contained significant blocks of sequence without evident CDS homology in the assembled *M. guttatus* genome, although similar sequences can be found in unassembled *M. guttatus* BAC and clone sequence available in GenBank. The size of these gene-poor blocks missing from the *M. guttatus* assembly ranged up to c. 10 Mb, and was consistent with the incorporation of more complete centromeric and pericentromeric sequence in the long-read *M. l. luteus* assembly. However, tandem duplicate arrays make scaffolding of these regions with HiC reads more prone to error than HiC scaffolding of chromosome arm regions.

### Carotenoid composition changes with intensity, and maps to QTLs containing beta-carotene metabolism and chromoplast-related genes

Liquid-liquid phase partitioning of flavonoids and carotenoids confirmed that the yellow pigmentation in the petals is due to carotenoids (Fig. 3). Flavonoid-based pigments (e.g., anthocyanins or chalcones) would be expected to migrate to the upper, aqueous layer but were not observed in the samples. Carotenoids are expected to migrate to the lower, methylene chloride layer. The intensity of yellow pigmentation observed in the lower layer of the samples corresponded to their respective floral colors.

To further interrogate the identity of the pigments, extracts were separated by high-performance liquid chromatography (HPLC). These data revealed that all flowers sampled synthesized the xanthophylls produced by the late carotenoid biosynthetic pathway enzymes (e.g., neoxanthin, deepoxyneoxanthin, and mimulaxanthin; Fig. 3d). However, the intermediate-yellow F1 and lightest-yellow F2 individuals showed an increased relative concentration of beta-carotene, a precursor to the xanthophylls.

Two carotenoid QTLs were found on linkage groups 10 and 14 (Fig. 3; File S1). At both QTLs, the derived light yellow state was associated with the allele from the lighter yellow *variegatus* parent. Each QTL contained a gene with best match to *Beta-Carotene Hydroxylase* (*BCH*; File S2), whose enzyme hydroxylates orange colored beta-carotene to produce the yellow colored zeaxanthin (Sun *et al*. 2022). QTL-10 also contained a gene identified as *ORANGE*, which promotes carotenoid biosynthesis as well as the formation of carotenoid-storing chromoplasts (Wrightstone *et al*. 2025).

Two other putatively carotenoid-related genes (File S2) were *CYP97A3/LUT5* at QTL-10, encoding a hydroxylase that helps convert alpha-carotene to lutein (Kim and DellaPenna 2006), and *Violaxanthin De-Epoxidase 1* (*VDE1*) at QTL-14, whose product converts orange colored violaxanthin to yellow-colored and more photoprotective zeaxanthin when light intensity is high (Bugos *et al*. 1998). Because neither parent species shows an accumulation of alpha-carotene, and because the yellow colors appear to be fixed rather than dependent on light intensity, we consider these candidate genes unlikely to contribute to the carotenoid intensity difference between *M. l. variegatus* and *M. cupreus*.

The carotenes in the upstream portion of the biosynthetic pathway are oxygenated to produce downstream xanthophylls (Fig. 3d). *ORANGE* promotes the development of carotenoid-storing chromoplasts, some of which almost exclusively store esterified xanthophylls rather than carotenes (Vishnevetsky *et al*. 1999). BCH catalyzes the transition from carotene to xanthophyll, and disruption of the *BCH* gene in tomato results in white flowers (despite the accumulation of orange-hued beta carotene), presumably due to the petals’ inability to store these upstream carotene pigments (Galpaz *et al*. 2006). Subsequent work has confirmed that xanthophyll esters are essential to both chromoplast development and the accumulation of yellow coloration, such that a reduction in xanthophyll esters leads to a major reduction in total carotenoid content (Ariizumi *et al*. 2014). Conversely, *BCH* overexpression has been shown to increase metabolic flux into the carotenoid biosynthetic pathway, resulting in higher total carotenoid accumulation (Zhou *et al*. 2011). Genes *ORANGE* and *BCH*, therefore, are both excellent candidates for the derived loss of yellow pigmentation in *M. l. variegatus*.

We hypothesize that a reduction in the function of *BCH* explains the relative accumulation of beta carotene in *M. l. variegatus*, while also contributing to lower total carotenoid content. The accumulated beta carotene may be less able to be stored in petal chromoplasts compared to the downstream xanthophylls, leading to its degradation. Finally, a disruption in *ORANGE* is expected to reduce both the activity of the carotenoid biosynthetic pathway and the biogenesis of the chromoplasts that are needed to store the resulting carotenoids. Functional analyses of the *BCH* and *ORANGE* candidate genes are needed to evaluate these hypotheses.

### The Petal Lobe Anthocyanin (PLA) QTL-9b recovers the anthocyanin-activating MYB5a/NEGAN transcription factor gene, which acts as both a switch and a patterner

In flowering plants, changes in color patterning have frequently been tracked to the anthocyanin-activating Subgroup 6 *R2R3 MYB* genes (Schwinn *et al*. 2006; Lowry *et al*. 2012; Yamagishi 2013; Streisfeld *et al*. 2013; Yuan *et al*. 2014; Twyford *et al*. 2018). The evolutionarily recent and genetically dominant gains of petal lobe anthocyanin pigmentation, in both the common orange morph of *M. cupreus* and the magenta-flowered *M. l. variegatus*, follow this pattern. Pigment gain has been mapped in each case to a single genomic region (*pla1* and *pla2*, respectively). Each genomic region spans a tandem array of Subgroup 6 *R2R3 MYB* genes, and the “activating” allele at each region is associated with higher expression of the late biosynthetic genes that are the targets of the R2R3 MYB (Cooley *et al*. 2011). In *M. l. variegatus*, the causal gene has been pinpointed as *MYB5a/NEGAN* (Zheng *et al*. 2021).

Consistent with the previously documented Mendelian nature of PLA, we observed a 3:1 ratio of PLA presence versus absence in our mapping population, with 228 out of 310 (73.5%) of the F2 plants showing the dominant *variegatus* phenotype of anthocyanin production in the petal lobes. *MYB5a/NEGAN* is located within the largest QTL, on linkage group 9 (QTL-9b in Fig. 4c; File S3). An analysis on just the subset of plants containing PLA also identified 14 quantitative patterning traits mapping to QTL-9b (File S1), showing that PLA is not only “switched on” by this genomic region, but patterned by it as well.

### PLA, when present, is patterned by five QTLs

A total of 39 quantitative traits (Table 3) mapped to five QTLs located on linkage groups 1, 9, 14, and 23 (Fig. 4). Many traits mapped to QTL-1, -9b, and -23, while only a single trait each mapped to QTL-9a and -14 (File S1). For approximately half of the mapped traits, the upper petals and the lower petal both mapped to the same QTL, suggesting a partly shared genetic program between these two floral regions. Two whole-flower qualitative traits were also mapped: globular spray to QTL-23, and blush to QTL-9b and QTL-23. Across the five anthocyanin QTLs, the *variegatus* allele was generally associated with more and larger spots (Fig. 4).

An AI-based assessment of candidate genes under the QTLs highlighted *MYB* genes both in the anthocyanin-activating clade and in an adjacent, flavonoid-regulating and floral development clade (File S2). Other possible candidates included genes in the anthocyanin biosynthetic pathway; *bHLH* genes contributing to the MYB-bHLH-WDR (MBW) transcriptional activation complex; and a handful of genes with functions that could potentially impact color or patterning, associated with traits such as vacuolar pH and pigment transport. A particularly compelling candidate was the *pla1* locus (Cooley *et al*. 2011), containing *MYB1, MYB2*, and *MYB3*, orthologous to the anthocyanin-activating gene *PELAN* and located within QTL-23.

### Blush is activated by a variegatus allele at QTL-23, and can be obscured by QTL-9b

At QTL-23, plants showed an absolute requirement for at least one *variegatus* allele in order to exhibit a thin, apparently finely-speckled layer of faint anthocyanin pigment at the petal base that we dubbed “blush”. Contrary to our expectations, QTL-9b showed the opposite pattern: blush was associated with the allele from the yellow morph of *cupreus*, which lacks petal lobe anthocyanins entirely. We noticed that plants scored as having blush typically lacked the darker red spots typically associated with the QTL-9b *variegatus* allele.

One explanation could be that this reflected the limitations of phenotyping, with the presence of red petal spots (a QTL-9b trait) tending to obscure the much fainter blush (a QTL-23 trait) when both are present. An alternative hypothesis is that the presence of spots somehow re-patterns or eliminates blush. Collectively, these results indicate that *M. l. variegatus* can activate a faint layer of anthocyanin pigmentation using a locus that is entirely separate from the *MYB5a/NEGAN* gene at QTL-9b that generates intense full-petal pigmentation in *M. l. variegatus* and spots in hybrids.

### Blush is an unpatterned phenotype, yet creates a speckling effect due to subcellular localization and can lead to spot formation in crosses

Microscopy of petal tissue exhibiting blush revealed that it is an unpatterned phenotype (Fig. 5). Every cell in the blush region of each petal contained anthocyanin pigment. The impression of fine speckling that blush gives seems to be created by the fact that anthocyanins are sequestered into the vacuoles of the cells (Grünig *et al*. 2026), with unpigmented cytoplasmic space outside of the vacuoles. Crosses between RILs characterized by strong versus weak blush yielded F1 hybrids with intermediate blush intensity (File S4). The F2s showed a range of blush intensities, evocative of a classic, unpatterned, quantitative trait.

**Figure 5.**
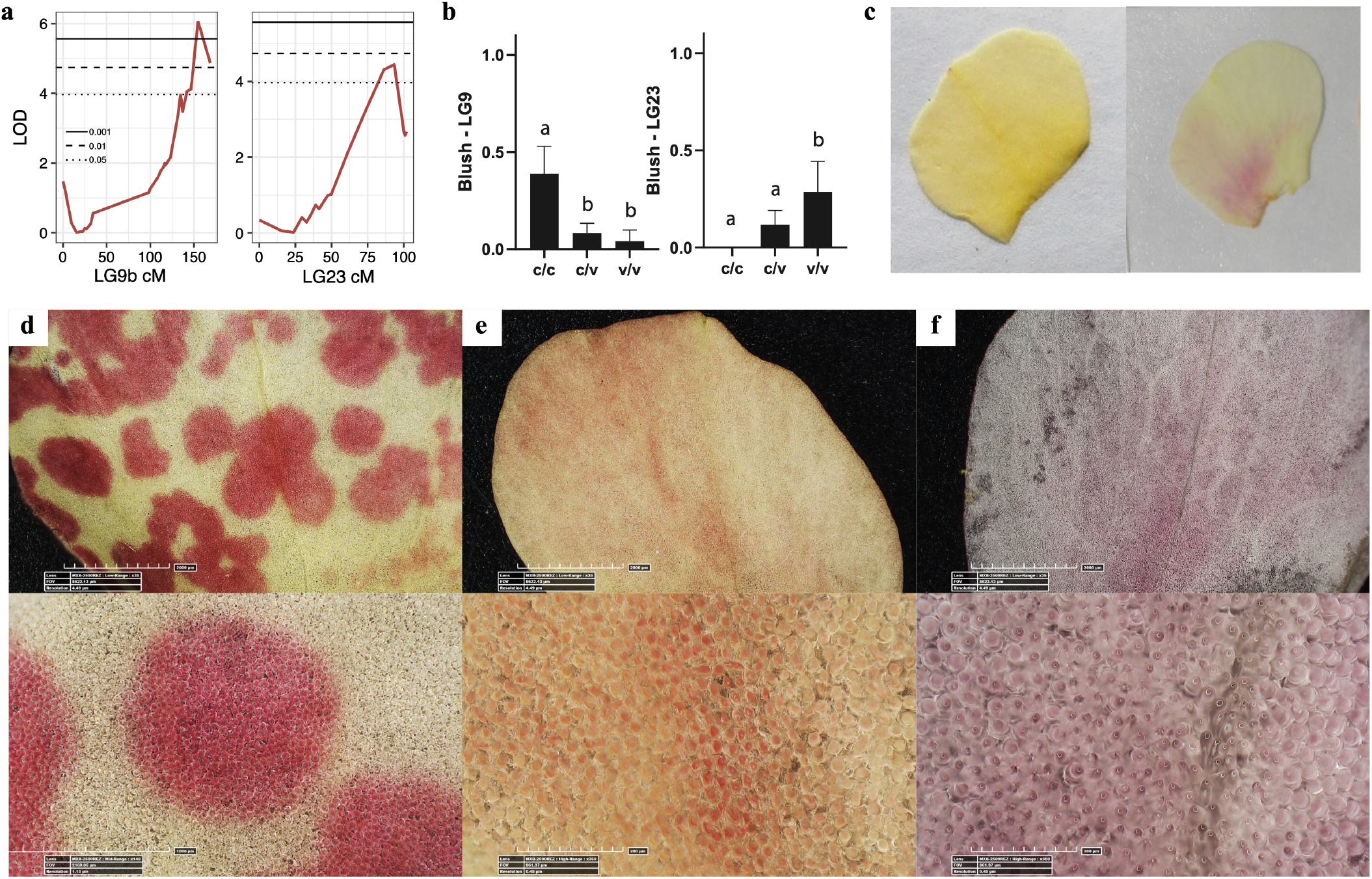
Blush maps to two QTLs with opposite effects. (a) At QTL-9b and QTL-23, blush is associated with (b) the *cupreus* (non-anthocyanin parent) and *variegatus* (anthocyanin parent) alleles, respectively. (c) The blush phenotype is quite faint, as seen in a comparison of petals without blush (left) versus with blush (right). Microscopy of RIL lines 149 (d), 111 (e), and 295 (f) indicates that spots (d) and blush (e, f) both represent continuous, unpatterned anthocyanin production at the cellular level. Each cell within the pigmented region produces anthocyanin, though the amount of pigment in blush regions is so small that the unpigmented cytoplasmic space around each vacuole can create an impression of fine-scale speckling.

A regulatory network capable of Turing pattern formation can also exhibit spatially uniform steady states under some parameter regimes, as modeled in Simmons *et al*. (2023). A simpler explanation is that candidate genes *MYB1, MYB2*, and/or *MYB3* are non-Turing (non-patterning) anthocyanin activators. In support of this hypothesis, their ortholog in *Mimulus lewisii* – the gene *PELAN* – is a non-self-activating *MYB* gene associated with solid, unpatterned petal anthocyanin pigmentation (Yuan *et al*. 2014).

An intriguing example of the interplay between blush and spot patterning was seen in the RIL cross labeled “Gigi” (Fig. 6a). One RIL parent had blush in the petal interior and no spots. The other parent had small spots along the petal edges and tips, but no interior anthocyanin. The F1 hybrid did have anthocyanin in the petal lobe interior, like the blush parent… but patterned into spots, rather than blush. In the F2 generation, most plants had interior spots, while a few had blush. Crosses made with other spotted RIL parents confirmed these results: F1s of blush x spot parents showed spots but not blush, and the blush trait reappeared in a small fraction of the F2s, while spots were present in the majority (File S4).

**Figure 6.**
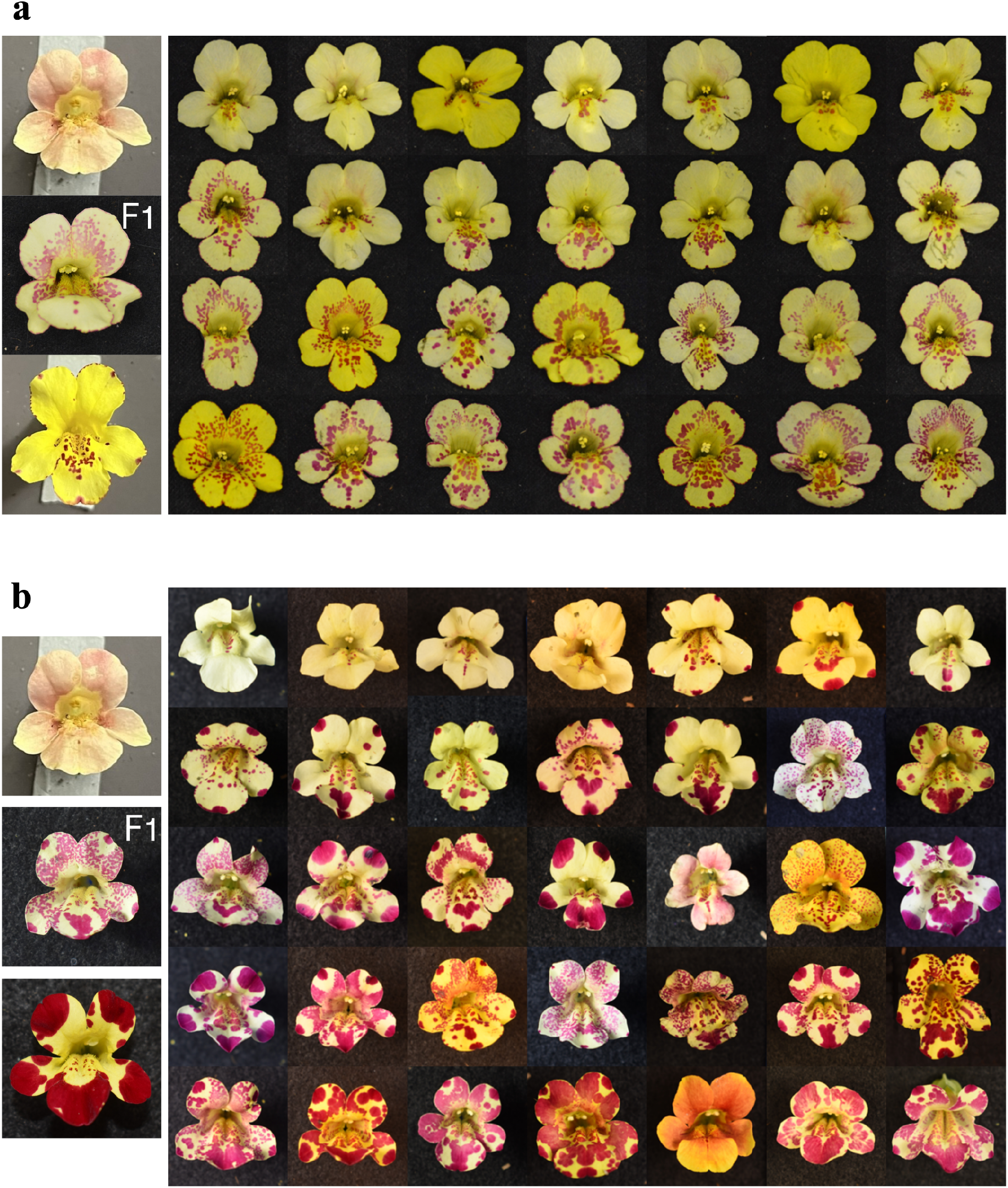
RIL crosses illustrate the interaction of spots (activated by QTL-9b) and blush (activated by QTL-23). (a) The “Gigi” cross: Left column shows RIL 111.5 (top), F1 (middle), and RIL 270.5 (bottom). Photo panel on right contains one flower per plant from 28 F2s. (b) The “Peggy” cross: Left column shows RIL 111.5 (top), F1 (middle), and RIL 292.5 (bottom). Photo panel on right contains one flower per plant from 35 F2s.

How might a solid trait like blush be repatterned into interior-petal spots, when parental spots were found only on the petal edge? One possibility is that a gene associated with the blush trait, and expressed in the petal interior, could activate the *variegatus* allele of *MYB5a/NEGAN*. That would lead to the production of spotted Petal Lobe Anthocyanin (PLA) in the interior-petal space. Because PLA overrides blush, the blush trait would only be produced in a plant homozygous for the non-PLA *cupreus* allele of *MYB5a/NEGAN* (Table 4).

**Table 4.** Epistasis model for spots versus blush. Hypothesized genotype-phenotype relationships are shown for QTL-9b and QTL-23. v, *variegatus* allele; c, *cupreus* allele

| QTL-9b (PLA) | QTL-23 (Blush) | Petal lobe trait |
| --- | --- | --- |
| v/_ | v/_ | spots |
| v/_ | c/c | spots |
| c/c | v/_ | blush |
| c/c | c/c | no anthocyanin |

Potential direct or indirect activators of *MYB5a/NEGAN* include transcription factors belonging to the bZIP, WRKY, NAC, ERF/AP2, and HD-ZIP gene families (Yang *et al*. 2022). Alternatively, *MYB* genes 1, 2, or 3 might themselves activate *MYB5a/NEGAN*, given that *MYB5a/NEGAN* is self-activating (Ding *et al*. 2020) and that *MYB1-2-3* encode proteins quite similar to MYB5a/NEGAN (Cooley *et al*. 2011).

### Tip spots show properties of both positional specification and Turing systems

Spots at the petal tips, seen in many of the F2s of the “Peggy” cross (Fig. 6b), provide striking visual illustrations of the mechanism of a Turing-style reaction-diffusion spot. Each spot, putatively caused by a random initial activation of *MYB5a/NEGAN* at its center, is surrounded by a clearly unpigmented zone of inhibition as expected from the outward diffusion of the (unknown) spot inhibitor (Turing 1952). Yet, in direct contrast to Turing’s predictions, tip spots are not randomly located. Rather, their location is highly predictable: they are found at the distal tip of the midvein of each petal. This trait highlights how two classic pattern-determining systems – Turing’s semi-stochastic repetitions versus the positional specification model proposed by Wolpert (1969) – are not mutually exclusive (Brückner 2026).

## Conclusions

Interspecies hybrids expand the range of phenotypic variation seen in nature, presenting opportunities to investigate traits that are not typically found in existing species. Here, we investigate the genetic architecture of a classic, spatially simple interspecies difference in yellow carotenoid pigmentation as well as that of a spatially complex anthocyanin patterning phenotype that emerges only in interspecies hybrids. We identify the carotenoid biosynthesis enzyme *BCH* and chromoplast and carotenoid-pathway regulator *ORANGE* as top candidates for the evolutionarily recent loss of yellow intensity in *M. l. variegatus*. The genetic architecture of hybrid-specific anthocyanin patterning is more complex, with at least five QTLs contributing to pattern variation. Pattern QTLs include two anthocyanin-activating loci as top candidates: the Turing system patterning gene *MYB5a/NEGAN* at QTL-9b, and the putatively non-patterning *MYB1-2-3/PELAN* at QTL-23. *variegatus* alleles of at least three loci appear to be required to approximate the fully-anthocyanin-pigmented phenotype of *M. l. variegatus* petals. Trait segregation in RIL and F2 crosses suggests the involvement of positional specification as well as Turing systems in the determination of anthocyanin spot placement. Collectively, our results begin to reveal the complex genetics behind an apparently simple flower color switch from yellow to magenta in *M. l. variegatus*.

## Acknowledgements

We thank Taylor Wilke for earlier work that helped pave the way for ideas explored in this paper. Thanks to John Kelly, Lila Fishman, and anonymous reviewers for insightful feedback on the manuscript. Funding was provided by NSF-DEB 1655311 (to AMC), NSF-DEB 1754075 (to AMC and JRP), NSF-IOS 2031272 (to AMC and JRP), Whitman College, and William & Mary.

