## Supplemental File 1 for "Genetic architectures of floral pigment and patterning in hybrid monkeyflowers"

| QTL | Trait | Upper-petal | Lower-petal |
|---|---|---|---|
| 1 | SpotBiggest | x | x |
| | CenterProp1 | x | |
| | PropRed | x | x |
| | CenterProp2 | x | x |
| | CenterDist1 | | x |
| | CenterDist2 | | x |
| | EdgeProp2 | x | x |
| | EdgeProp3 | x | |
| | EdgeDist1 | x | |
| | ThroatProp1 | x | x |
| | ThroatProp2 | x | |
| | DistalProp2 | x | x |
| | ProximalProp2 | x | x |
| | Quad1Prop2 | x | x |
| | Quad2Prop2 | x | x |
| | Quad3Prop2 | x | x |
| | Quad4Prop2 | x | x |
| 9a | Quad1Prop2 | x | |
| 9b | SpotBiggest | x | |
| | PropRed | x | x |
| | CenterProp2 | x | x |
| | CenterDist1 | x | |
| | EdgeProp2 | x | x |
| | EdgeProp3 | x | x |
| | EdgeDist2 | x | |
| | ThroatProp2 | x | |
| | DistalProp2 | x | x |
| | ProximalProp2 | x | x |
| | Quad1Prop2 | x | |
| | Quad2Prop2 | x | x |
| | Quad3Prop2 | x | x |
| | Quad4Prop2 | x | x |
| | Blush | (whole-flower trait) | |
| 14 | EdgeProp3 | x | x |
| 23 | SpotNumber | x | x |
| | CenterNumTouch | | x |
| | CenterNumMostly | | x |
| | CenterNumCentroids | | x |
| | CenterProp1 | x | x |
| | CenterProp2 | | x |
| | CenterDist2 | | x |
| | EdgeNumWithin | x | x |
| | EdgeNumTouch | x | x |
| | EdgeNumTouch2 | x | x |
| | EdgeNumMostly | x | x |
| | EdgeProp1 | x | x |
| | EdgeProp3 | | x |
| | EdgeDist1 | | x |
| | EdgeDist2 | | x |
| | DistalNum | x | x |
| | DistalProp1 | x | x |
| | ProximalNum | x | x |
| | ProximalProp1 | x | x |
| | ProximalProp2 | | x |
| | Quad1Num | | x |
| | Quad1Prop1 | x | x |
| | Quad1Prop2 | | x |
| | Quad2Num | x | |
| | Quad2Prop1 | x | x |
| | Quad2Prop2 | x | x |
| | Quad3Num | | x |

| | | | |
|---|---|---|---|
| | Quad3Prop1 | x | |
| | Quad4Num | x | |
| | Quad4Prop1 | | x |
| | Globular spray | (whole-flower trait) | |
| | Blush | (whole-flower trait) | |

**QTLs for quantitative anthocyanin traits:**

| Trait category | Trait name | QTL-1 | QTL-9a | QTL-9b | QTL-14 | QTL-23 | Full trait description followed by label used on the QTL graph |
|---|---|---|---|---|---|---|---|
| Spot characteristics | SpotBiggest | upper | | upper | | | Area of the largest detected spot |
| | | lower | | | | | *biggestSpotArea* |
| | SpotSmallest | | | | | | Area of the smallest detected spot |
| | | | | | | | *smallestSpotArea [no QTLs for this trait]* |
| | SpotMean | | | | | | Mean area of all spots |
| | | | | | | | *AvSpotSize [no QTLs for this trait]* |
| | SpotMedian | | | | | | Median area of all spots |
| | | | | | | | *MedSpotSize [no QTLs]* |
| | SpotNumber | | | | | upper | Total number of spots on petal |
| | | | | | | lower | *NuSpots* |
| | PropRed | upper | | upper | | | Proportion of petal covered in red pigment |
| | | lower | | lower | | | *PropRed* |
| Center | CenterNumWithin | | | | | | Number of spots contained entirely in the center zone |
| | | | | | | | *NuSpotsContainedInCenter [no QTLs]* |
| | CenterNumTouch | | | | | | Number of spots touching the center zone |
| | | | | | | lower | *NuSpotsTouchCenter* |
| | CenterNumMostly | | | | | | Number of spots with a majority of their area in the center zone |
| | | | | | | lower | *NuSpotsMostlyInCenter* |
| | CenterNumCentroids | | | | | | Number of spots whose center of mass is located in the center zone |
| | | | | | | lower | *NuSpotCentroidsInCenter* |
| | CenterProp1 | upper | | | | upper | Proportion of total spot area in center zone |
| | | | | | | lower | *PropSpotsInCenter* |
| | CenterProp2 | upper | | upper | | | Proportion of center zone covered by spots |
| | | lower | | lower | | lower | *CenterCoveredBySpots* |
| | CenterDist1 | | | upper | | | Mean distance between spot centers of mass and the center of all spots |
| | | lower | | | | | *AvgDist2CtrAllSpots* |
| | CenterDist2 | | | | | | Mean distance between spot centers of mass and the center of only the spots that are lo |
| | | lower | | | | lower | *AvgDist2CtrCtrSpots* |
| Edge | EdgeNumWithin | | | | | upper | Number of spots entirely contained in edge zone |
| | | | | | | lower | *NuSpotsContainedInEdge* |
| | EdgeNumTouch | | | | | upper | Number of spots touching edge zone |
| | | | | | | lower | *NuSpotsTouchEdge* |
| | EdgeNumTouch2 | | | | | upper | Number of spots touching actual petal edge (not the edge zone) |
| | | | | | | lower | *NuSpotsTouchActualEdge* |
| | EdgeNumMostly | | | | | upper | Number of spots with a majority of their area in the edge zone |
| | | | | | | lower | *NuSpotsMostlyInEdge* |
| | EdgeProp1 | | | | | upper | Proportion of total spot area in edge |
| | | | | | | lower | *PropSpotsInEdge* |
| | EdgeProp2 | upper | | upper | | | Proportion of edge zone covered by spots |
| | | lower | | lower | | | *EdgeCoveredBySpots* |
| | EdgeProp3 | upper | | upper | upper | | Proportion of length of petal edge (not edge zone) covered in spots |
| | | | | lower | lower | lower | *RealEdgeSpotted* |
| | EdgeDist1 | upper | | | | | Mean distance between the petal rim and the closest edge of each spot |
| | | | | | | lower | *AvDistSpotEdge2Edge* |
| | EdgeDist2 | | | upper | | | Mean distance between the petal rim and each spot's center of mass |
| | | | | | | lower | *AvDistSpotCentroid2Edge* |
| Throat | ThroatNumTouch | | | | | | Number of spots touching throat zone |
| | | | | | | | *NuSpotsTouchThroat [no QTLs]* |
| | ThroatNumTouch2 | | | | | | Number of spots touching the petal base |
| | | | | | | | *NuSpotsTouchCut [no QTLs]* |
| | ThroatNumMostly | | | | | | Number of spots mostly in throat zone |
| | | | | | | | *NuSpotsMostlyInThroat [no QTLs]* |
| | ThroatProp1 | upper | | | | | Proportion of total spot area in throat |
| | | lower | | | | | *PropSpotsInThroat* |
| | ThroatProp2 | upper | | upper | | | Proportion of throat zone covered by spots |
| | | | | | | | *ThroatCoveredBySpots* |
| Distal | DistalNum | | | | | upper | Number of spots with centers of mass in distal region |
| | | | | | | lower | *NuDistSpots* |
| | DistalProp1 | | | | | upper | Proportion of total spot area in distal region |
| | | | | | | lower | *PropSpotsDist* |
| | DistalProp2 | upper | | upper | | | Proportion of distal region covered by spots |
| | | lower | | lower | | | *DistCoveredBySpots* |
| Proximal | ProximalNum | | | | | upper | Number of spots with centers of mass in proximal region |
| | | | | | | lower | *NuProxSpots* |
| | ProximalProp1 | | | | | upper | Proportion of total spot area in proximal region |
| | | | | | | lower | *PropSpotsInProx* |
| | ProximalProp2 | upper | | upper | | | Proportion of proximal region covered by spots |
| | | lower | | lower | | lower | *ProxCoveredBySpots* |
| Petal Quadrants | Quad1Num | | | | | | Number of spots with centers of mass in Quadrant 1 |
| | | | | | | lower | *NuQ1spots* |
| | Quad1Prop1 | | | | | upper | Proportion of total spot area in Quadrant 1 |
| | | | | | | lower | *PropSpotsQ1* |
| | Quad1Prop2 | upper | upper | upper | | | Proportion of Quadrant 1 covered by spots |
| | | lower | | | | lower | *Q1CoveredBySpots* |
| | Quad2Num | | | | | upper | Number of spots with centers of mass in Quadrant 2 |
| | | | | | | | *NuQ2spots* |
| | Quad2Prop1 | | | | | upper | Proportion of total spot area in Quadrant 2 |
| | | | | | | lower | *PropSpotsQ2* |
| | Quad2Prop2 | upper | | upper | | upper | Proportion of Quadrant 2 covered by spots |
| | | lower | | lower | | lower | *Q2CoveredBySpots* |
| | Quad3Num | | | | | | Number of spots with centers of mass in Quadrant 3 |
| | | | | | | lower | *NuQ3spots* |
| | Quad3Prop1 | | | | | upper | Proportion of total spot area in Quadrant 3 |
| | | | | | | | *PropSpotsQ3* |
| | Quad3Prop2 | upper | | upper | | | Proportion of Quadrant 3 covered by spots |
| | | lower | | lower | | | *Q3CoveredBySpots* |
| | Quad4Num | | | | | upper | Number of spots with centers of mass in Quadrant 4 |
| | | | | | | | *NuQ4spots* |
| | Quad4Prop1 | | | | | | Proportion of total spot area in Quadrant 4 |
| | | | | | | lower | *PropSpotsQ4* |
| | Quad4Prop2 | upper | | upper | | | Proportion of Quadrant 4 covered by spots |
| | | lower | | lower | | | *Q4CoveredBySpots* |

### avg Dist2Center All Spots

#### Upper Petal

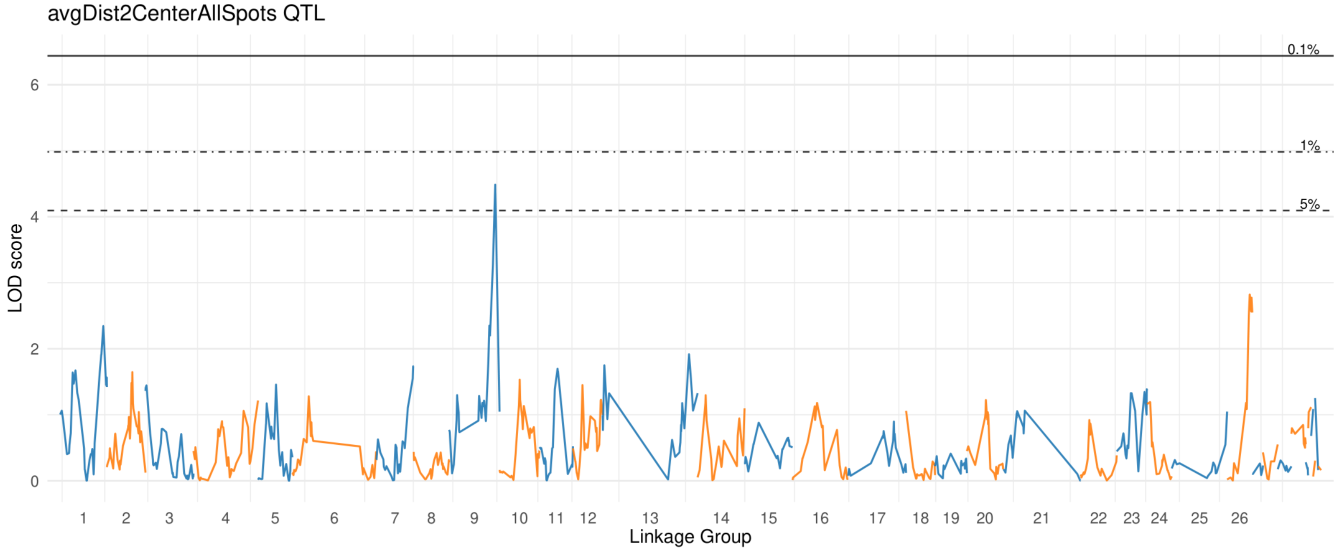


#### Lower Petal

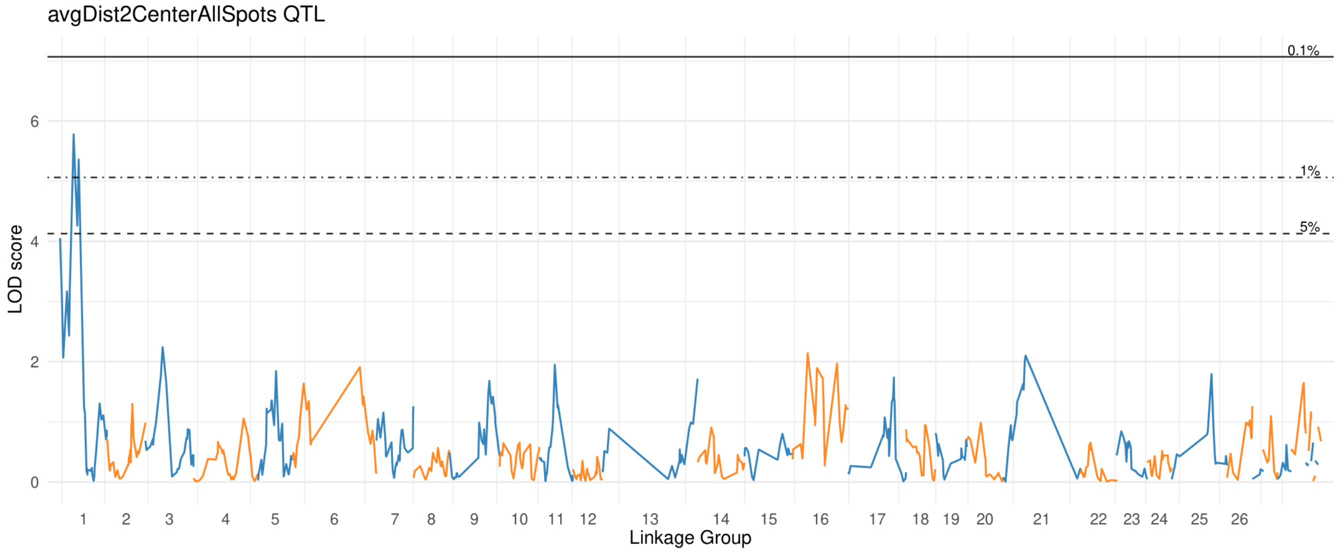

### avg Dist2Center Center Spots

#### Upper Petal

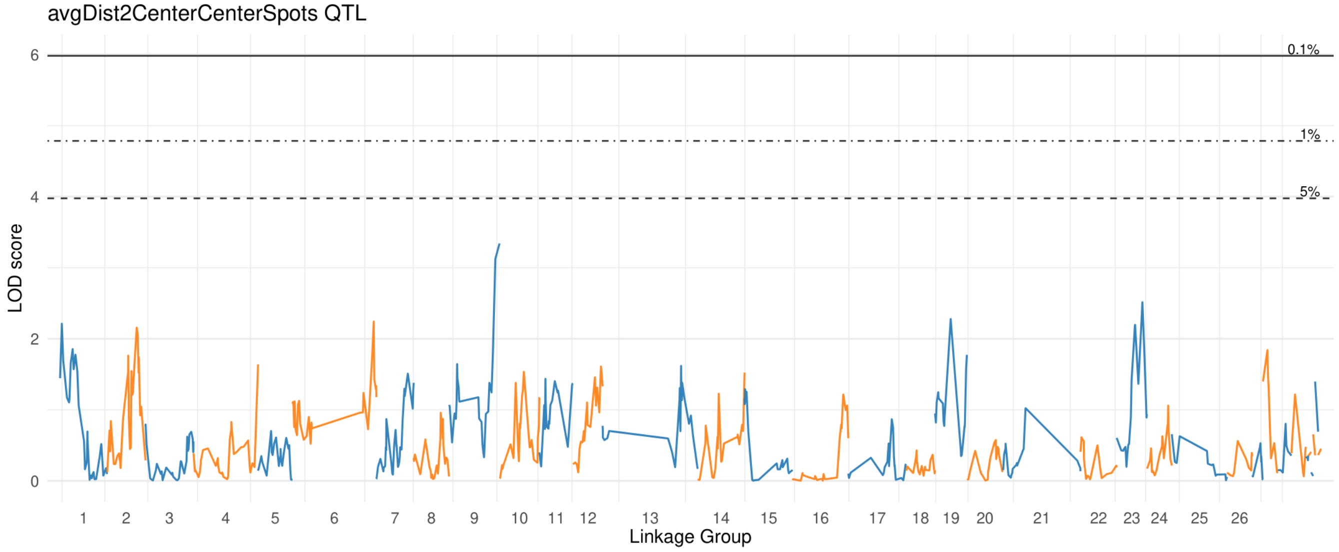


#### Lower Petal

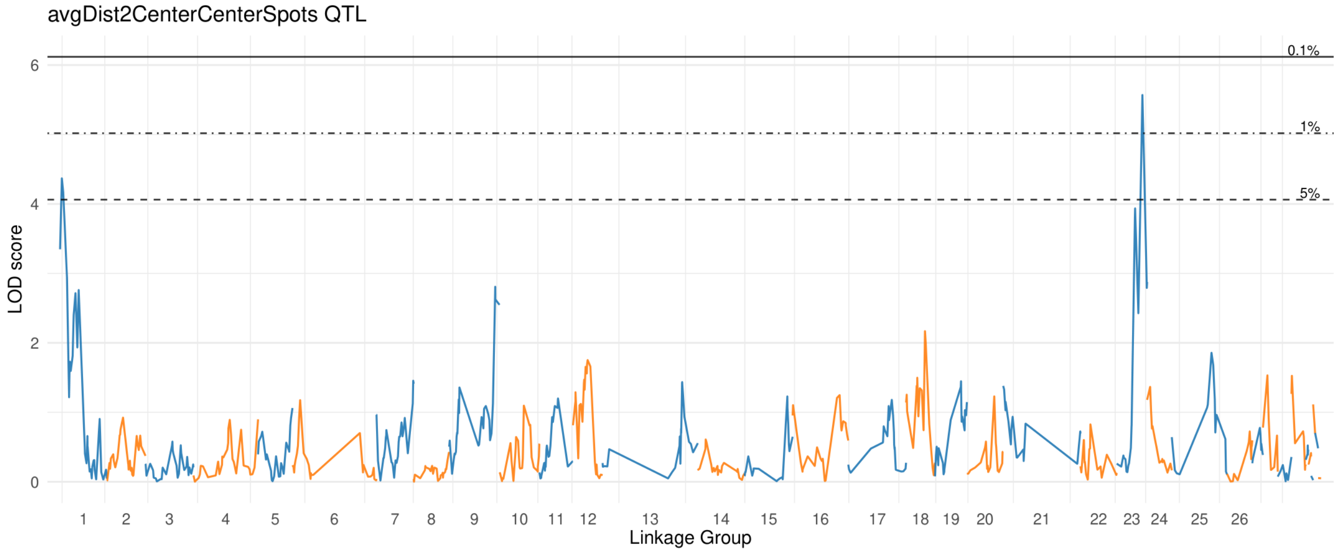

### avg Dist Spot Centroid2Edge

#### Upper Petal

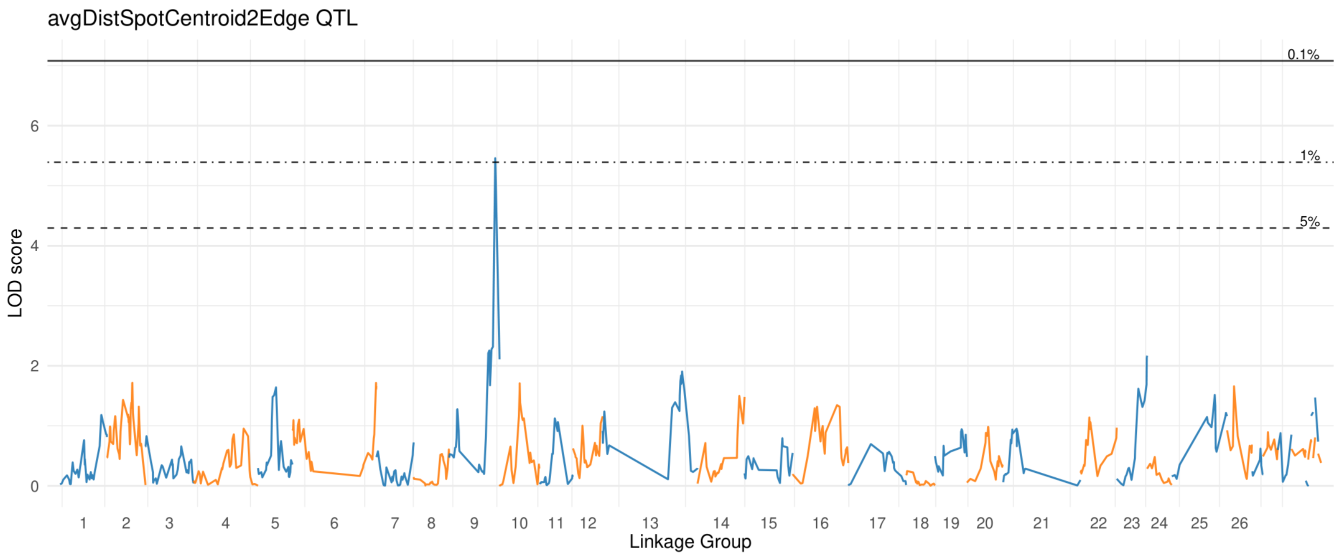


#### Lower Petal

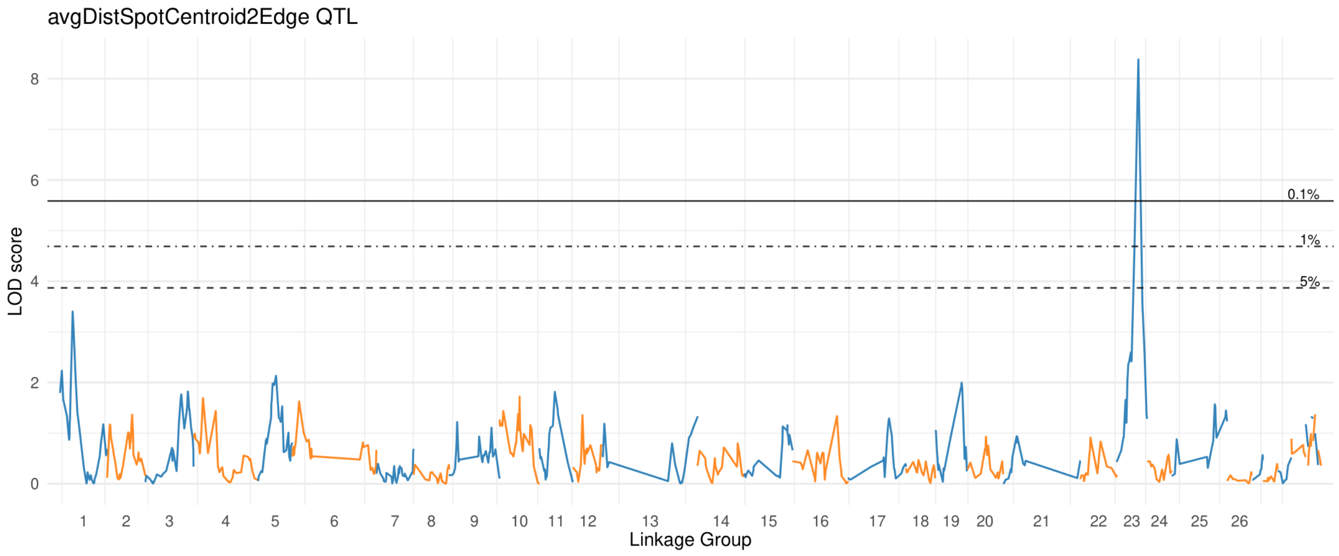

### avg Dist Spot Edge2Edge

#### Upper Petal

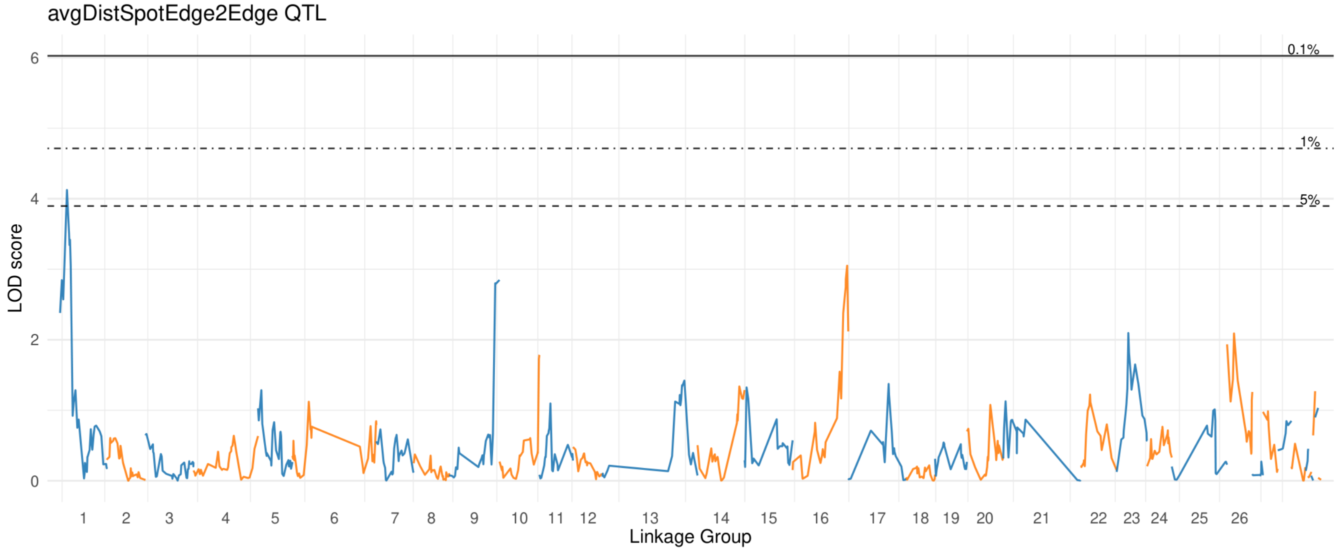


#### Lower Petal

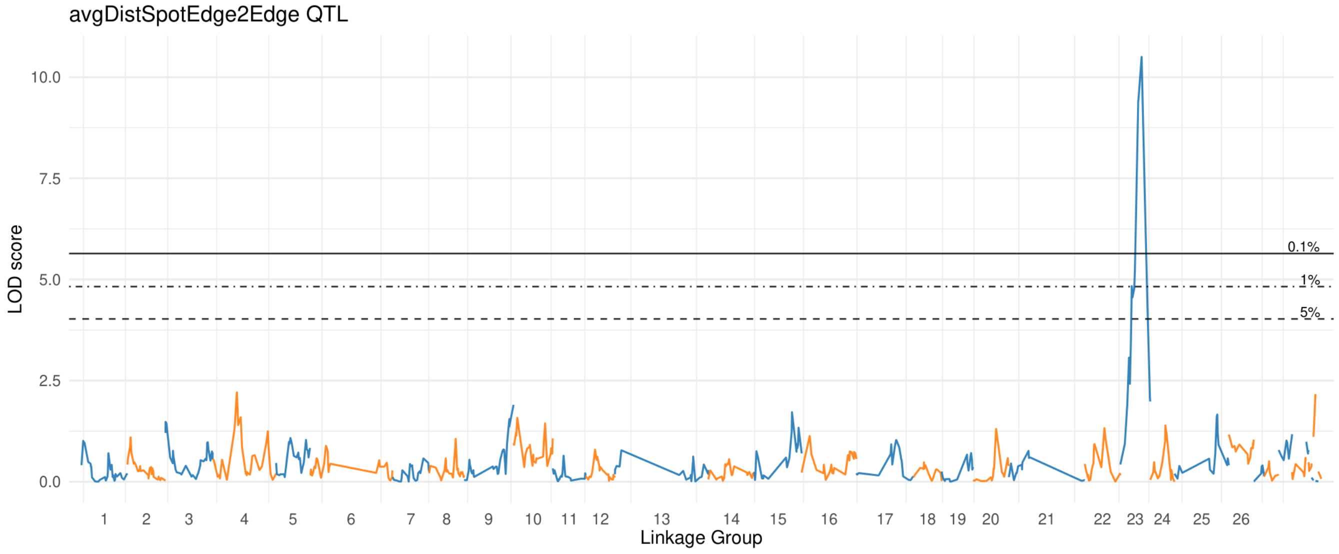

### avg Spot Size

#### Upper Petal

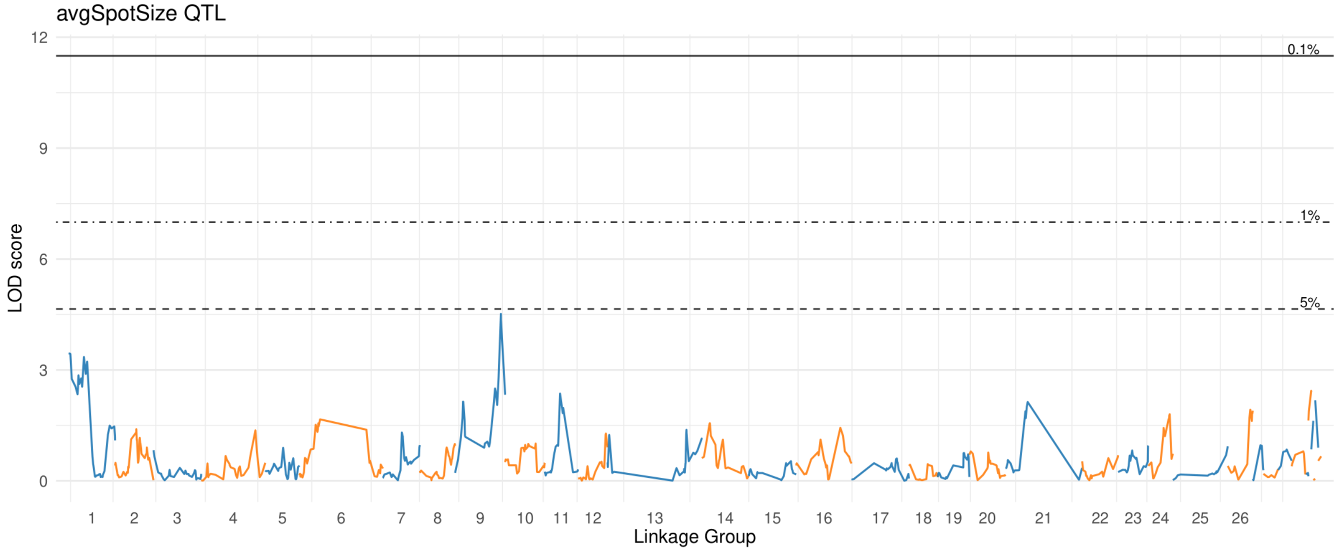


#### Lower Petal

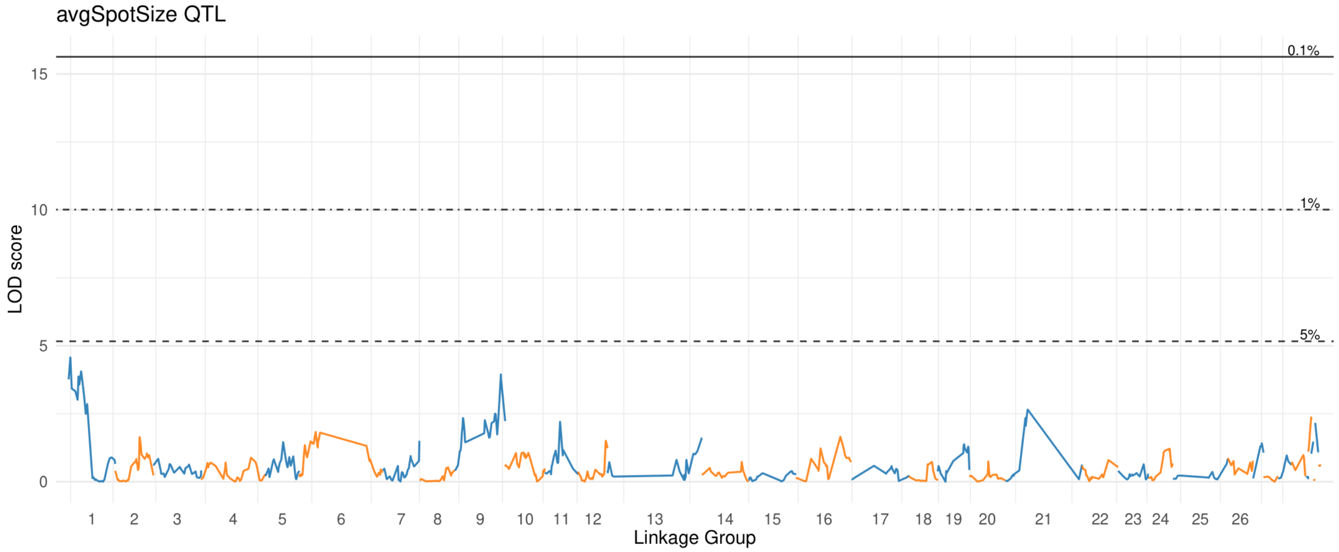

### biggest Spot Area

#### Upper Petal

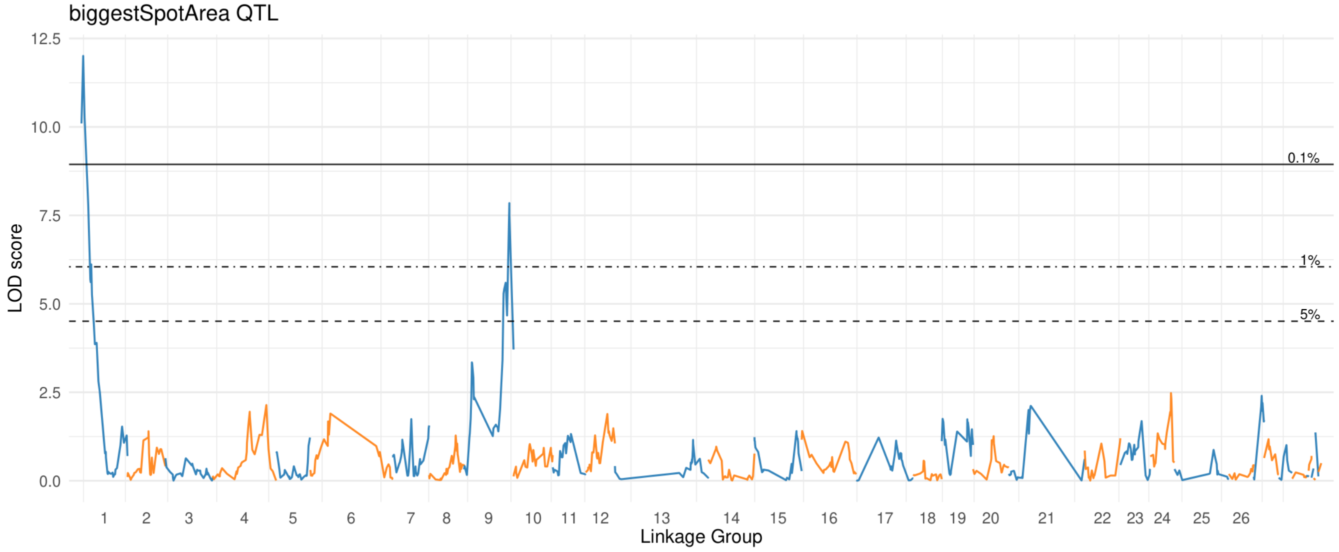


#### Lower Petal

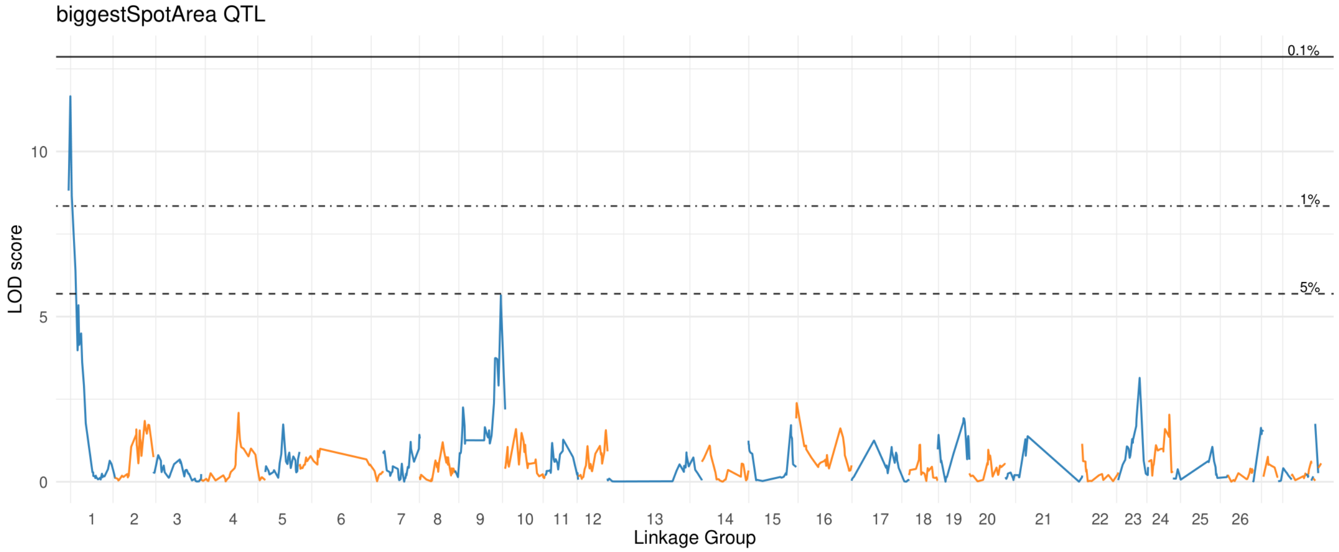

### center Coveredby Spots

#### Upper Petal

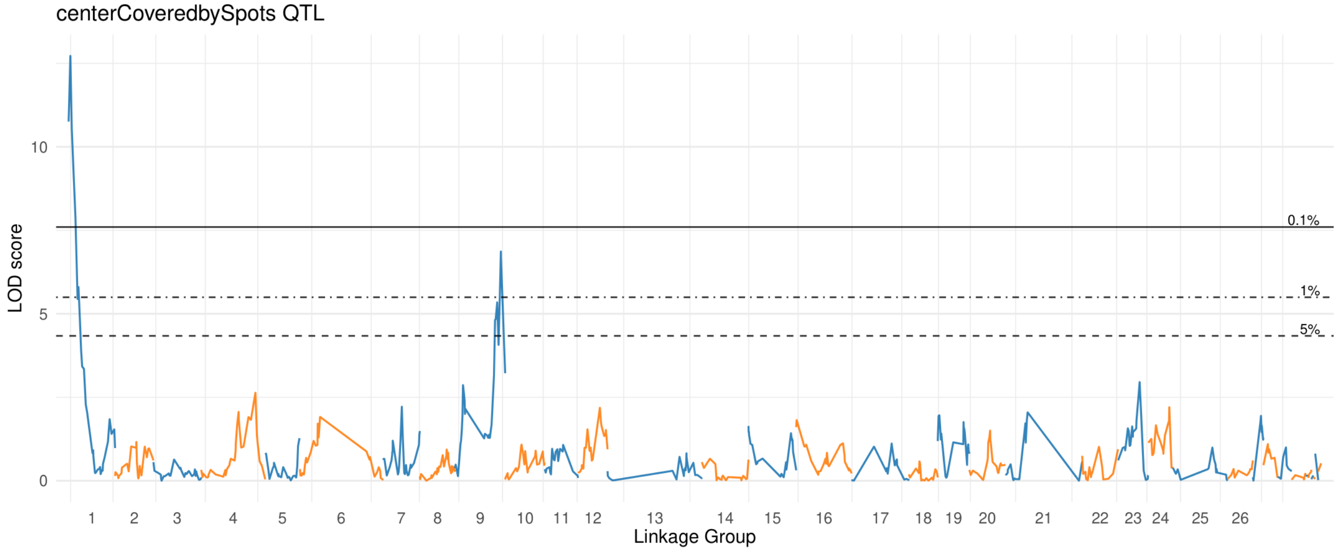


#### Lower Petal

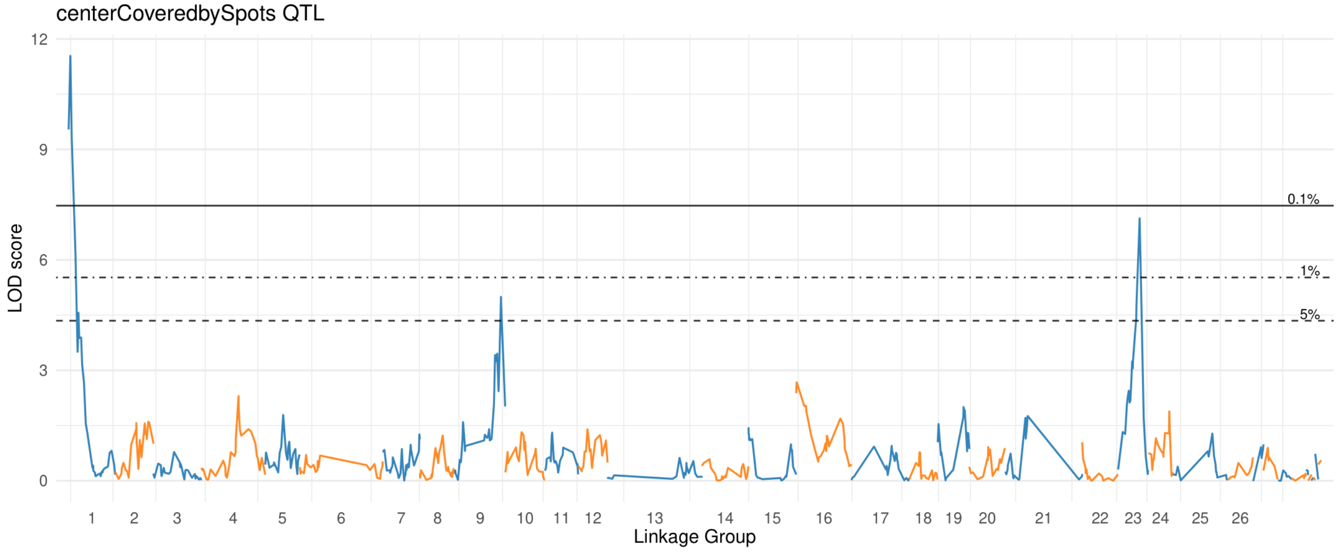

### dist Coveredby Spots

#### Upper Petal

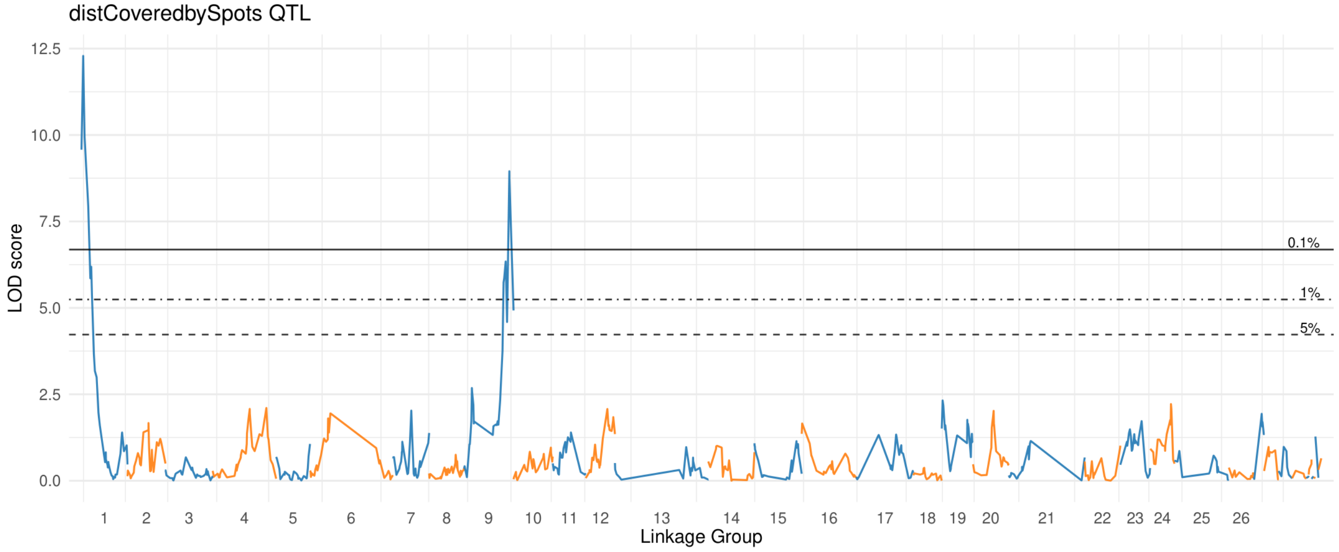


#### Lower Petal

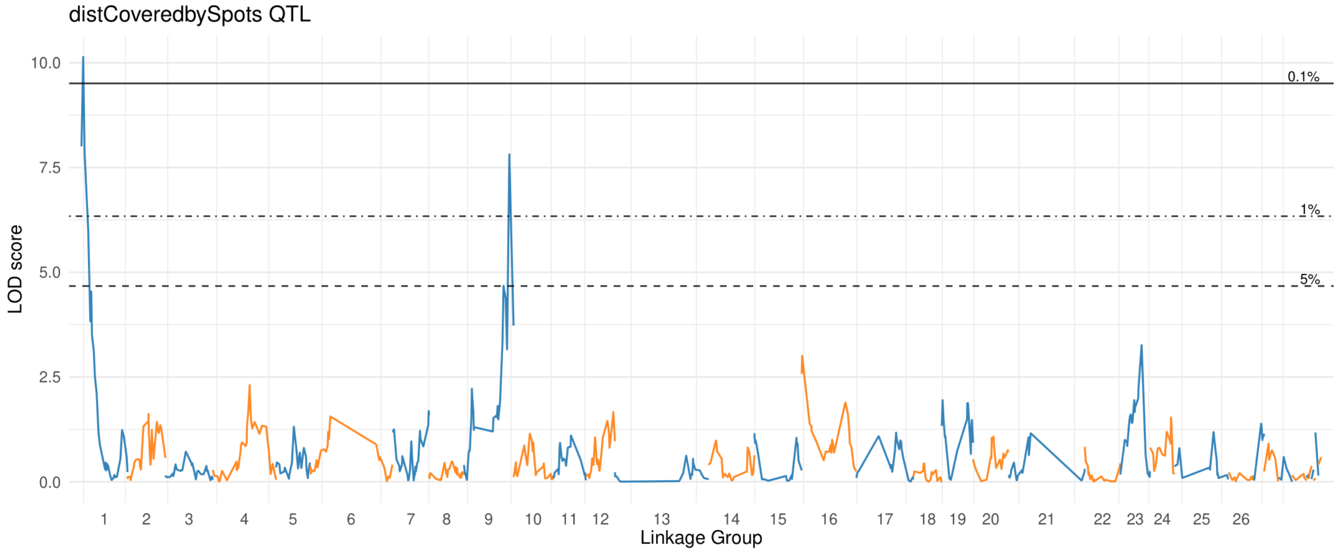

### edge Coveredby Spots

#### Upper Petal

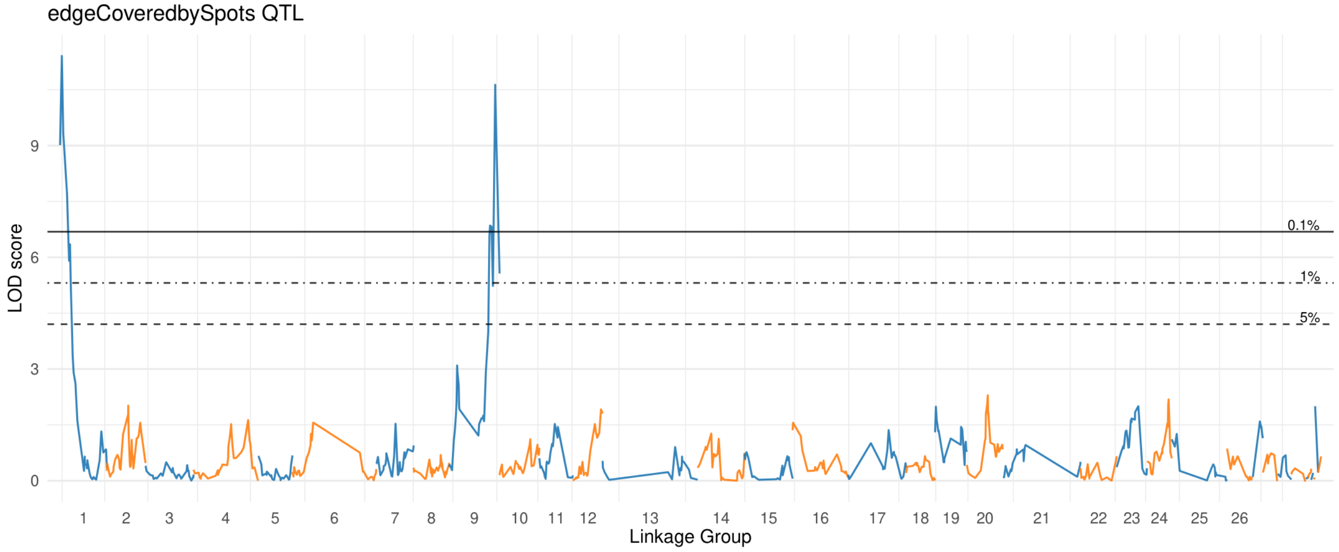


#### Lower Petal

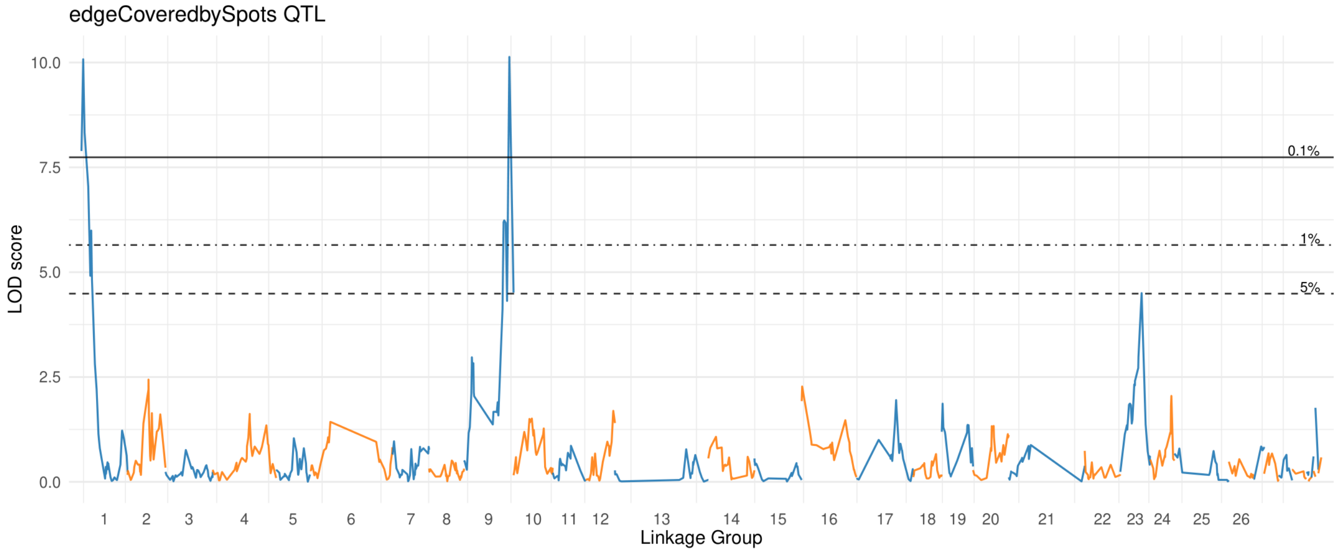

### med Spot Size

#### Upper Petal

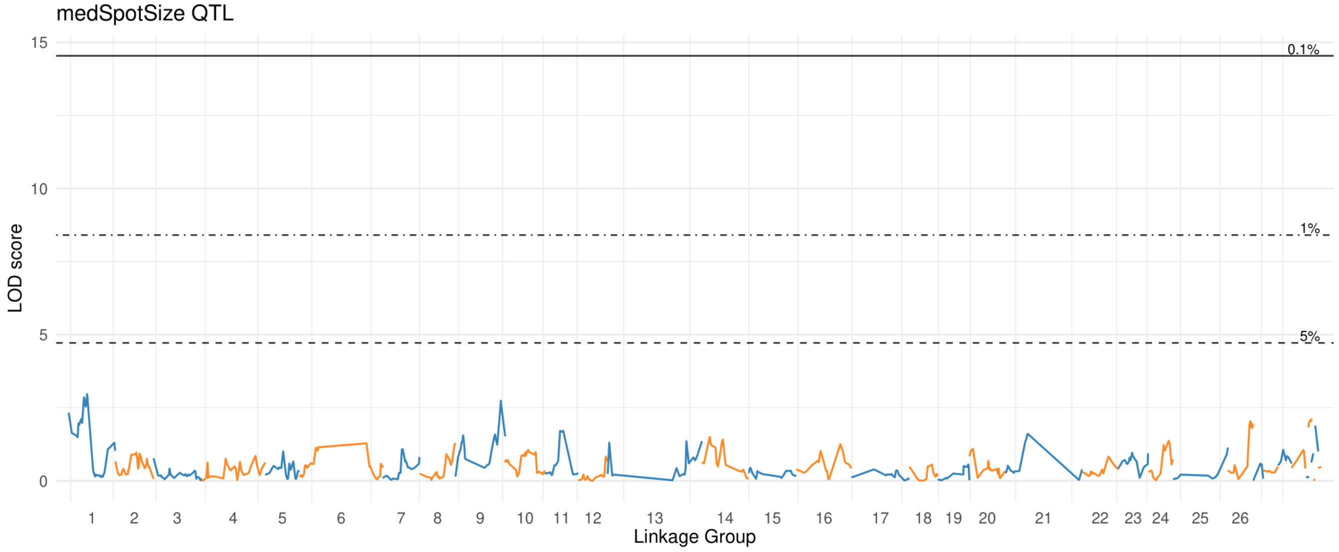


#### Lower Petal

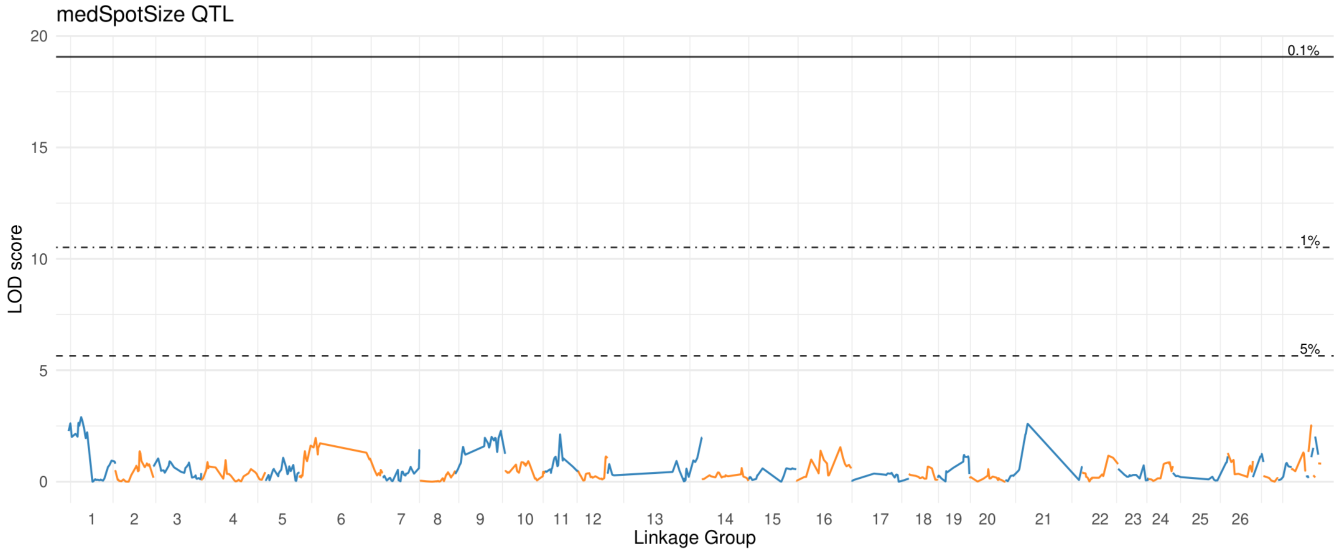

### nu Dist Spots

#### Upper Petal

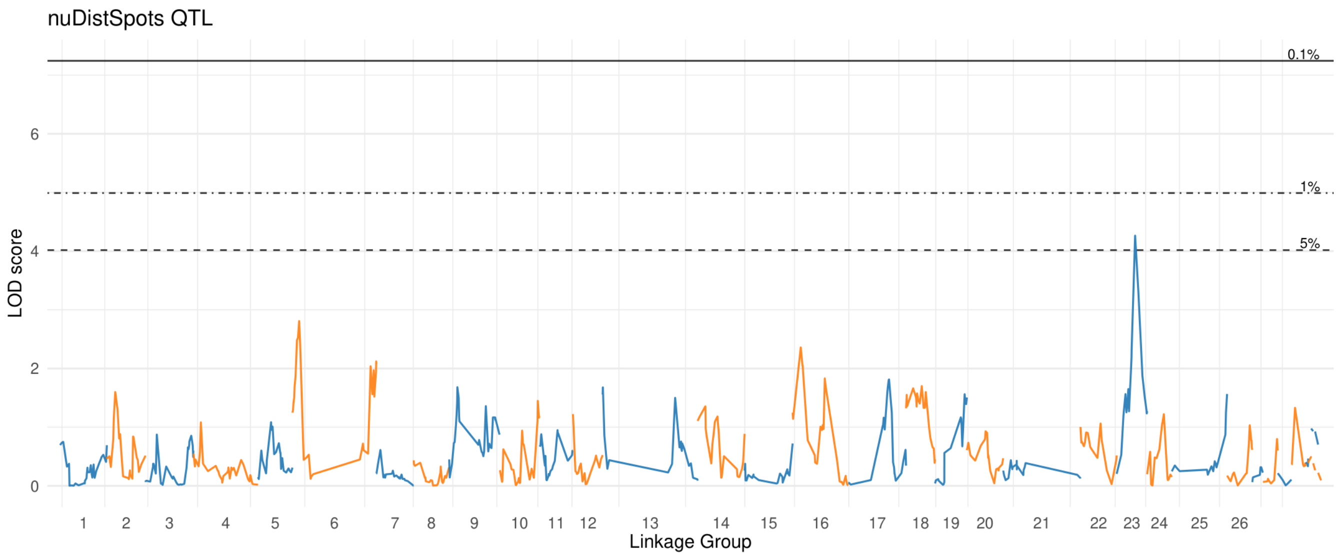


#### Lower Petal

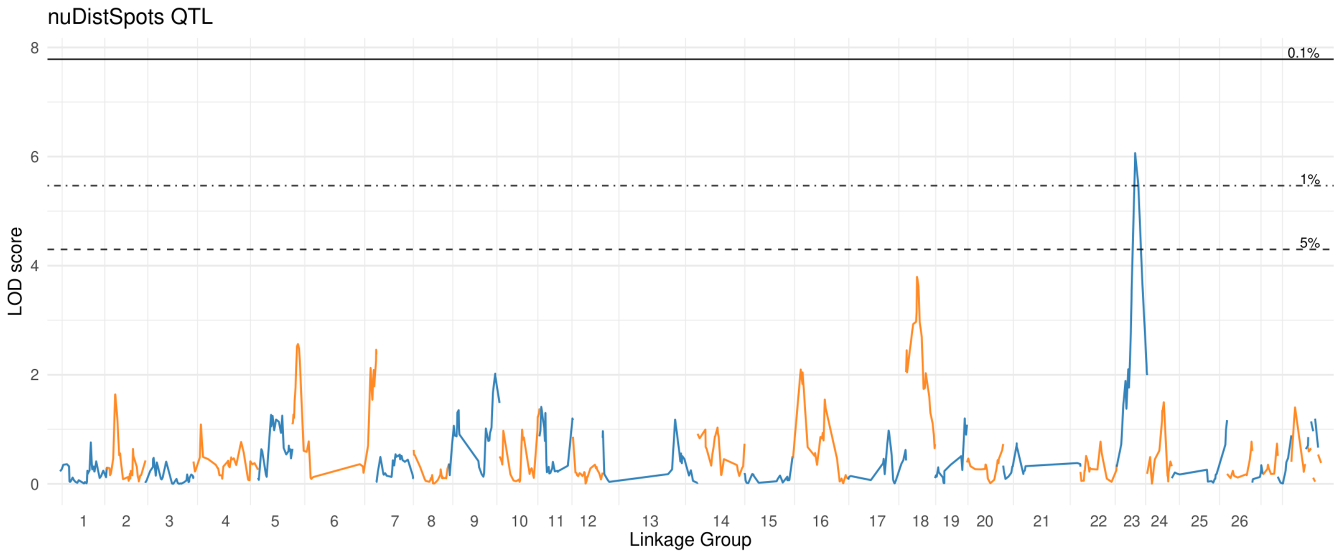

### nu Prox Spots

#### Upper Petal

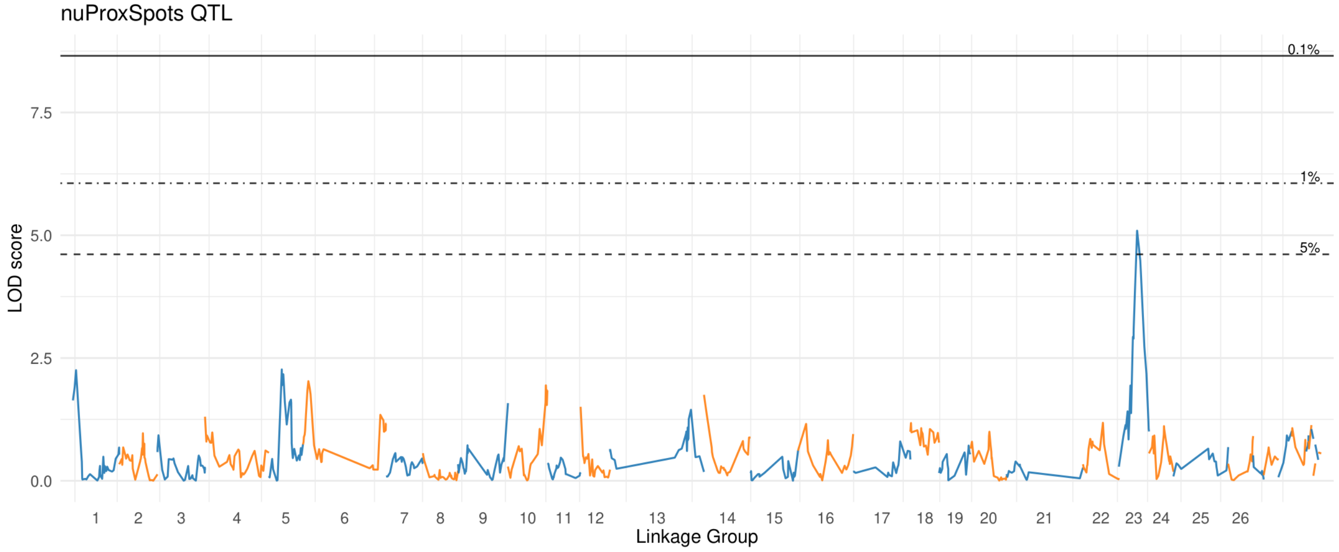


#### Lower Petal

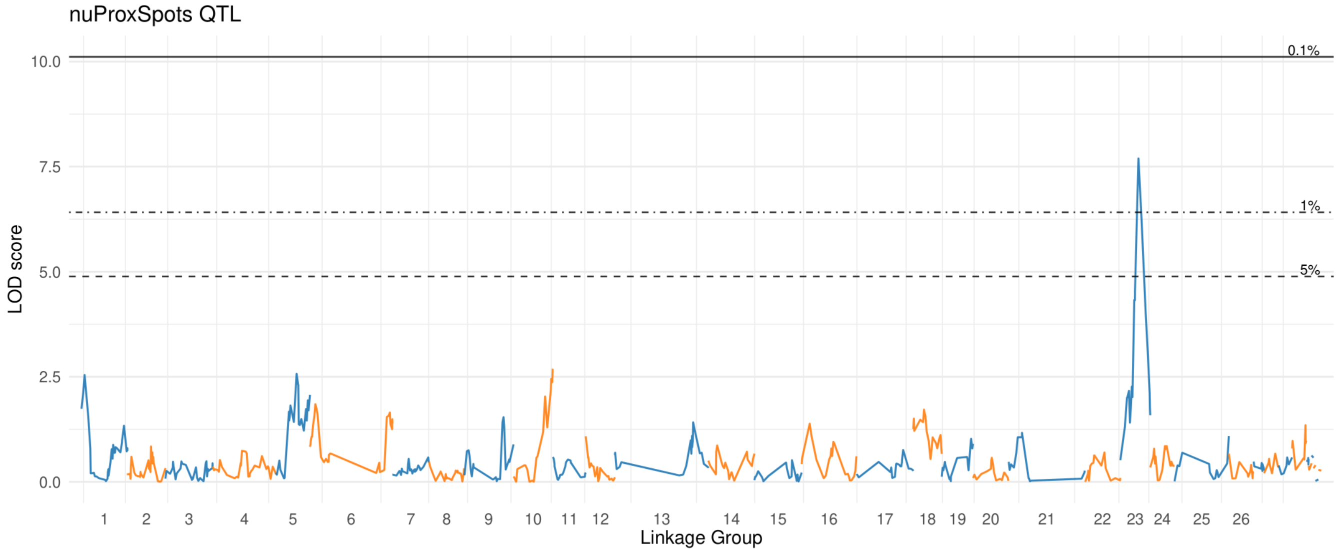

### nu Quad IIISpots

#### Upper Petal

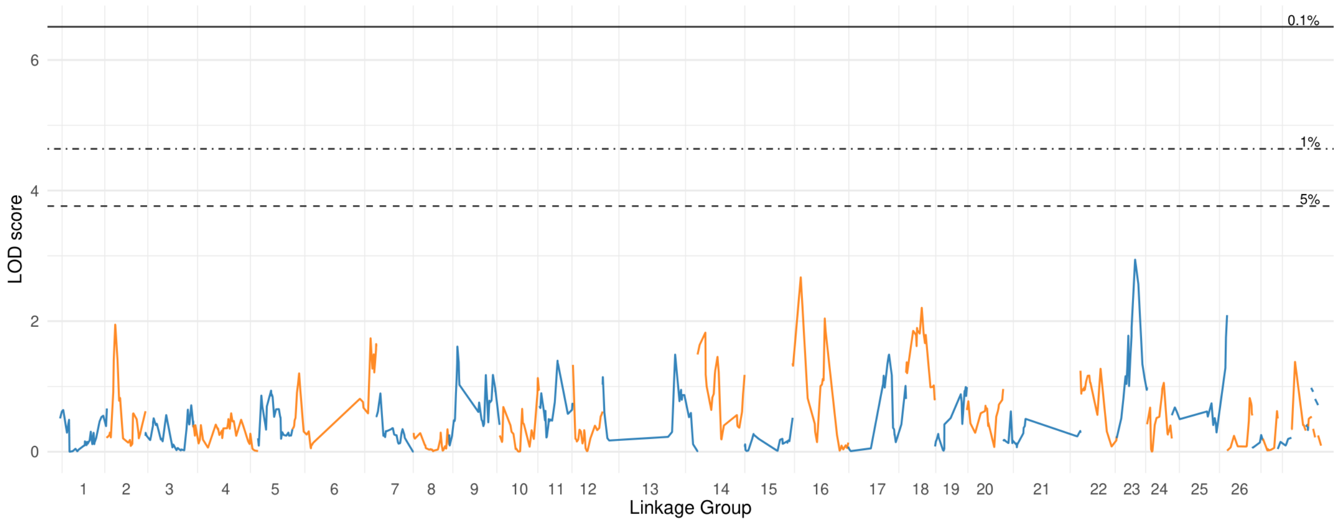


#### Lower Petal

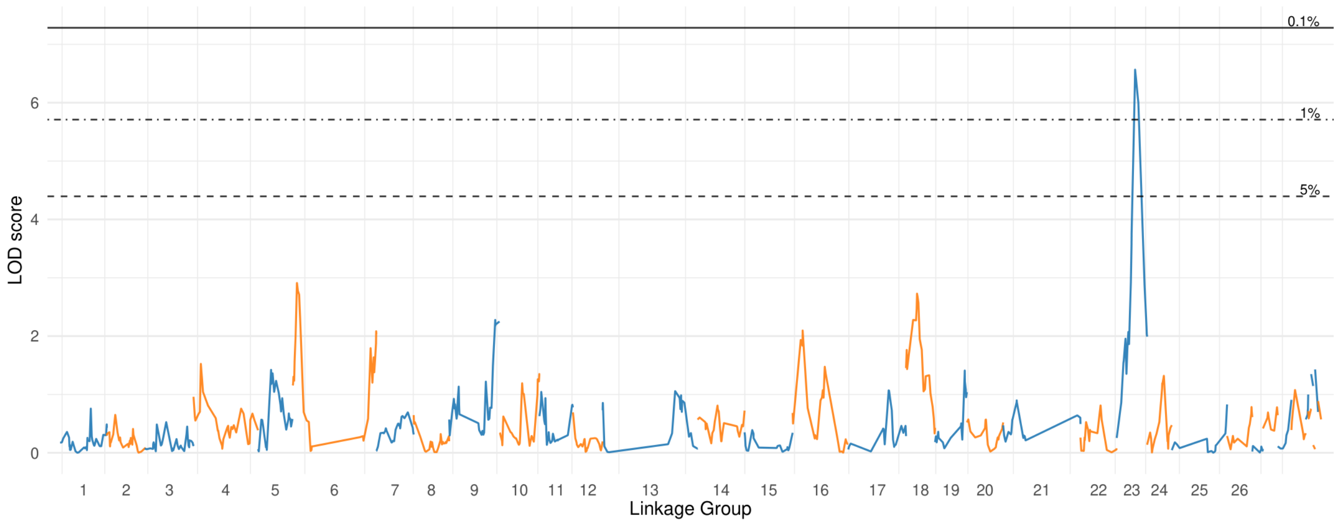

### nu Quad IISpots

#### Upper Petal

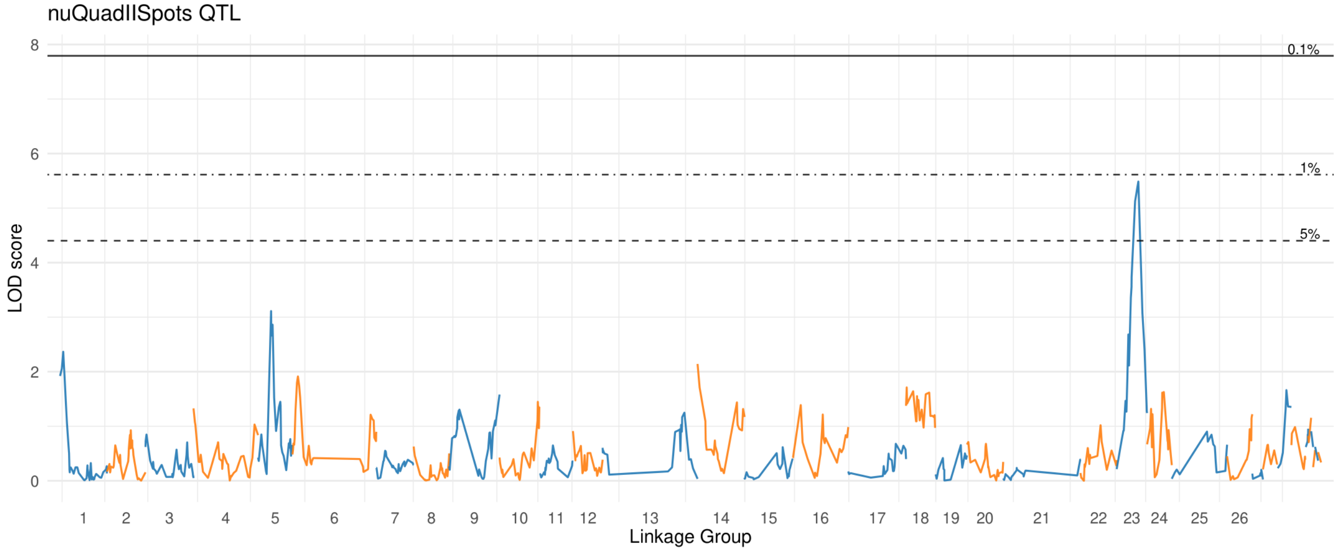


#### Lower Petal

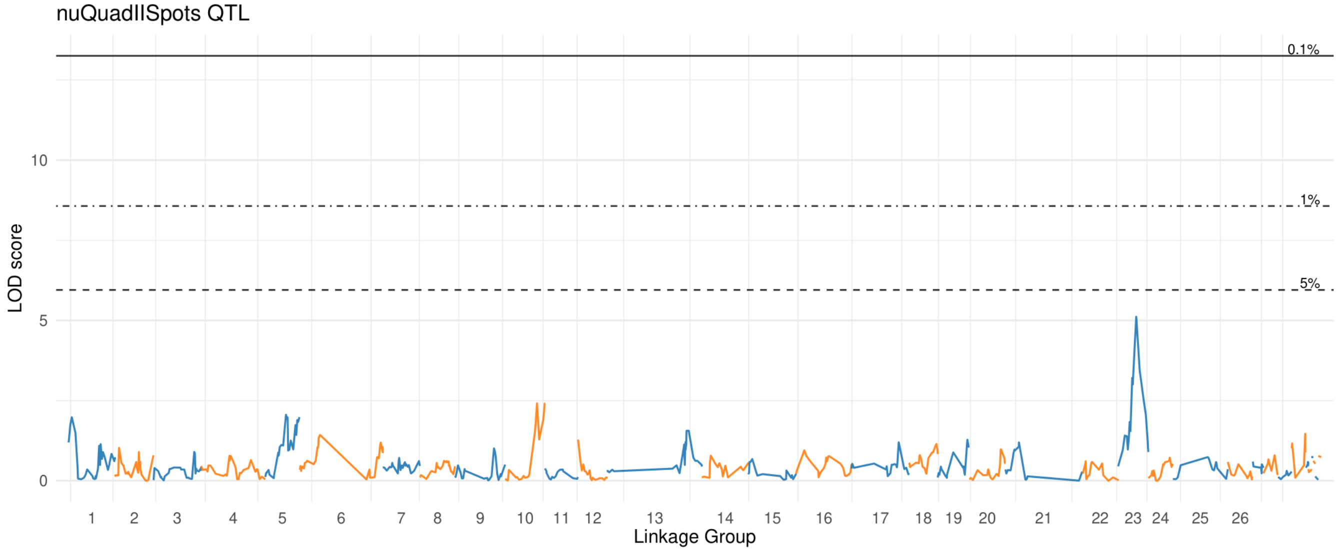

### nu Quad ISpots

#### Upper Petal

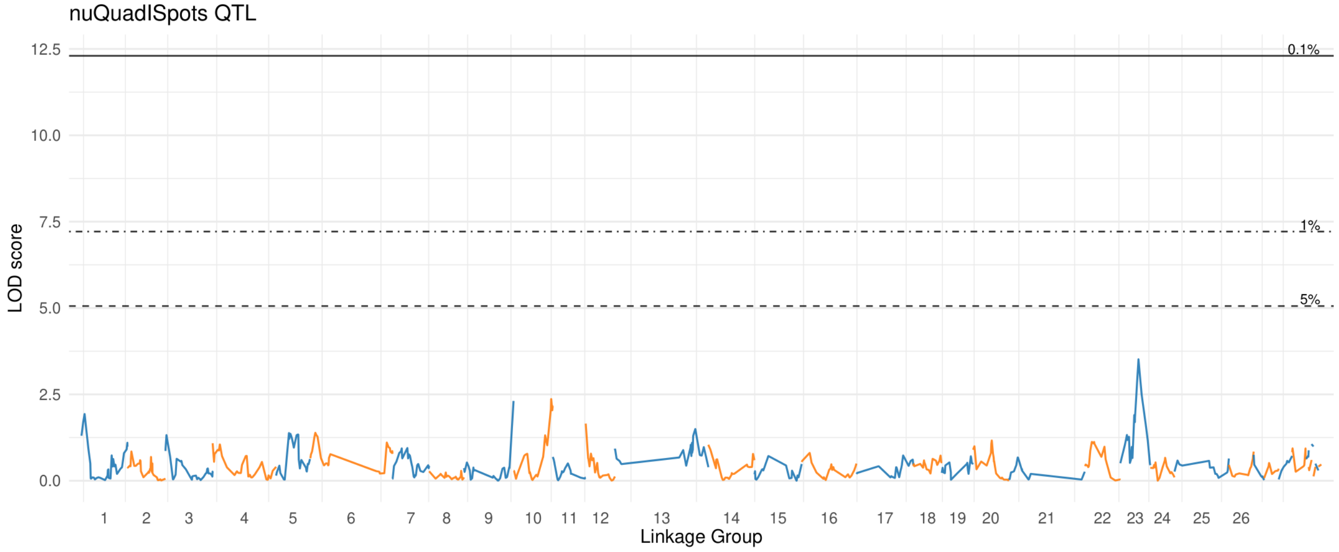


#### Lower Petal

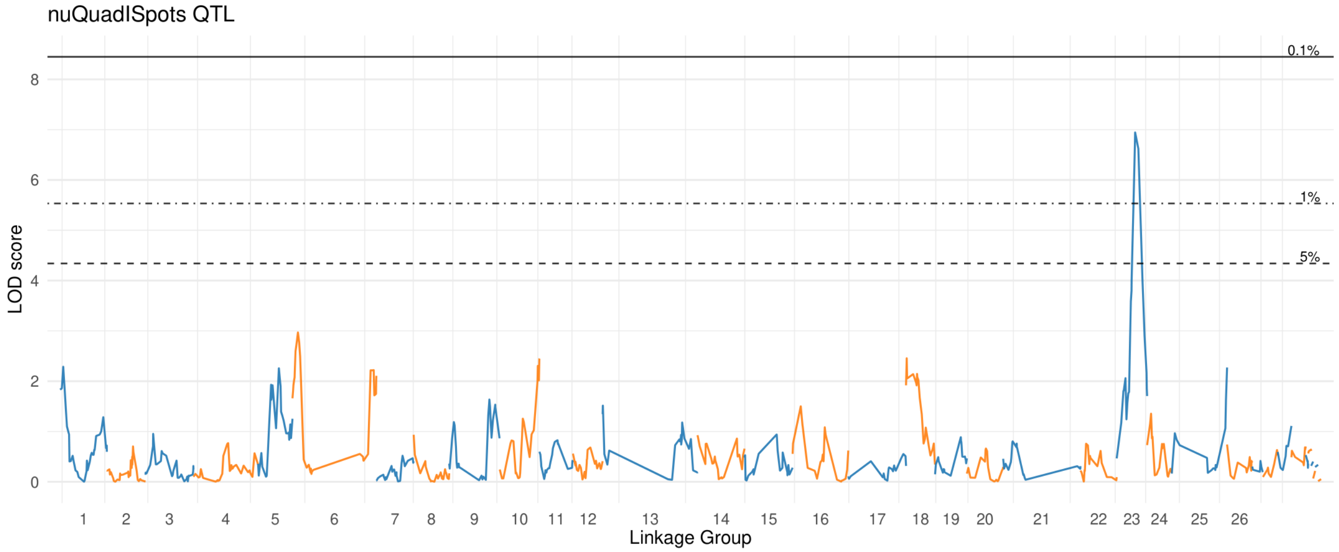

### nu Quad IVSpots

#### Upper Petal

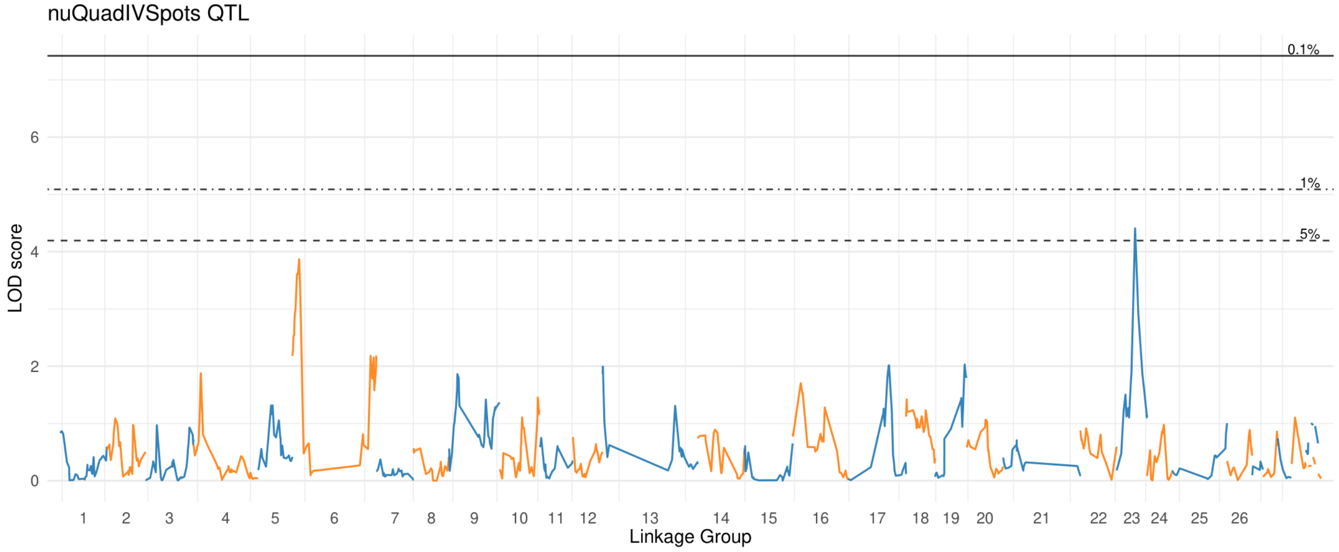


#### Lower Petal

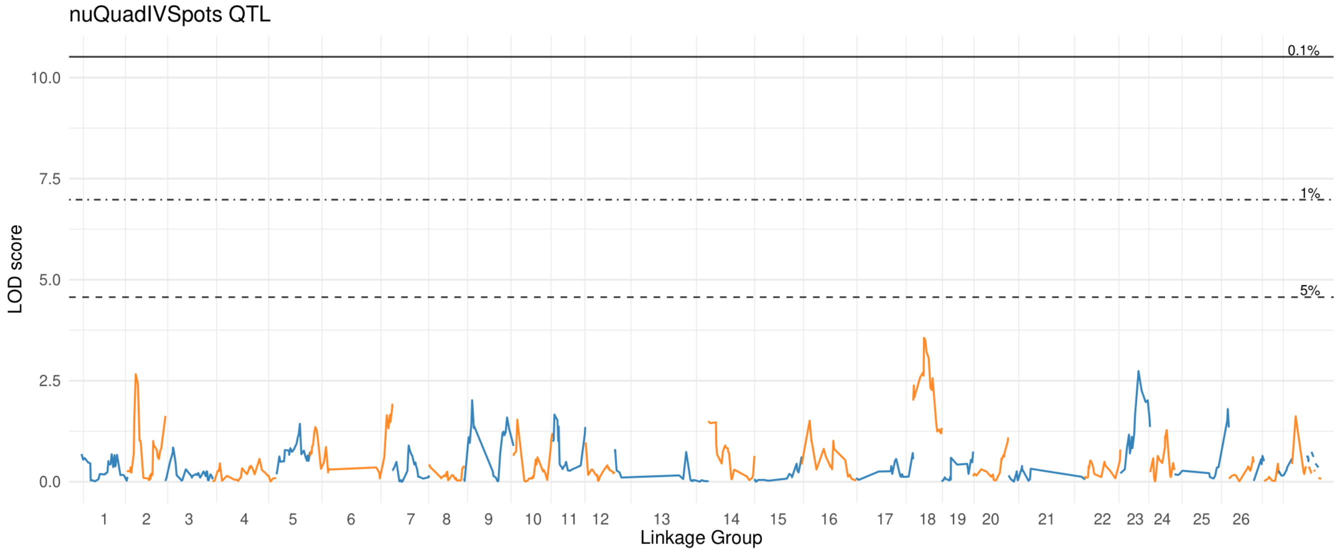

### nu Spot Centroids In Center

#### Upper Petal

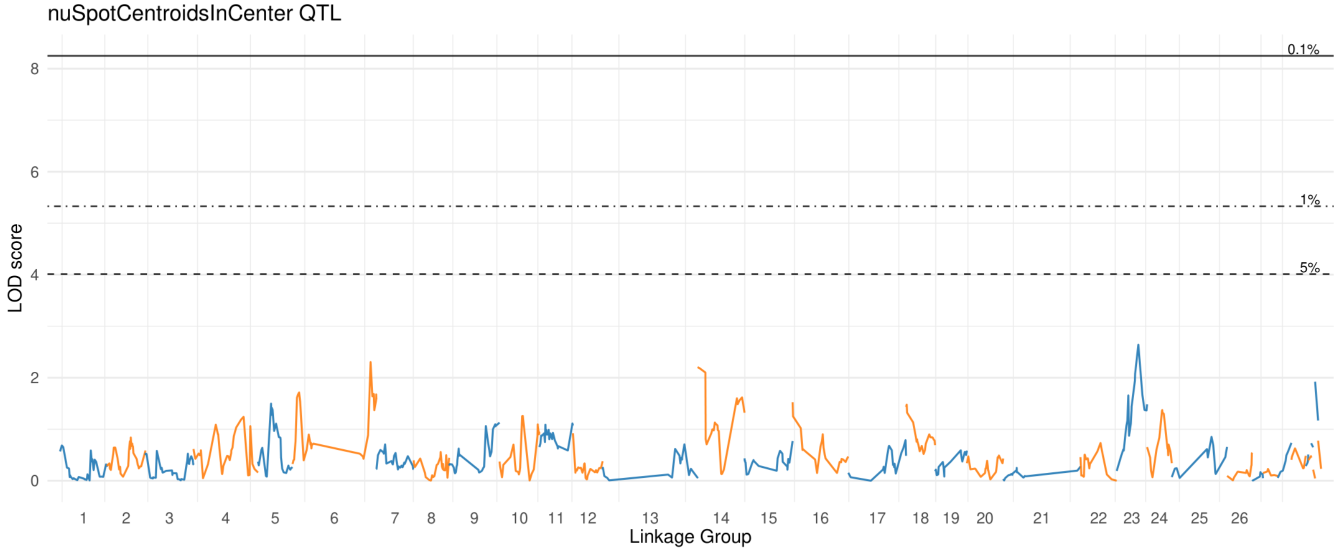


#### Lower Petal

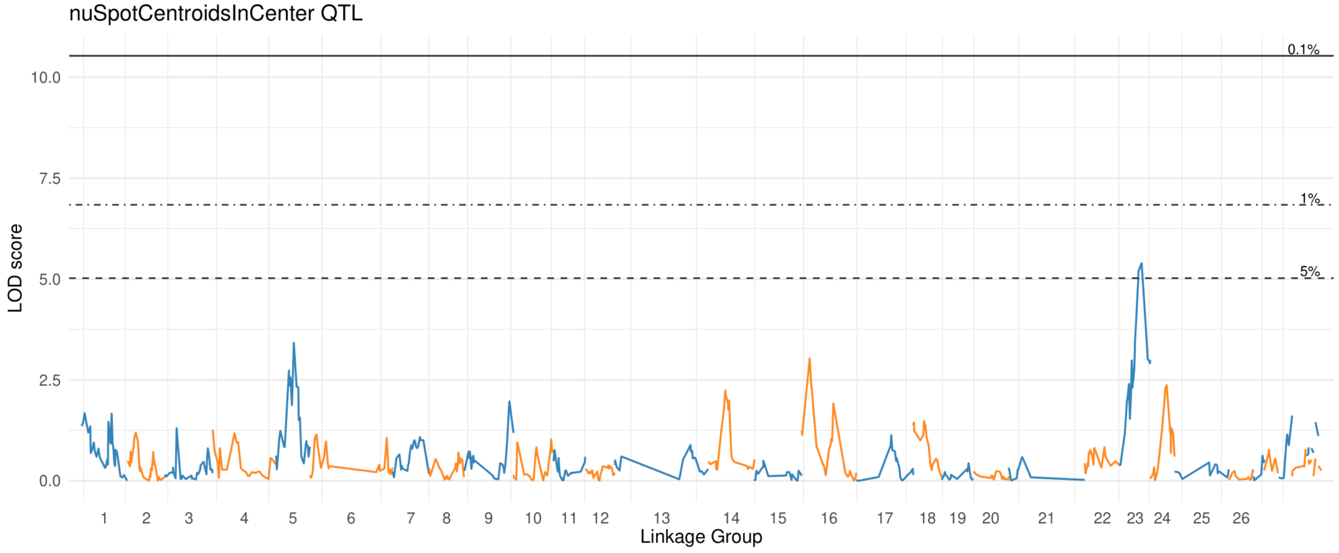

### nu Spots

#### Upper Petal

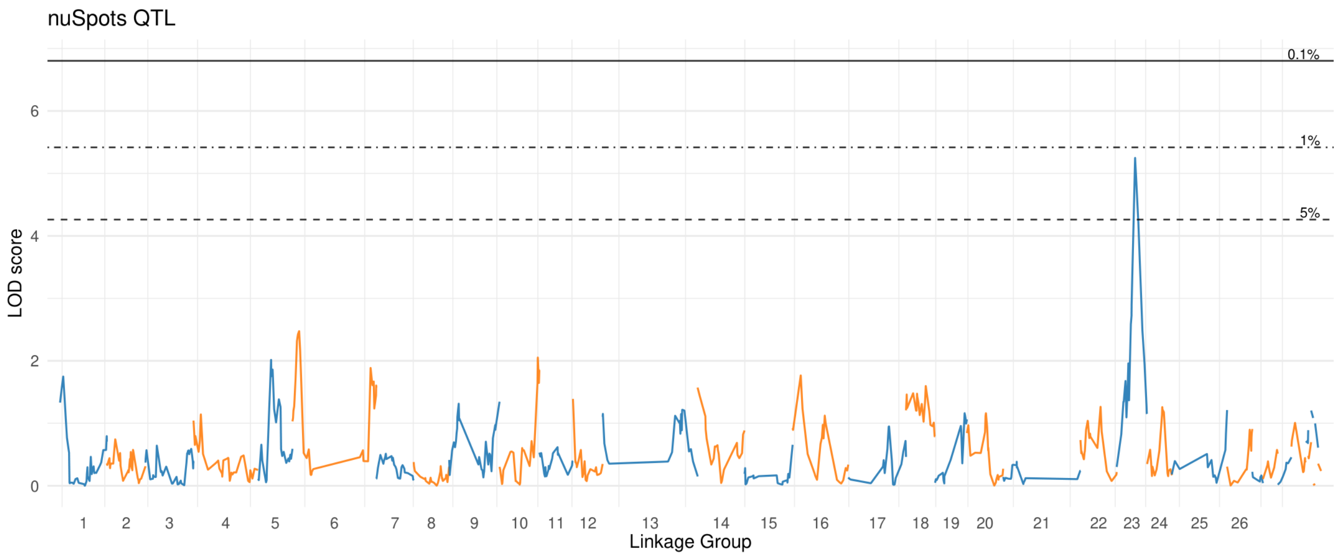


#### Lower Petal

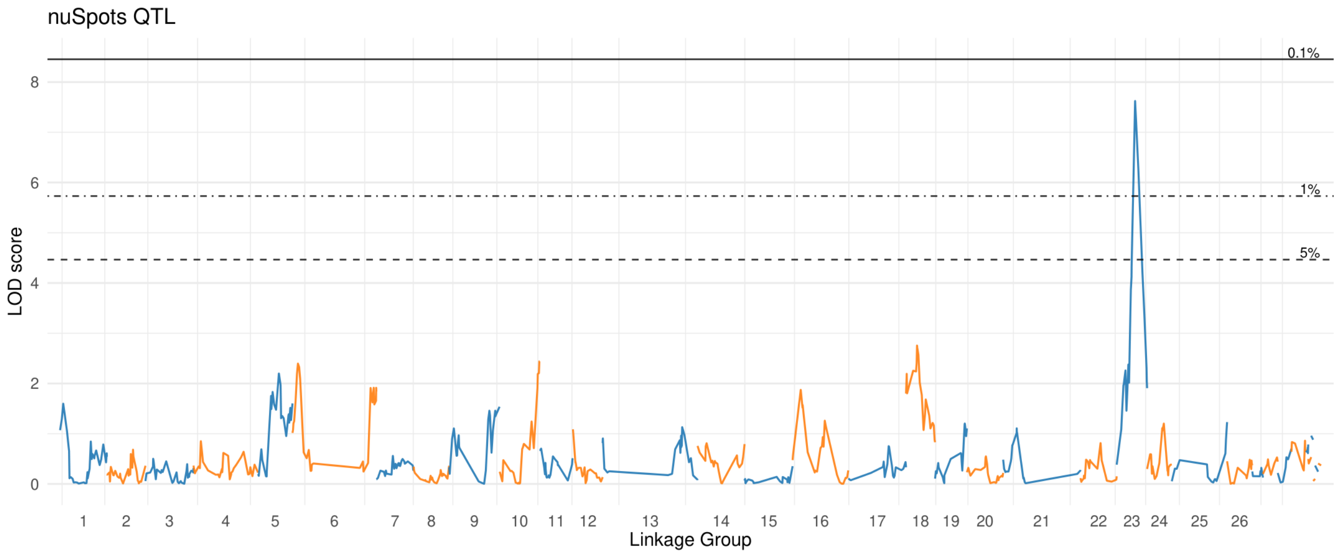

### nu Spots Contained In Center

#### Upper Petal

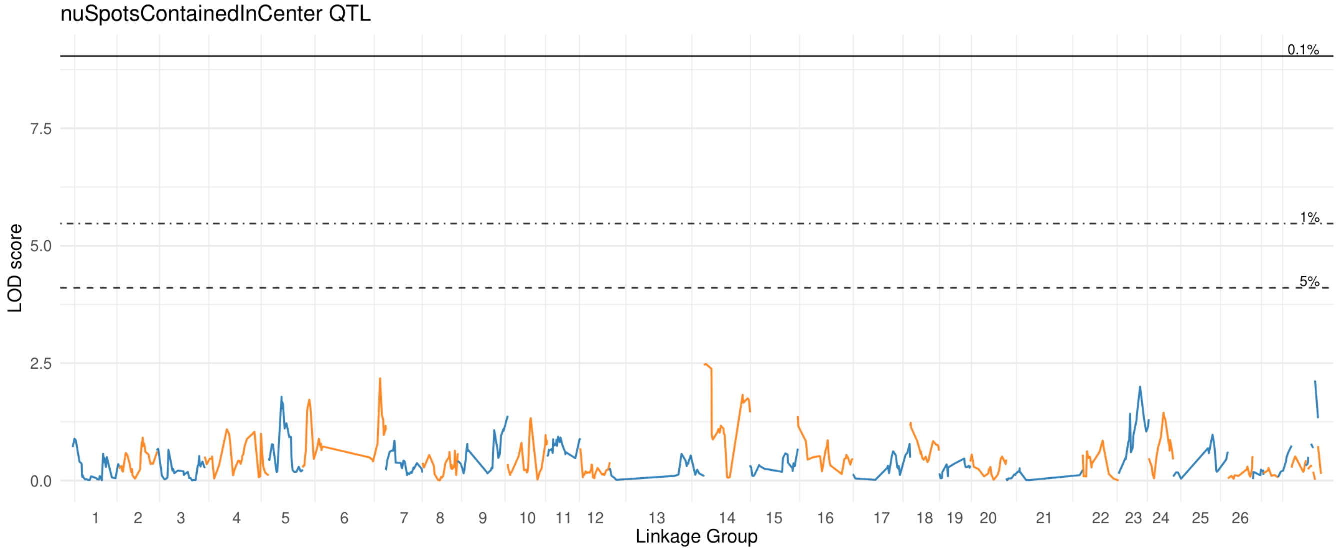


#### Lower Petal

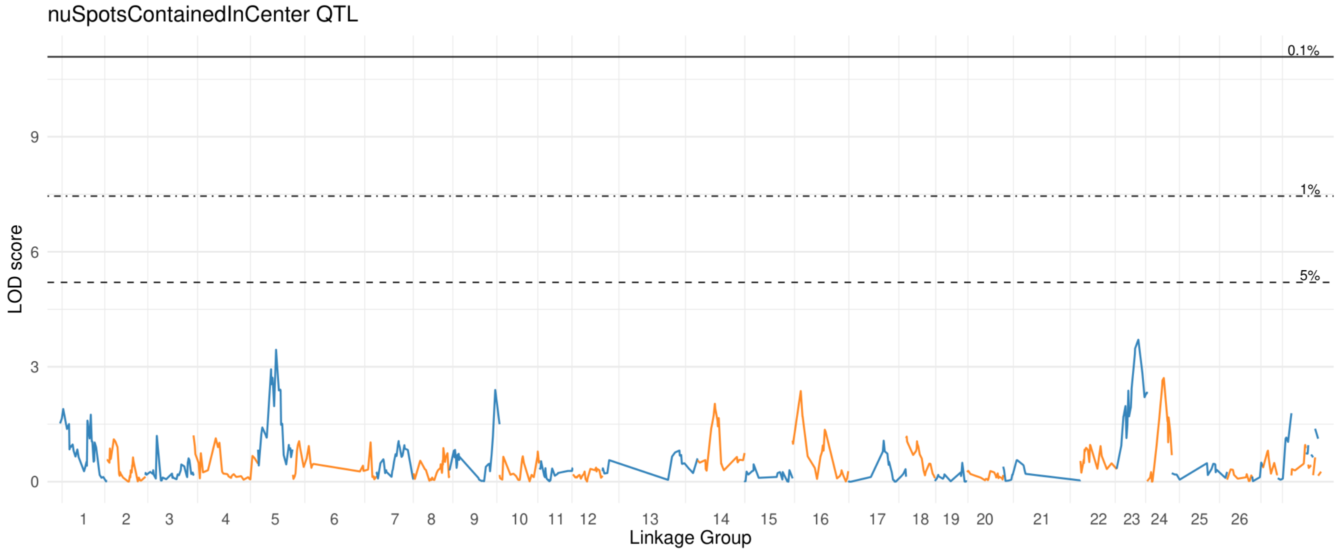

### nu Spots Contained In Edge

#### Upper Petal

nuSpotsContainedInEdge QTL

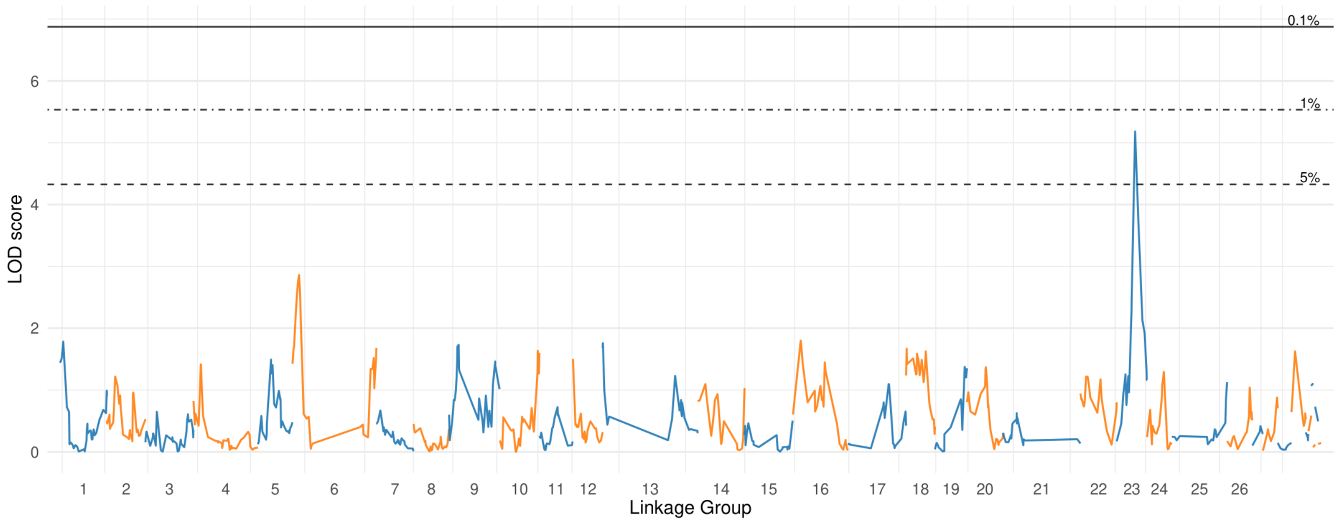


#### Lower Petal

nuSpotsContainedInEdge QTL

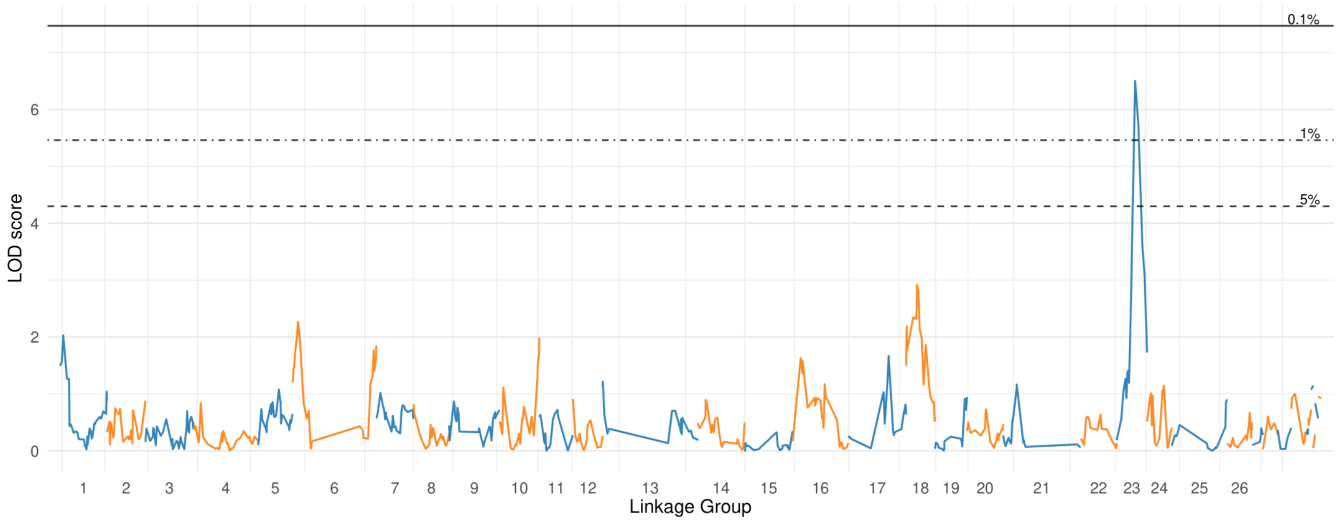

### nu Spots Mostly In Center

#### Upper Petal

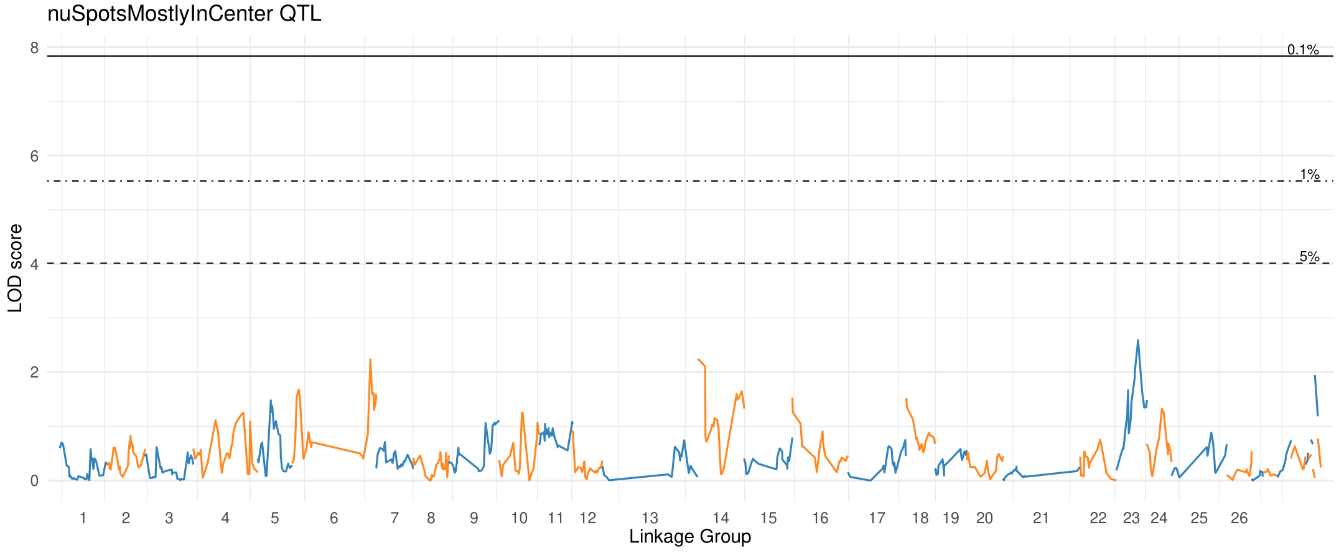


#### Lower Petal

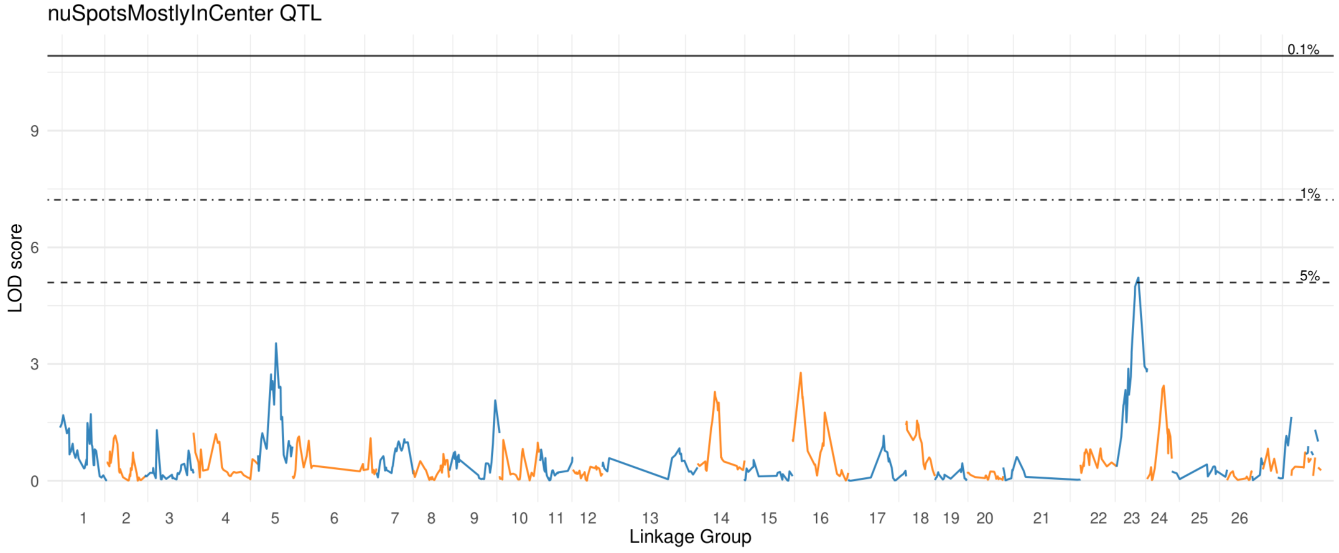

### nu Spots Mostly In Edge

#### Upper Petal

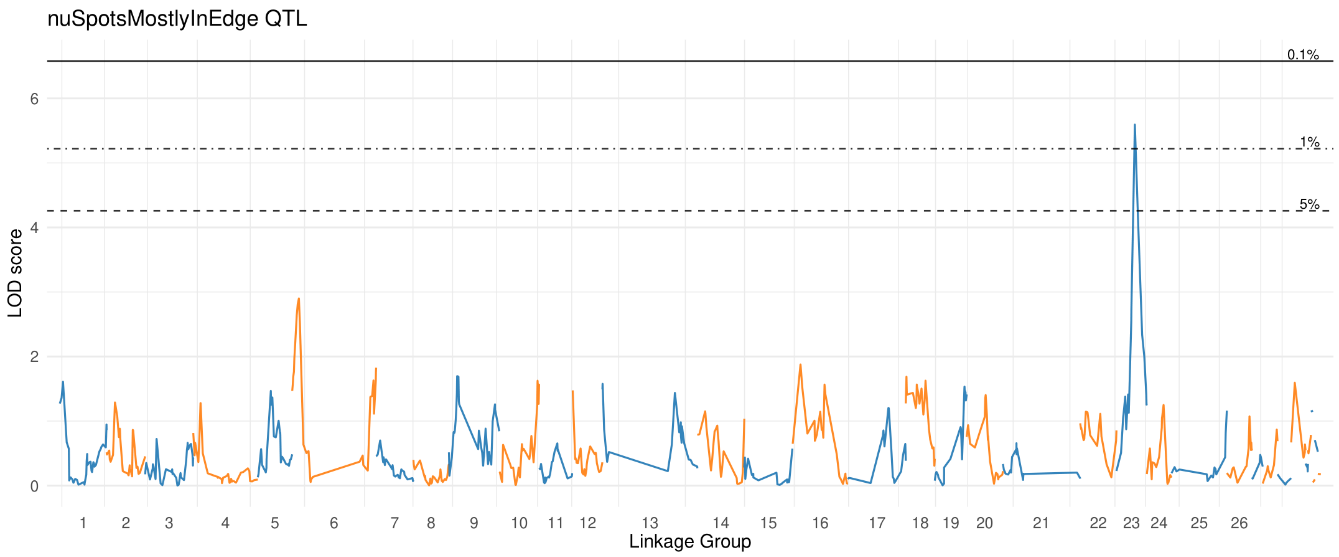


#### Lower Petal

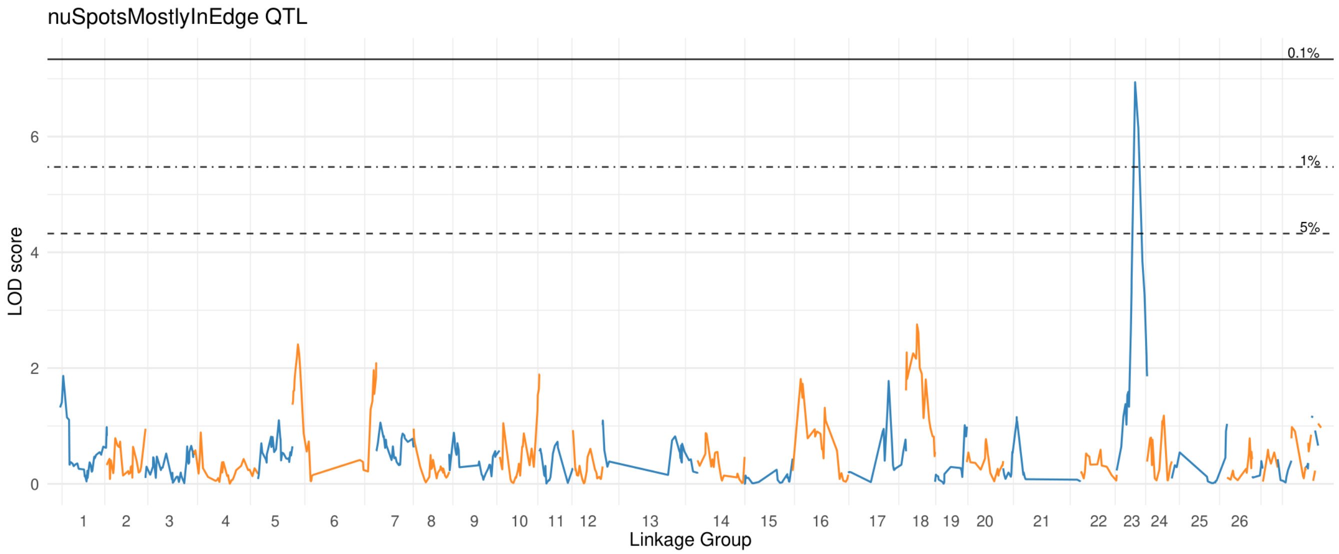

### nu Spots Mostly In Throat

#### Upper Petal

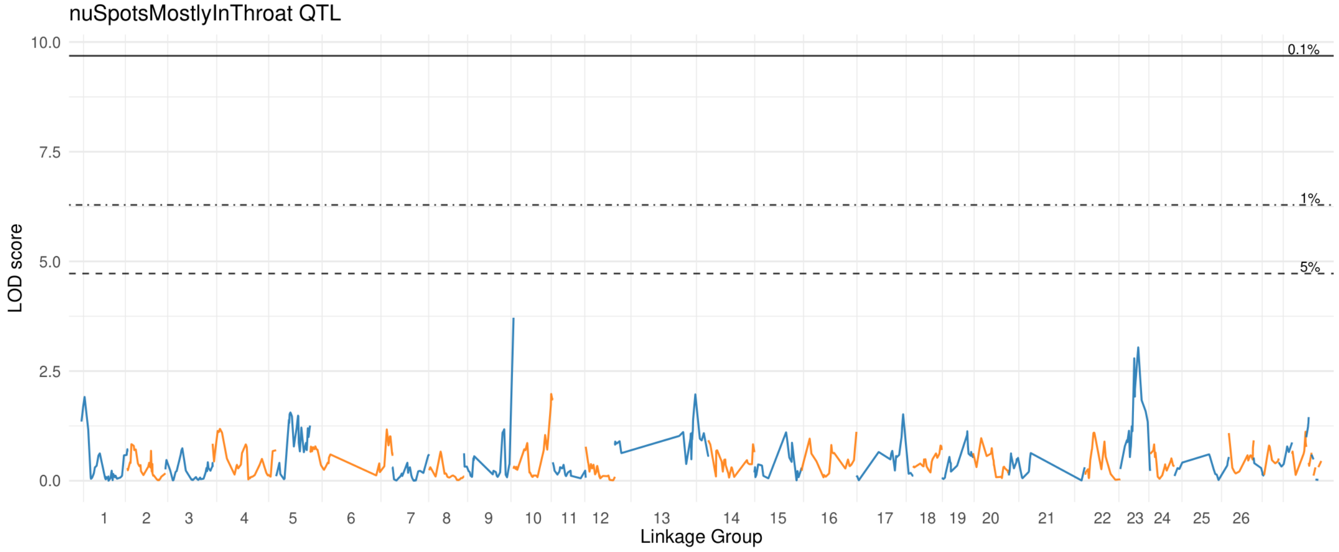


#### Lower Petal

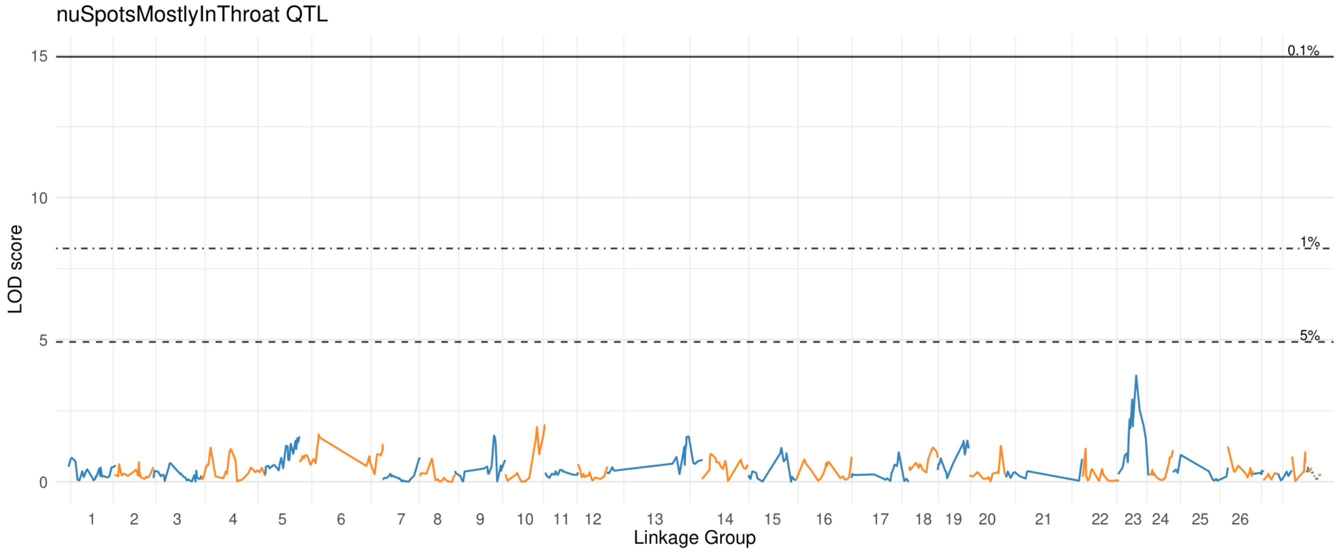

### nu Spots Touch Actual Edge

#### Upper Petal

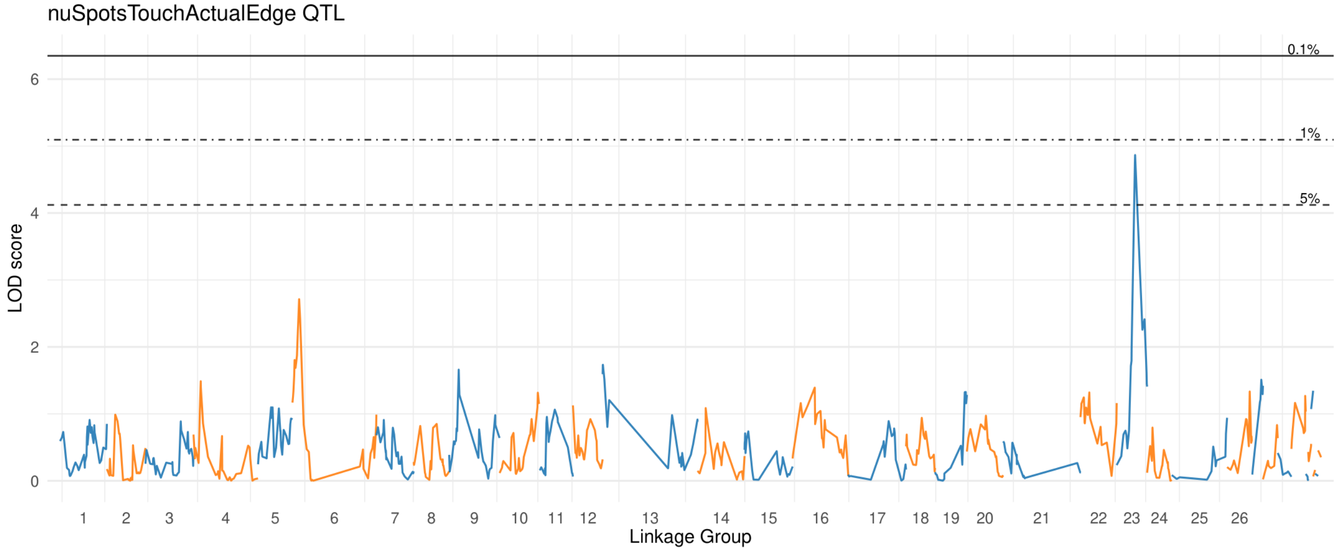


#### Lower Petal

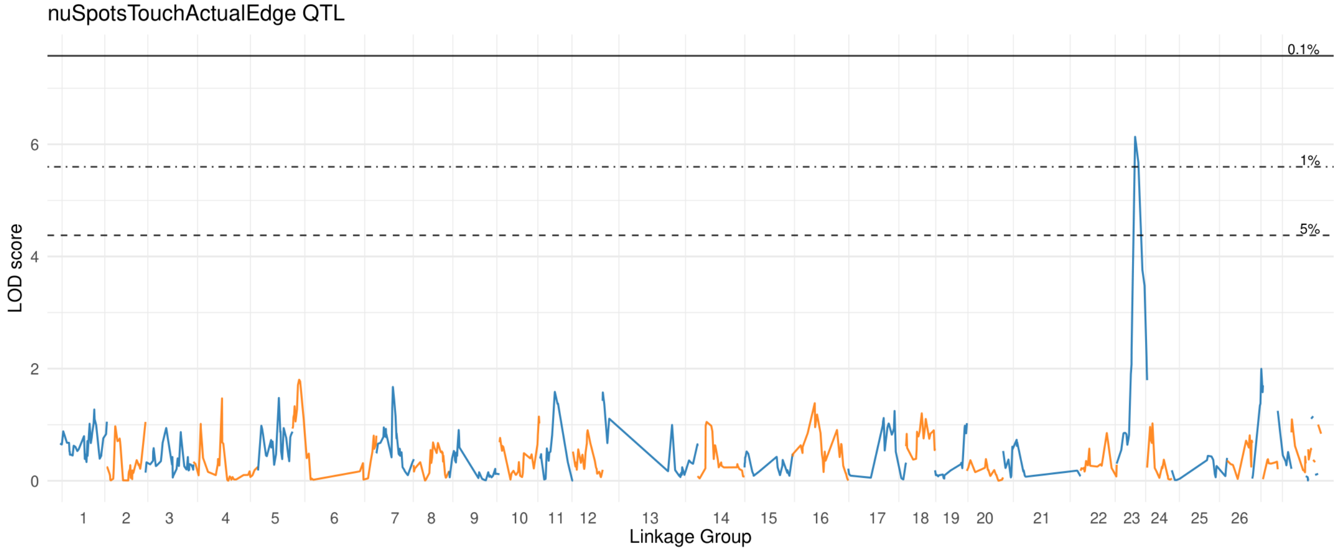

### nu Spots Touch Center

#### Upper Petal

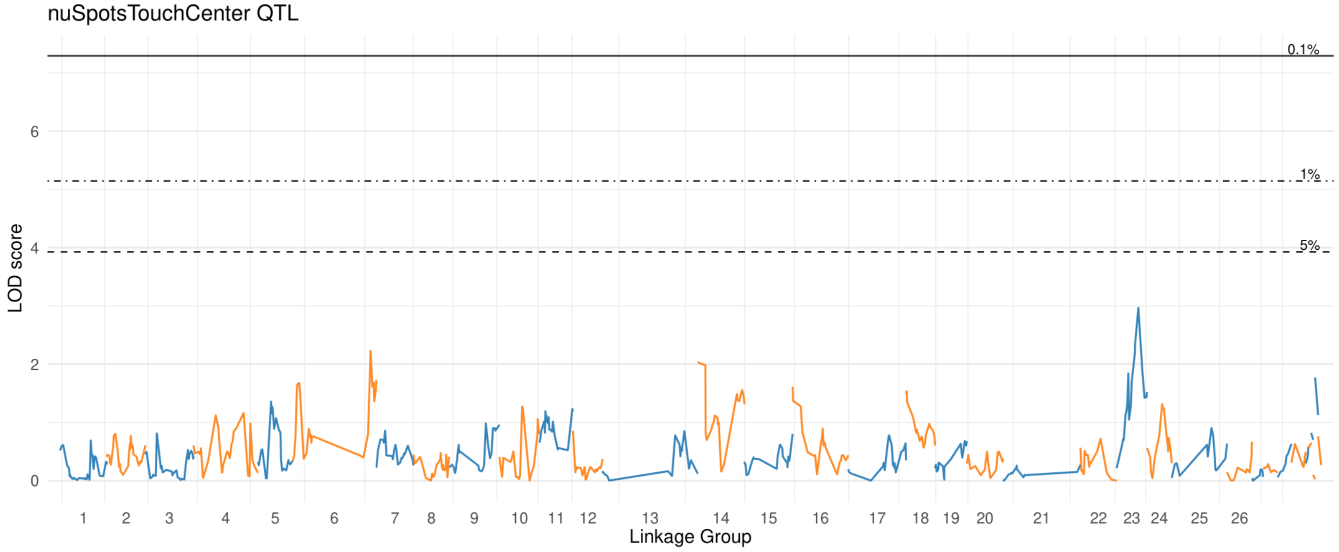


#### Lower Petal

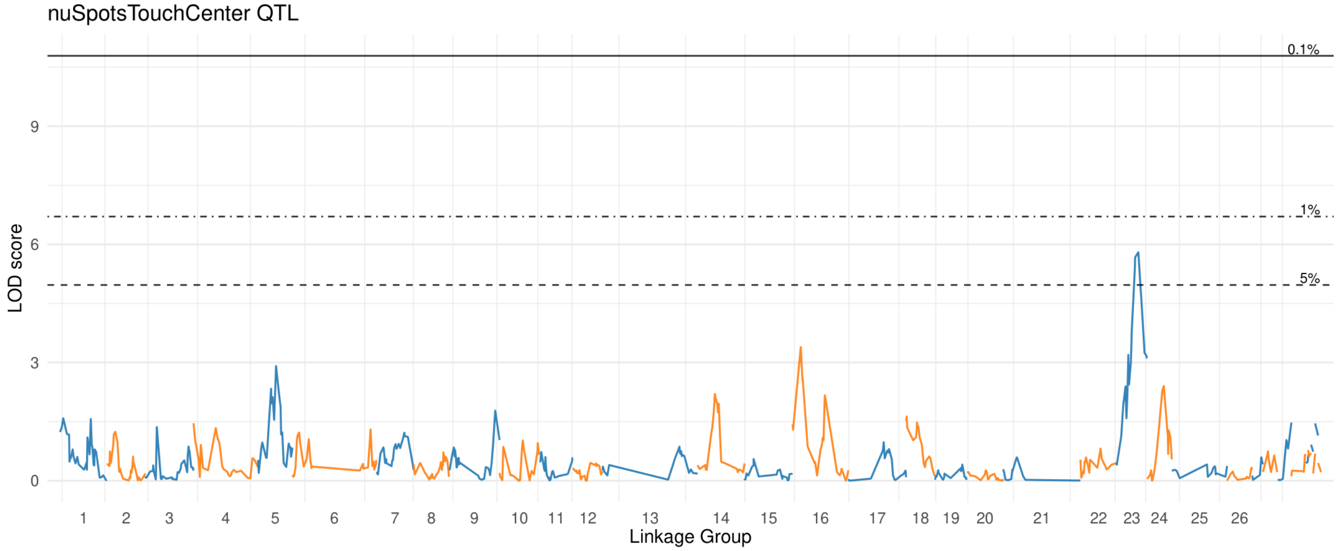

### nu Spots Touch Cut

#### Upper Petal

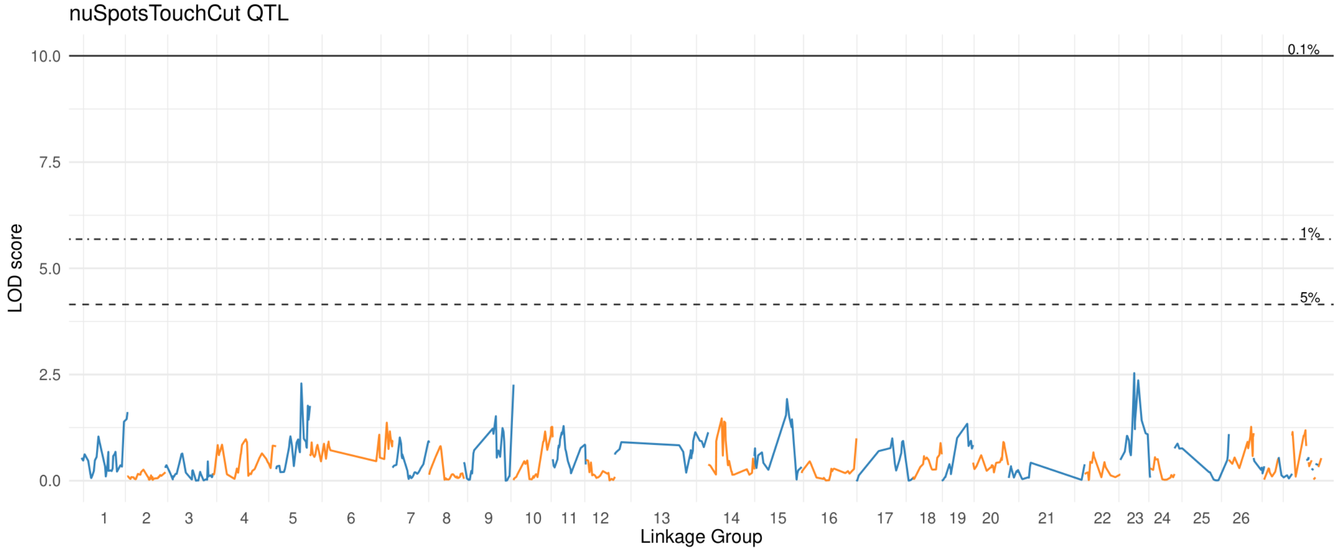


#### Lower Petal

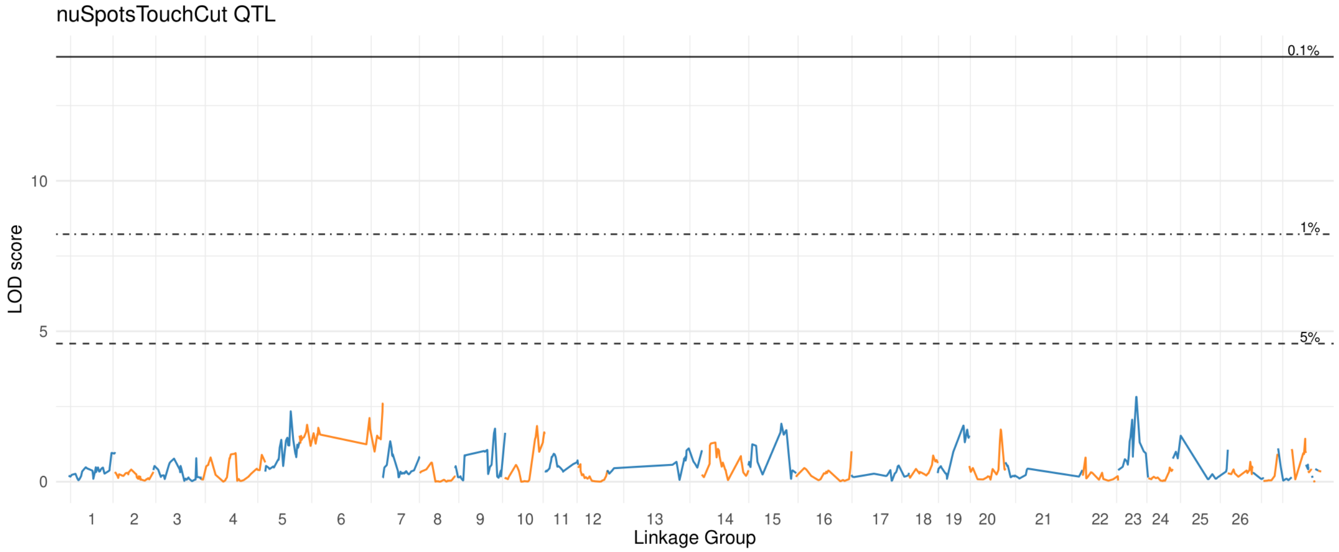

### nu Spots Touch Edge

#### Upper Petal

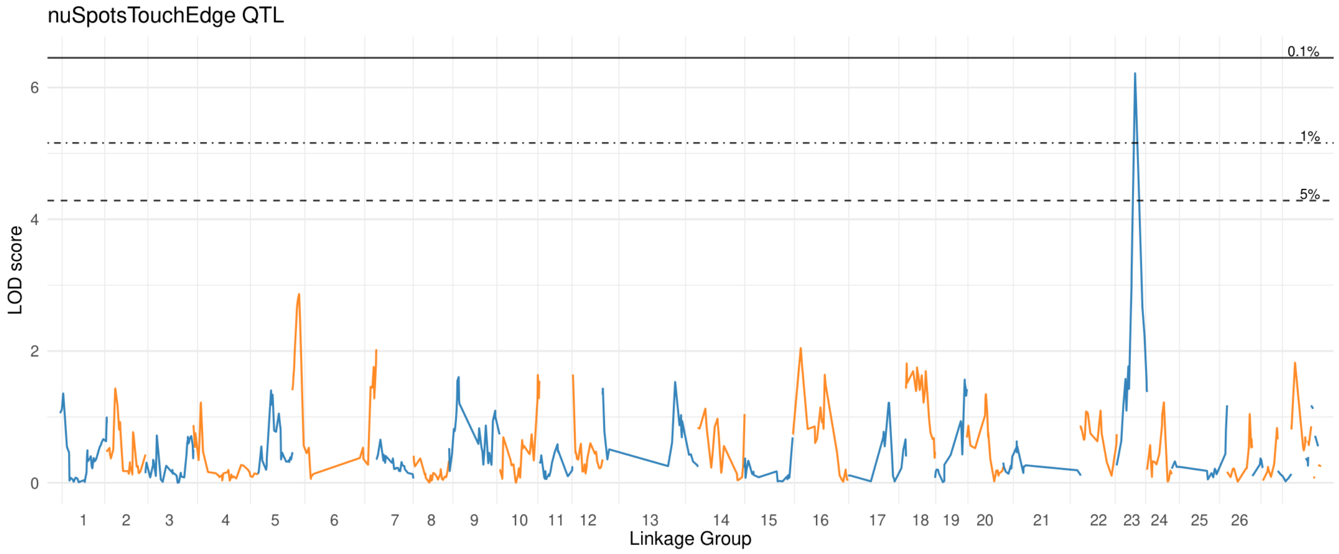


#### Lower Petal

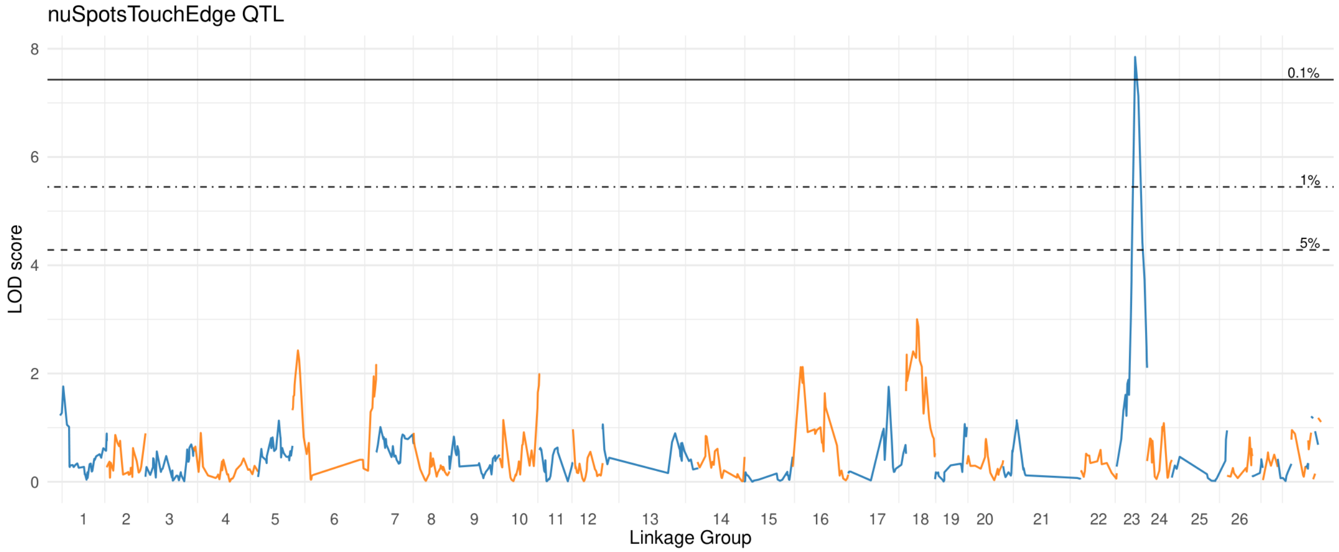

### nu Spots Touch Throat

#### Upper Petal

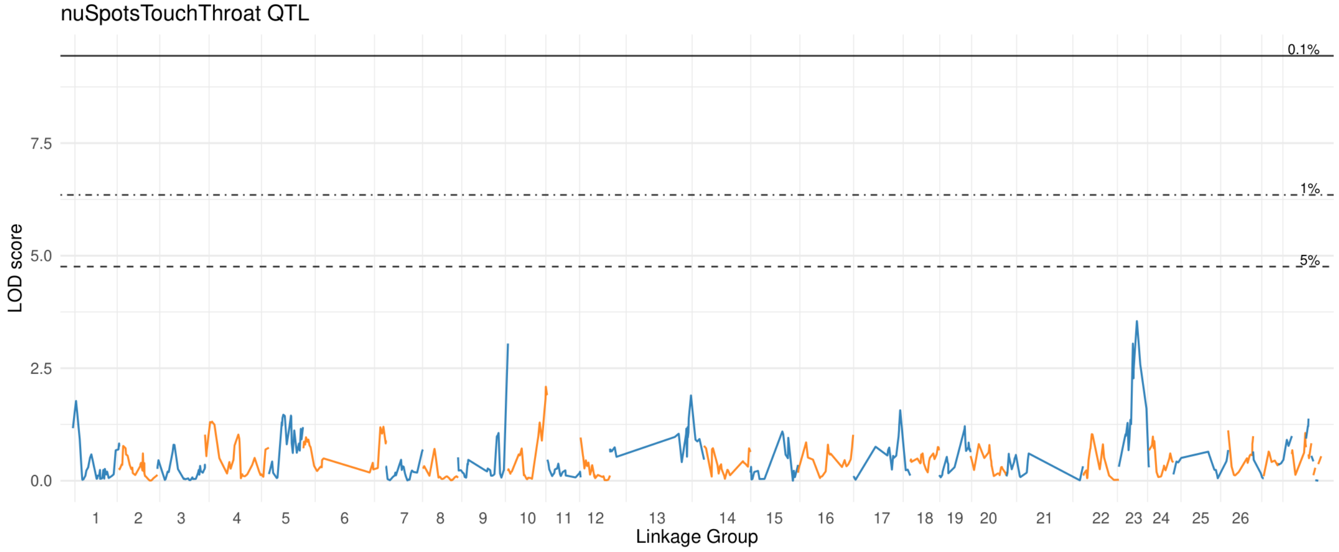


#### Lower Petal

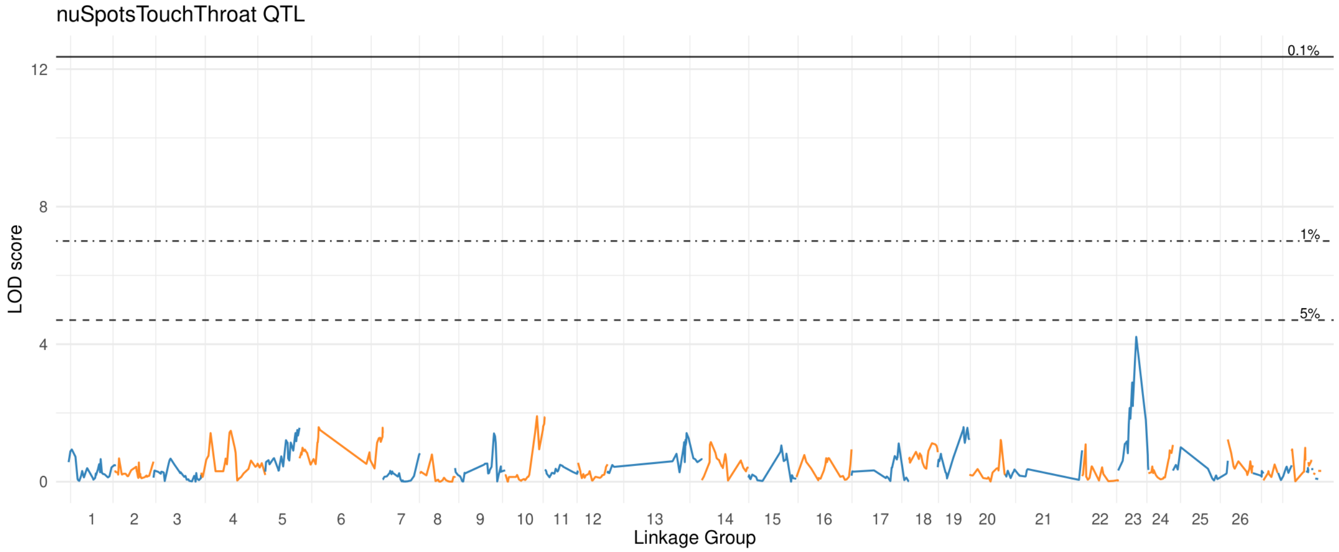

### prop Spots In Center

#### Upper Petal

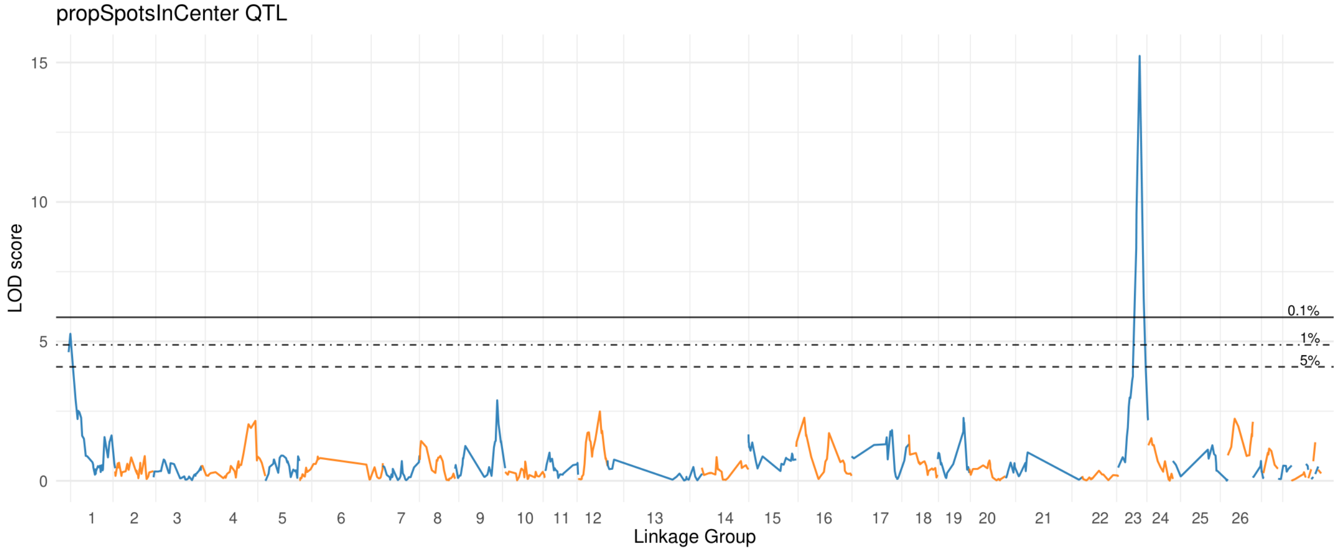


#### Lower Petal

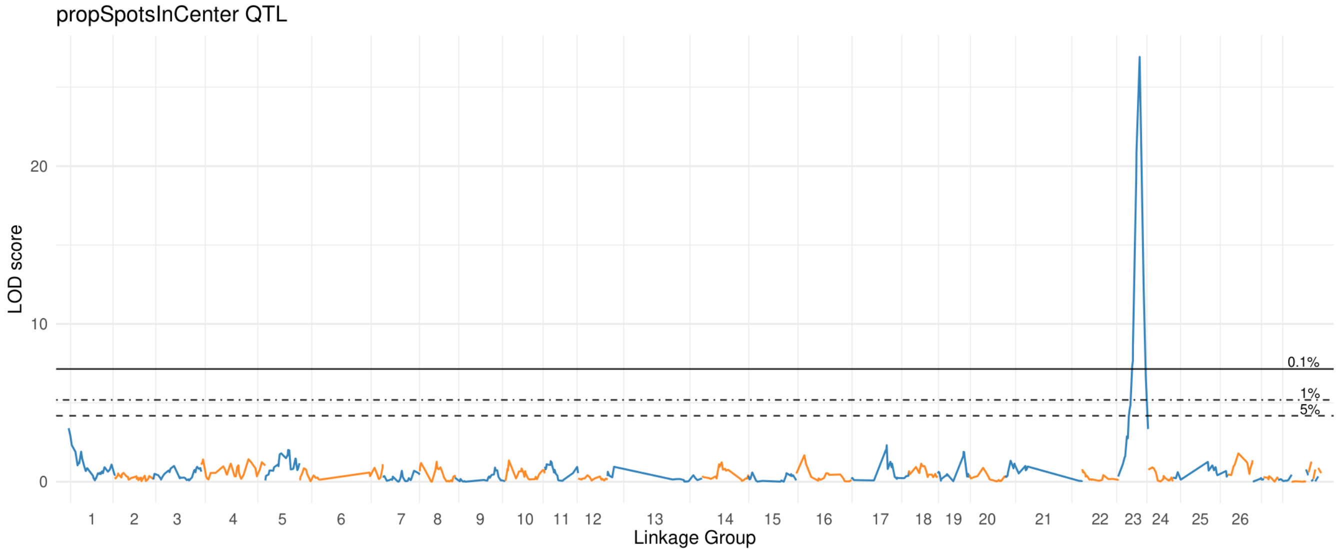

### prop Spots In Dist

#### Upper Petal

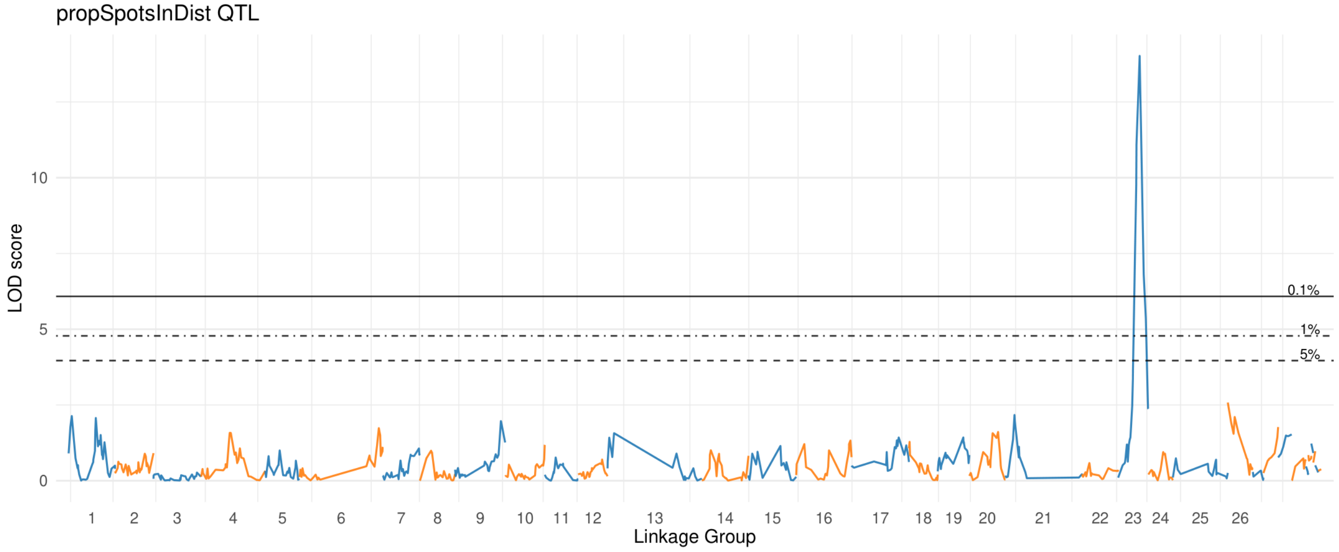


#### Lower Petal

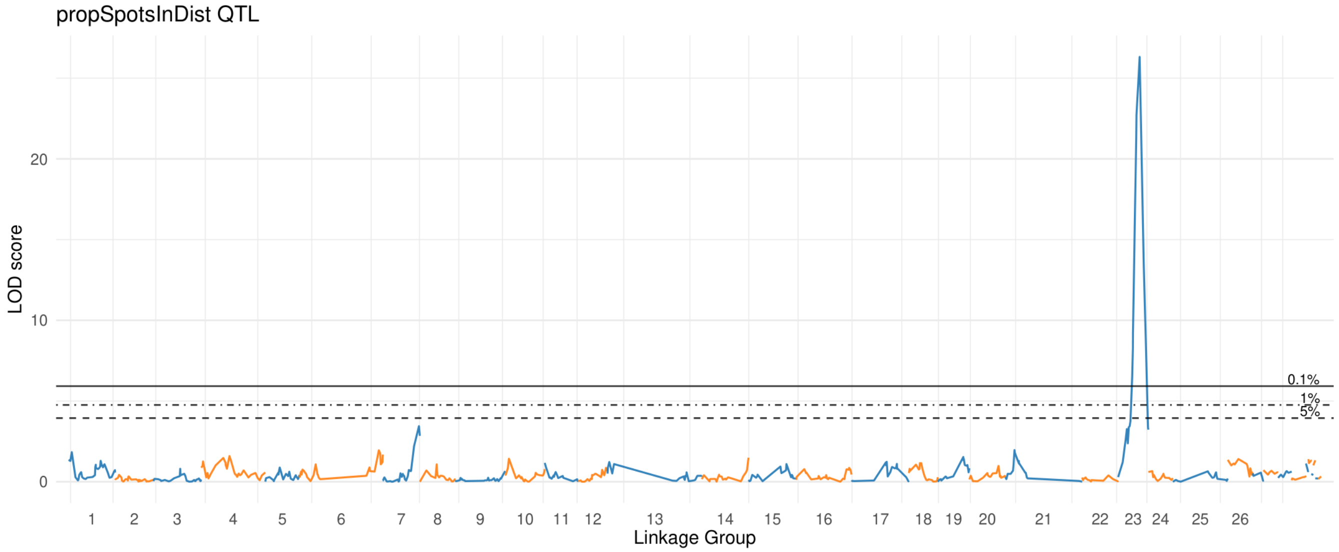

### prop Spots In Edge

#### Upper Petal

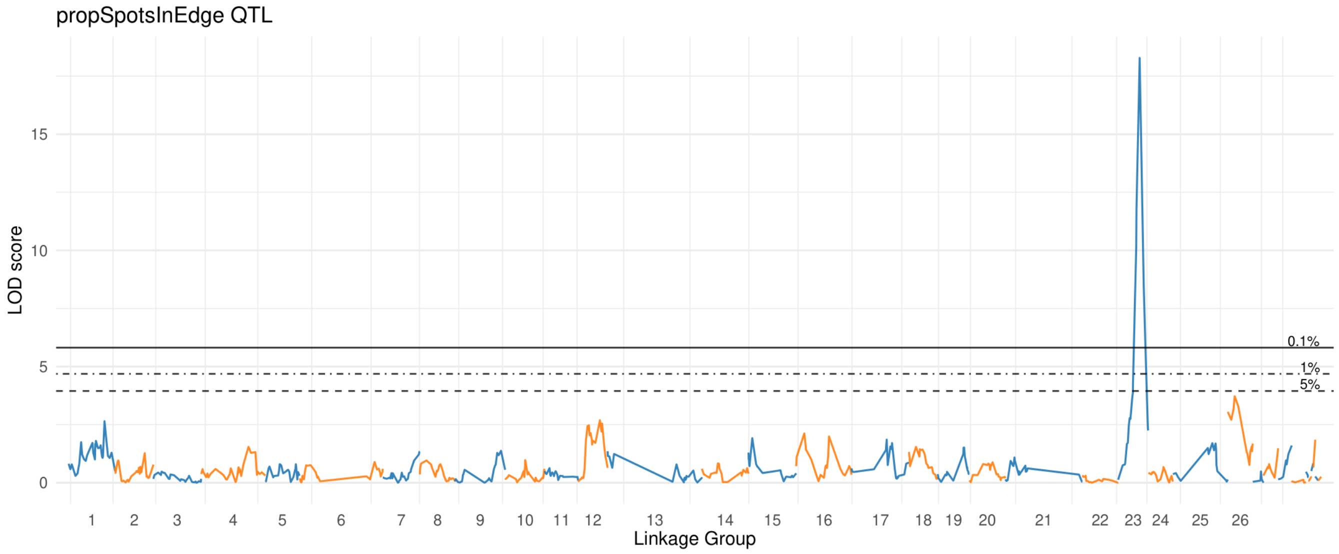


#### Lower Petal

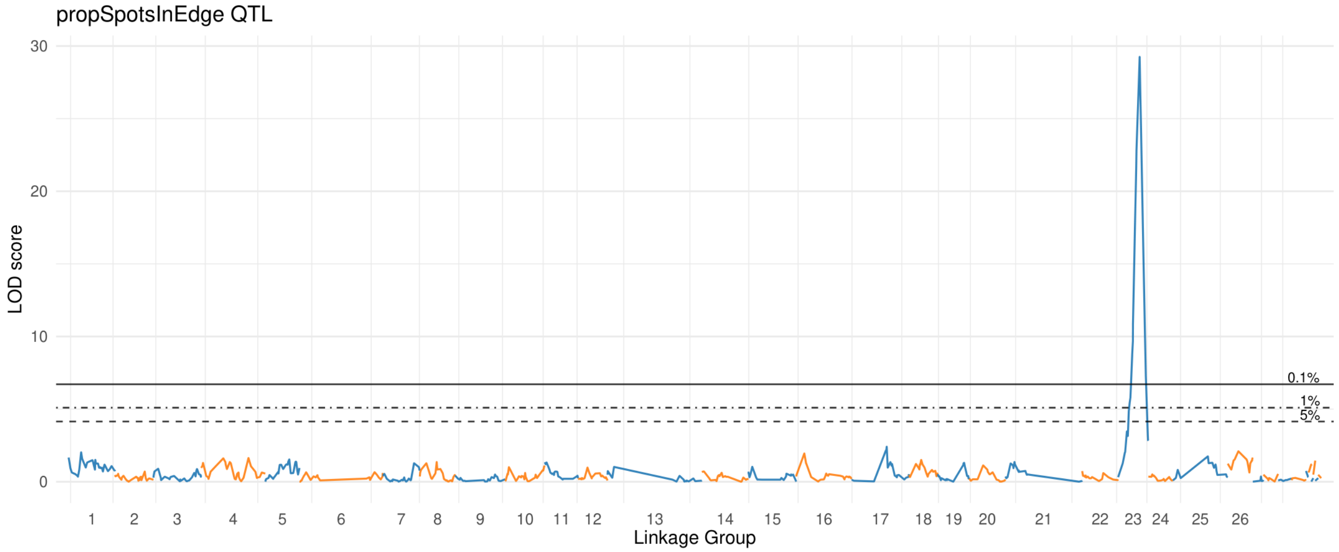

### prop Spots In Prox

#### Upper Petal

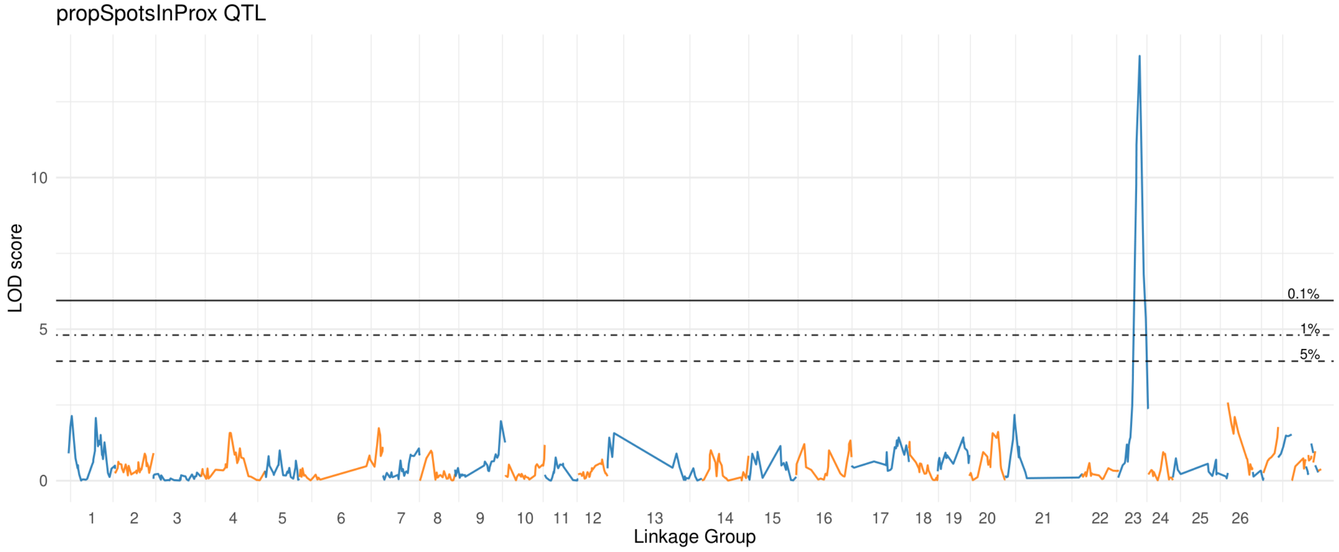


#### Lower Petal

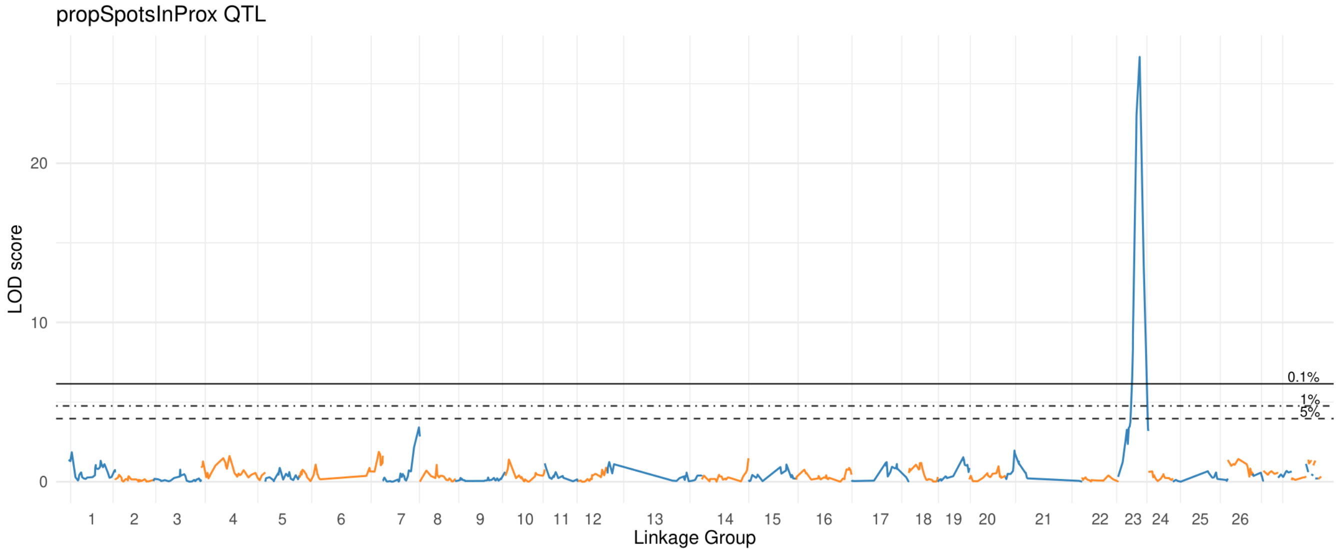

### prop Spots In Quad I

#### Upper Petal

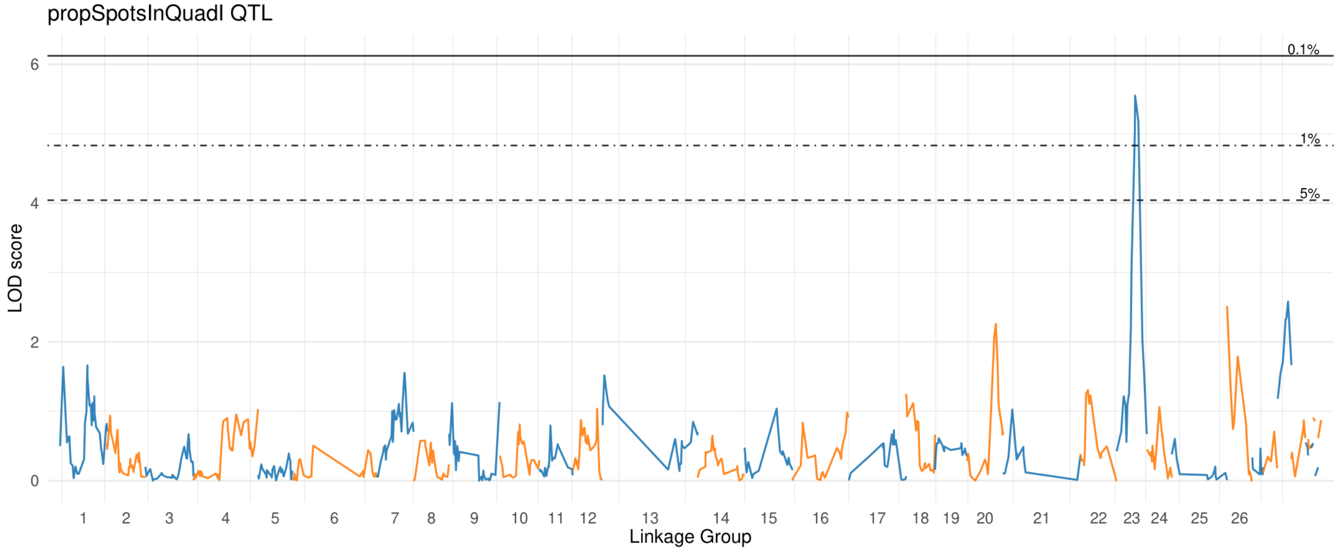


#### Lower Petal

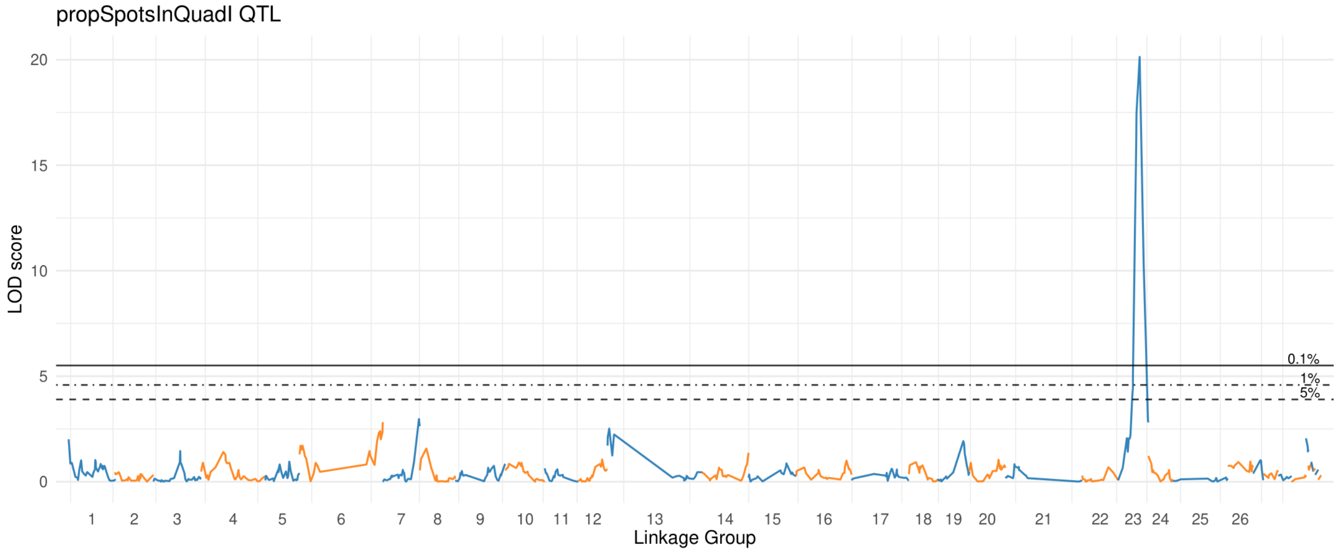

### prop Spots In Quad II

#### Upper Petal

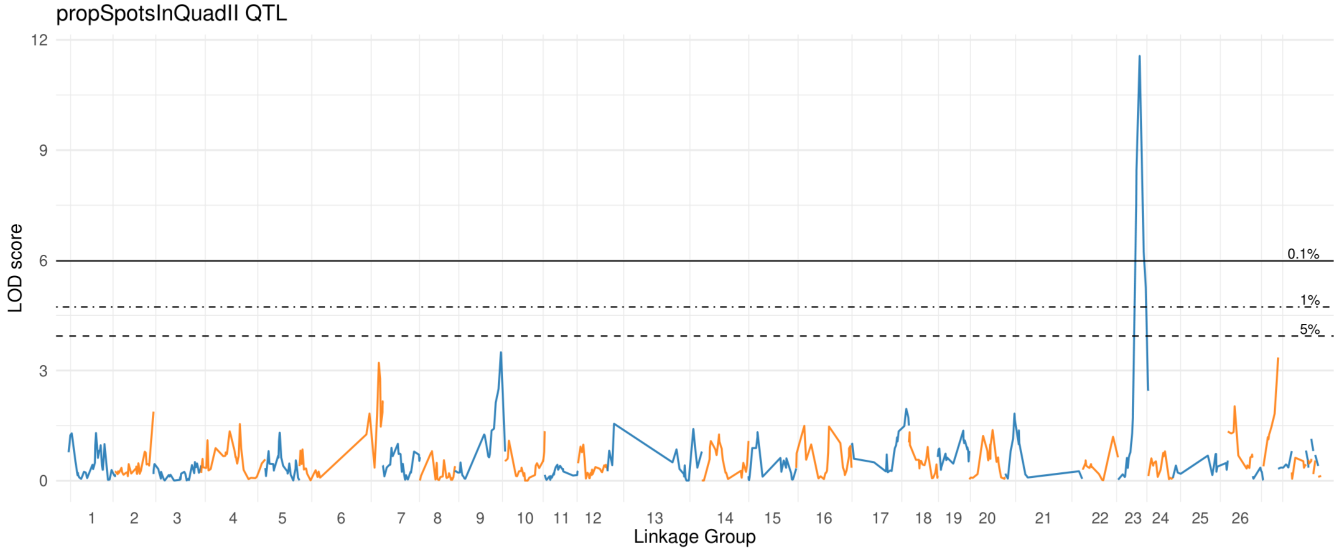


#### Lower Petal

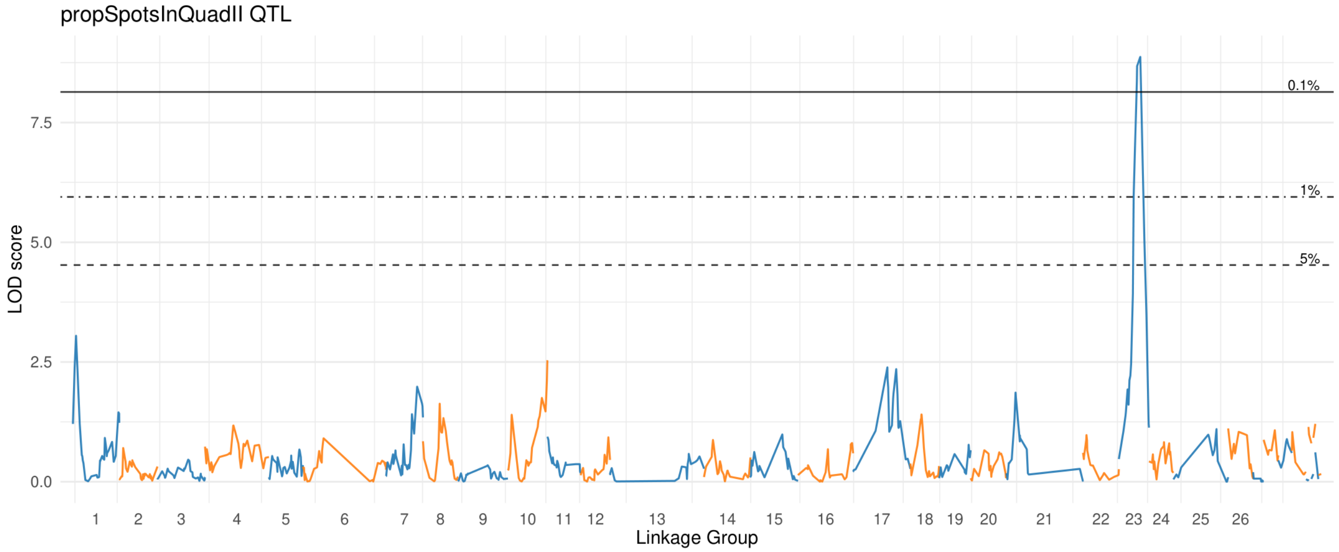

### prop Spots In Quad III

#### Upper Petal

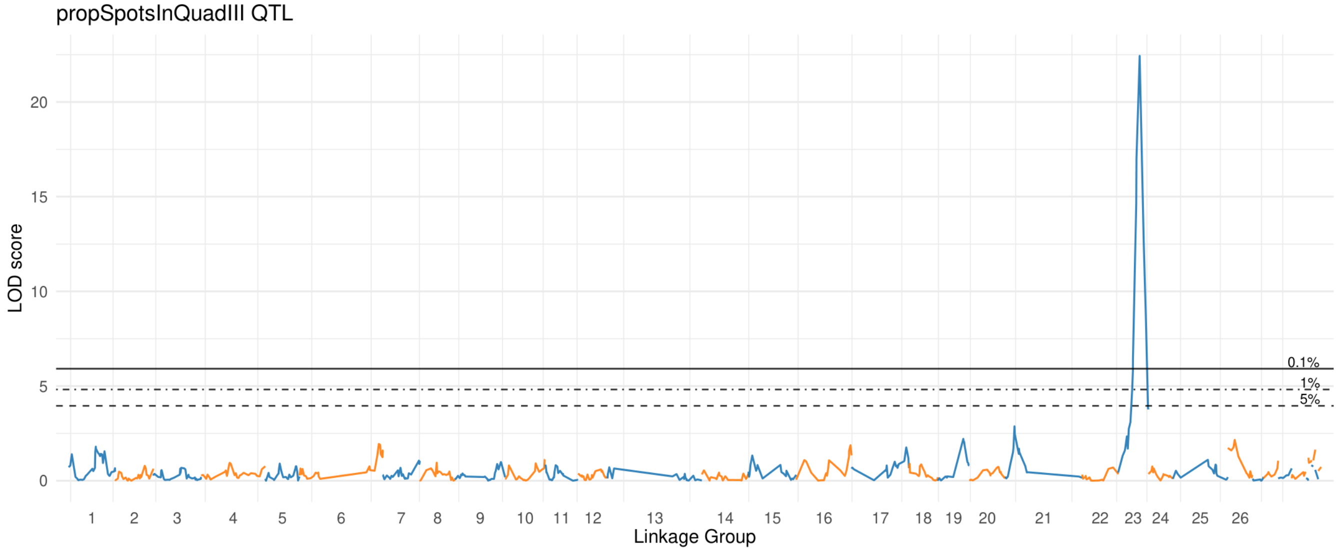


#### Lower Petal

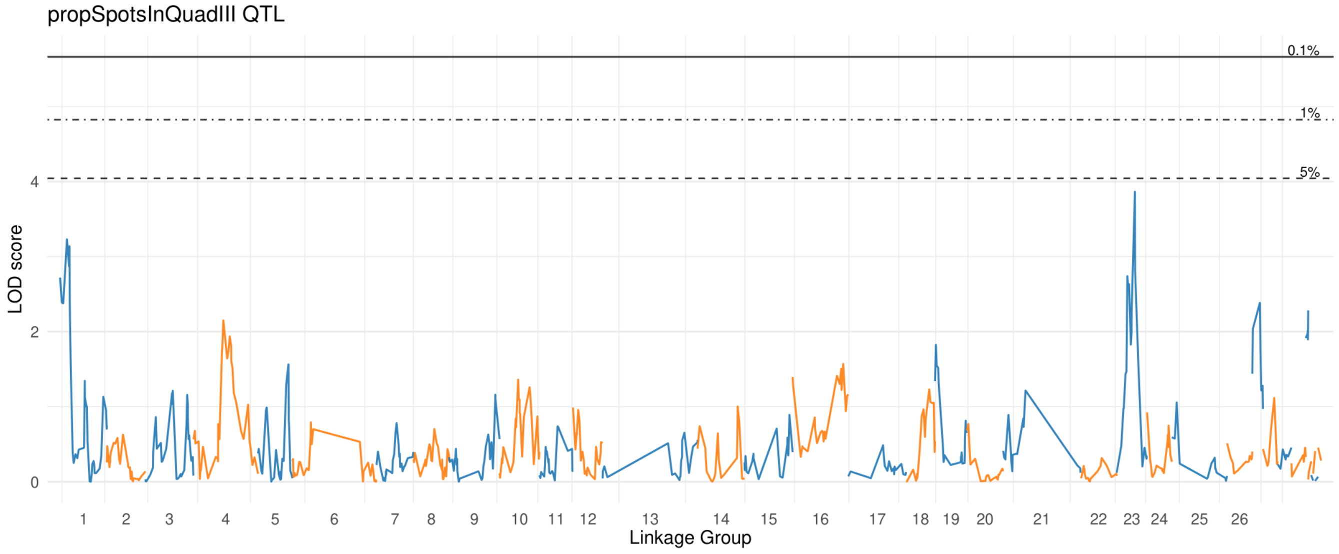

### prop Spots In Quad IV

#### Upper Petal

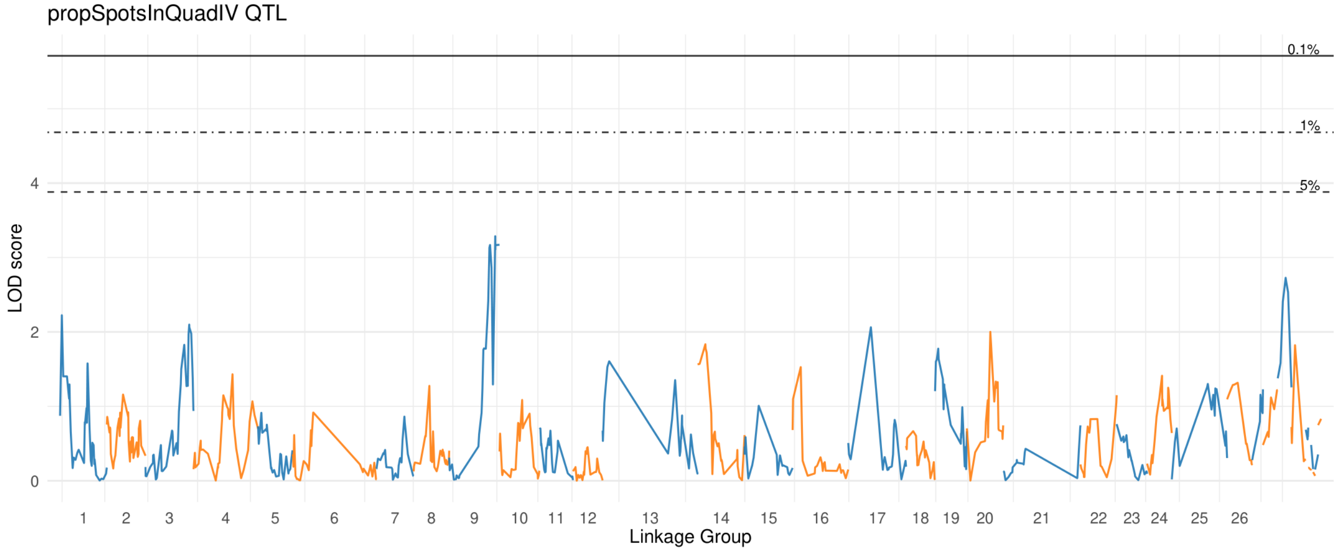


#### Lower Petal

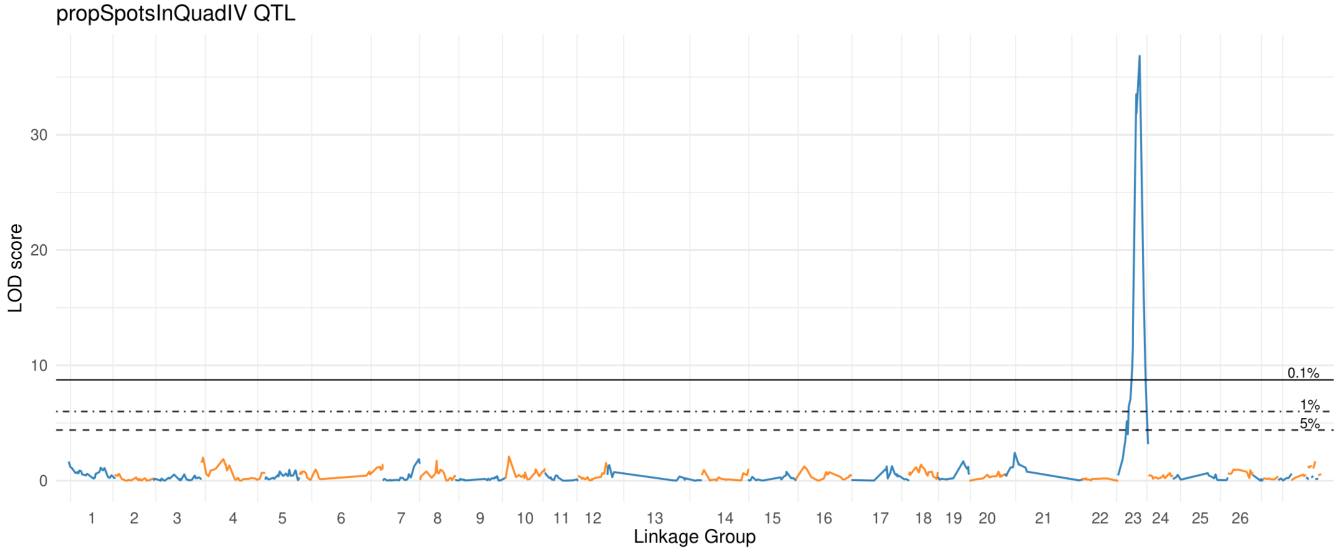

### prop Spots In Throat

#### Upper Petal

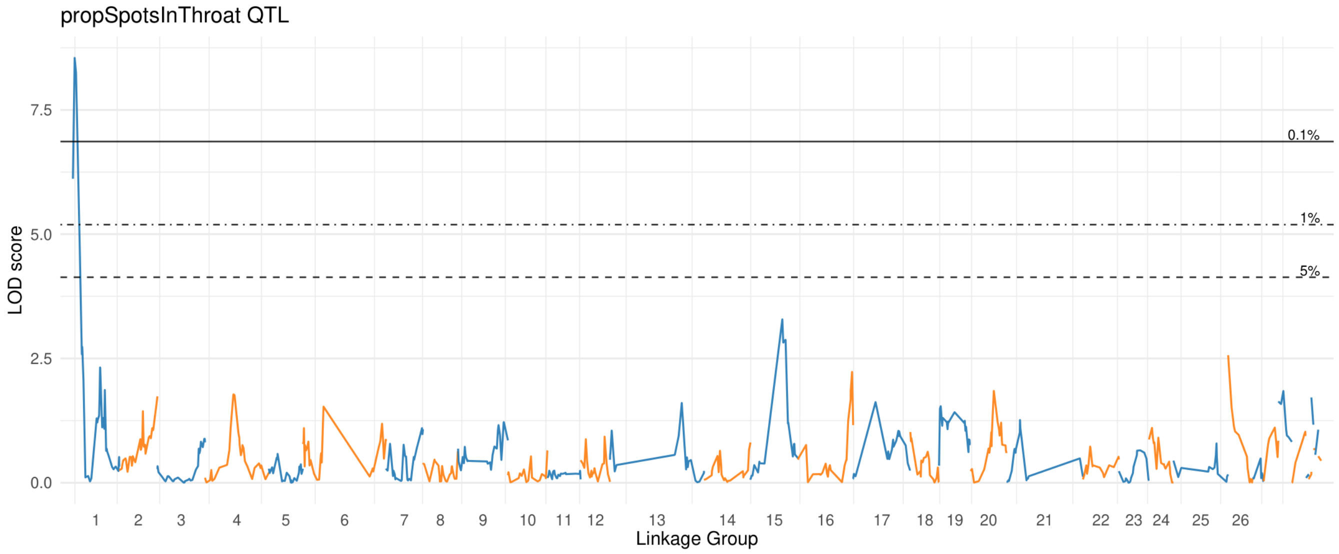


#### Lower Petal

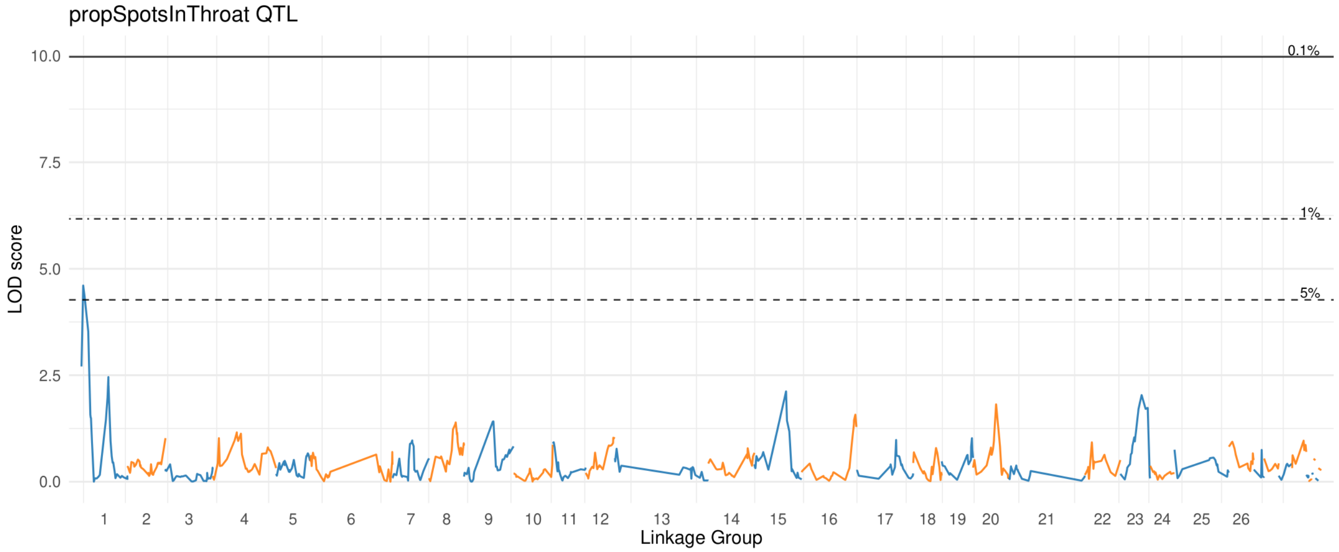

### prox Coveredby Spots

#### Upper Petal

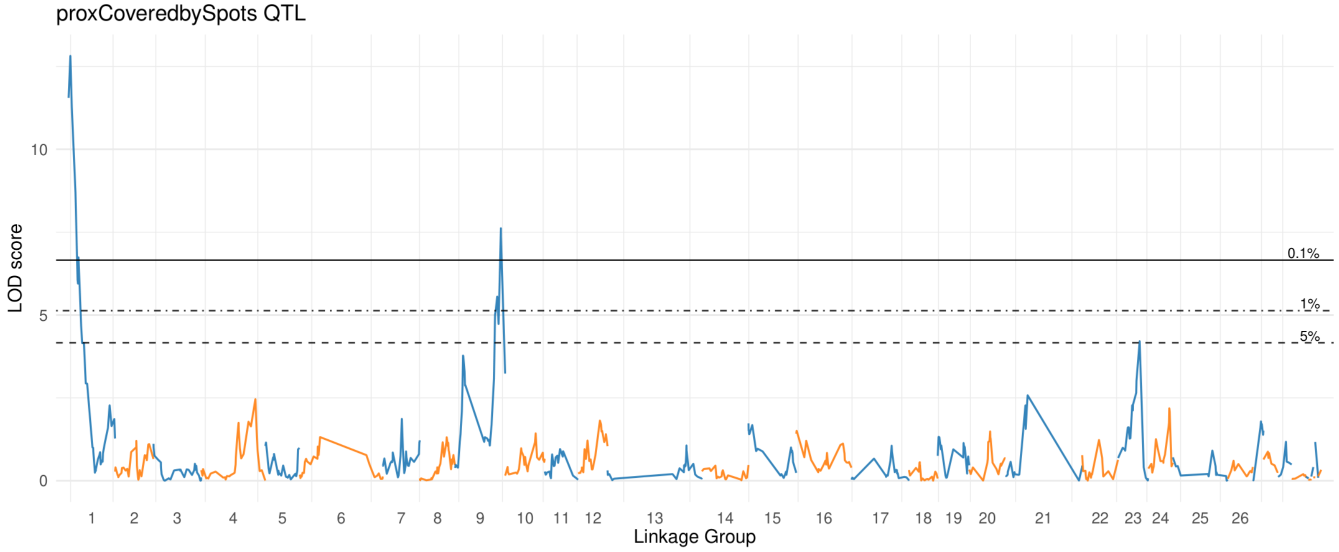


#### Lower Petal

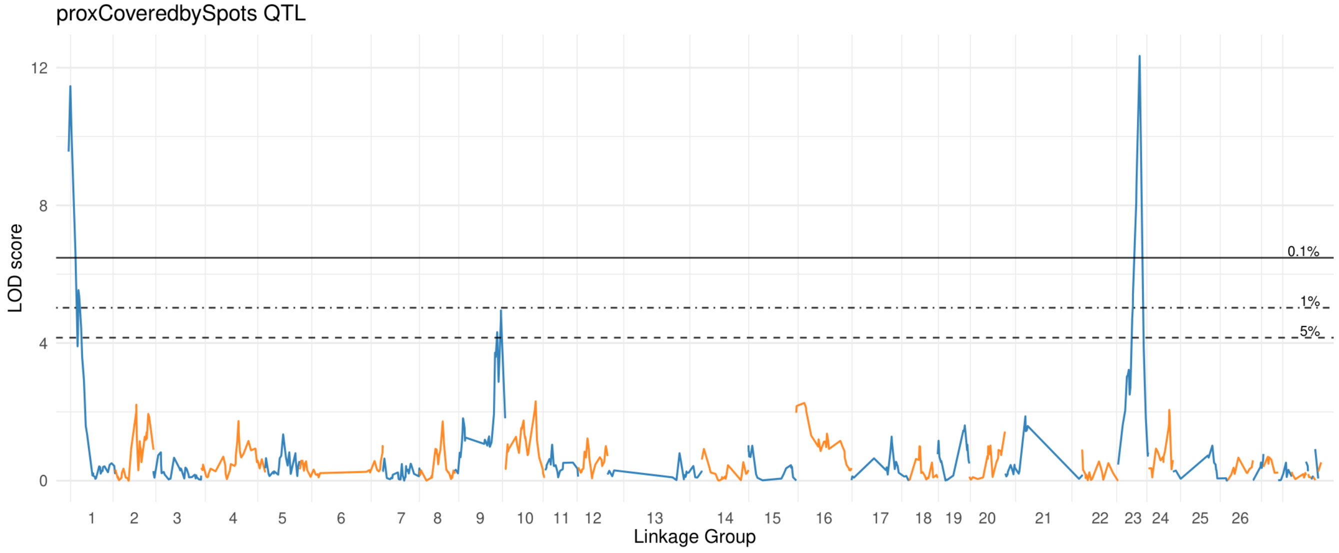

### quad ICoveredby Spots

#### Upper Petal

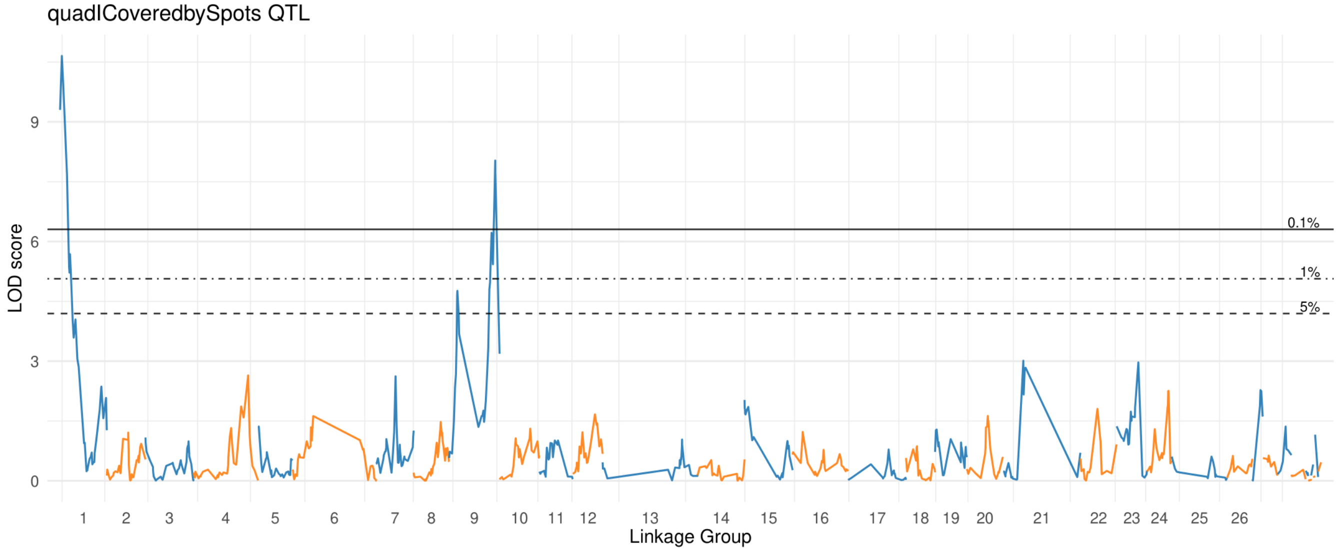


#### Lower Petal

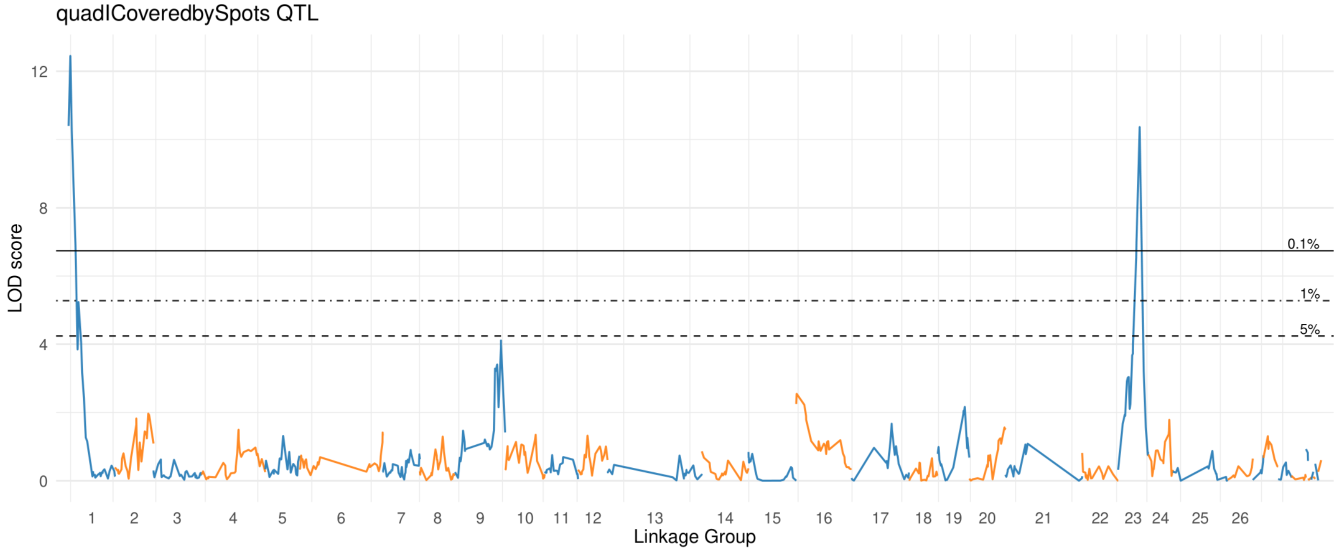

### quad IICoveredby Spots

#### Upper Petal

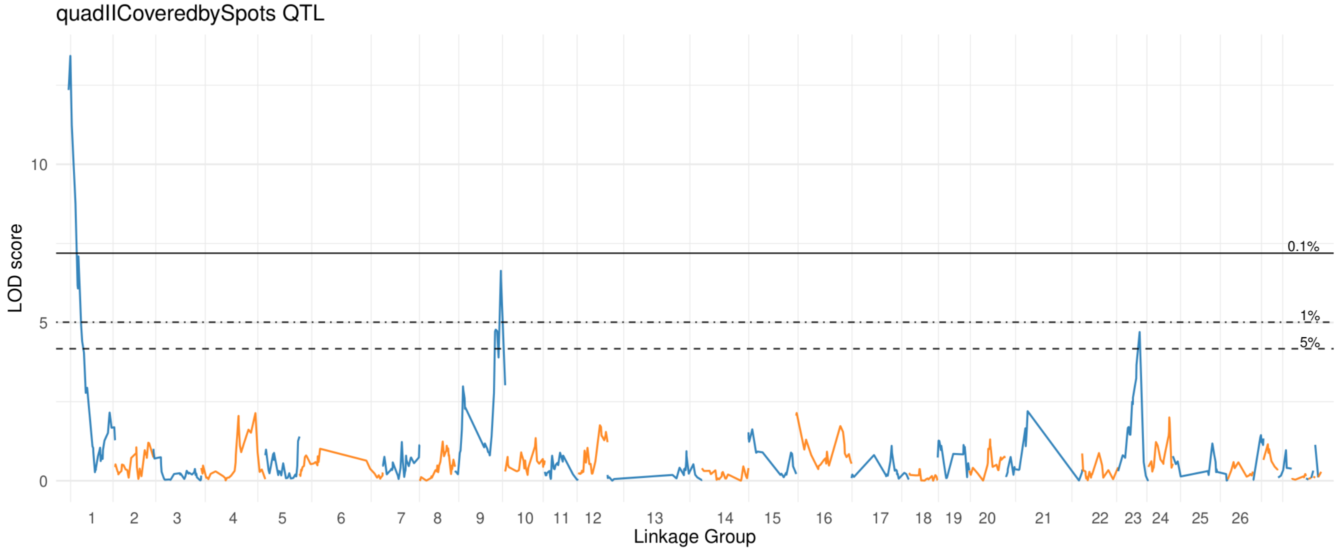


#### Lower Petal

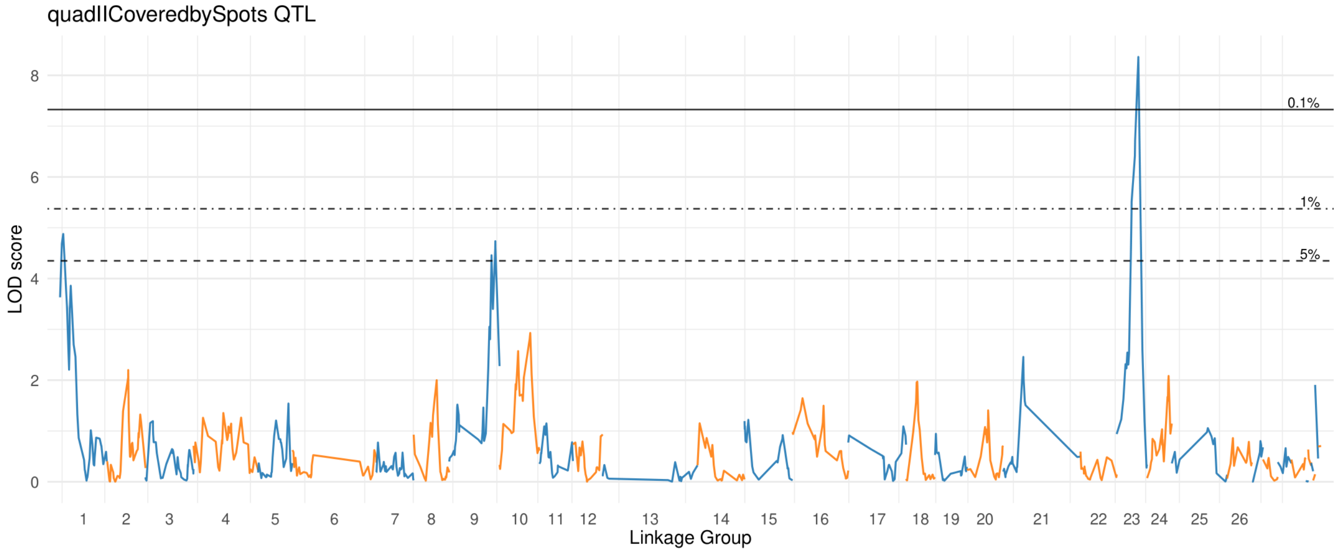

### quad IIICoveredby Spots

#### Upper Petal

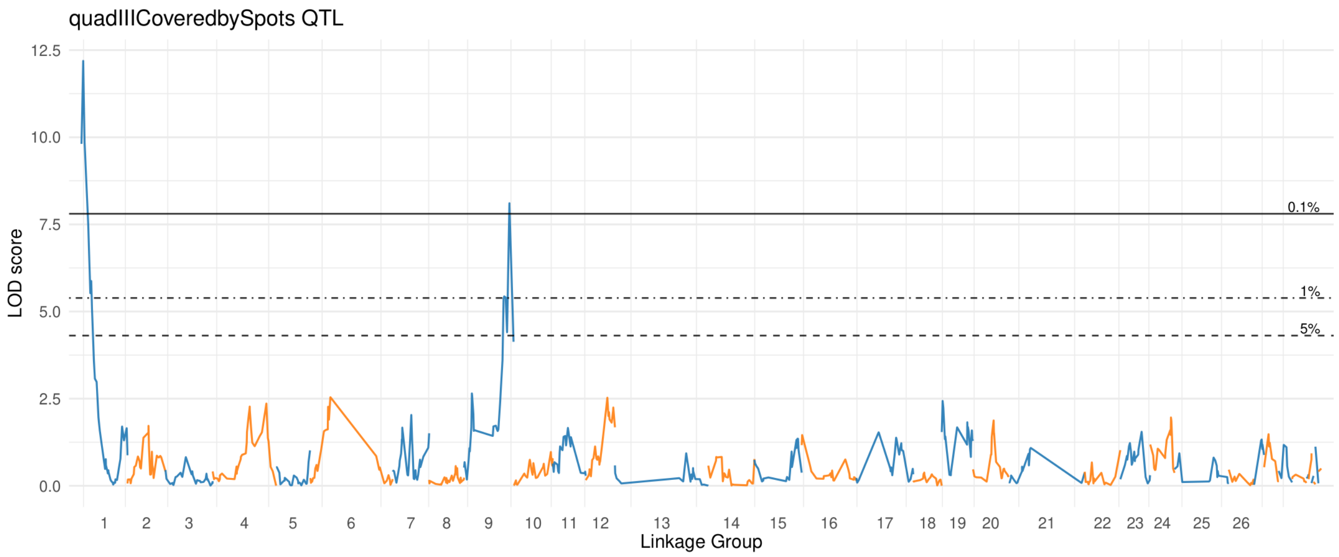


#### Lower Petal

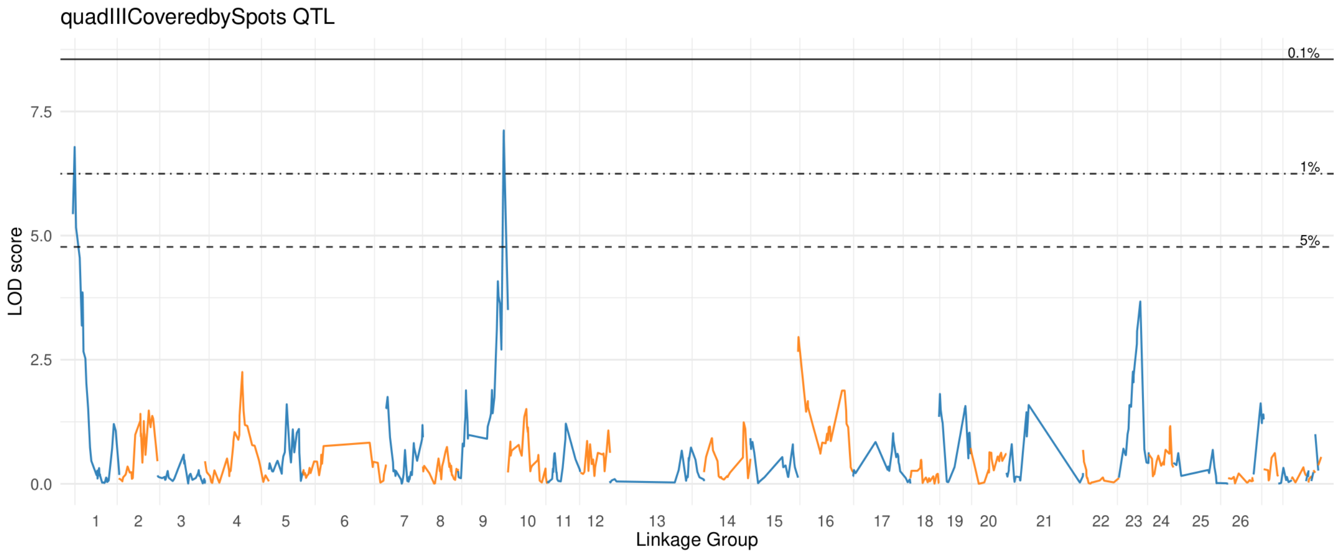

### quad IVCoveredby Spots

#### Upper Petal

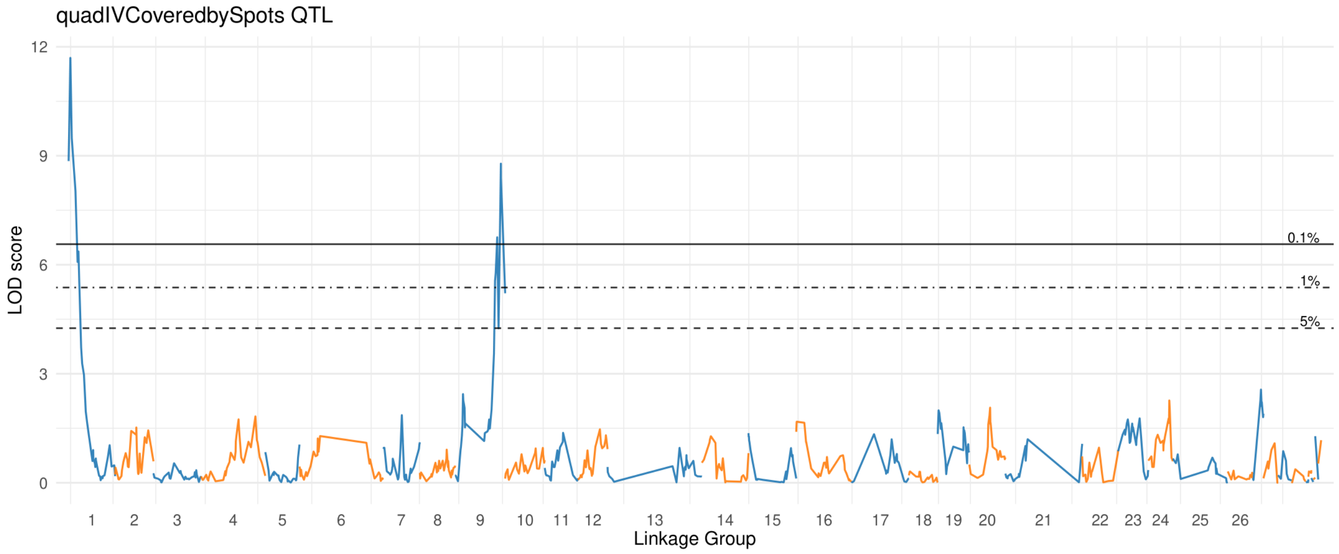


#### Lower Petal

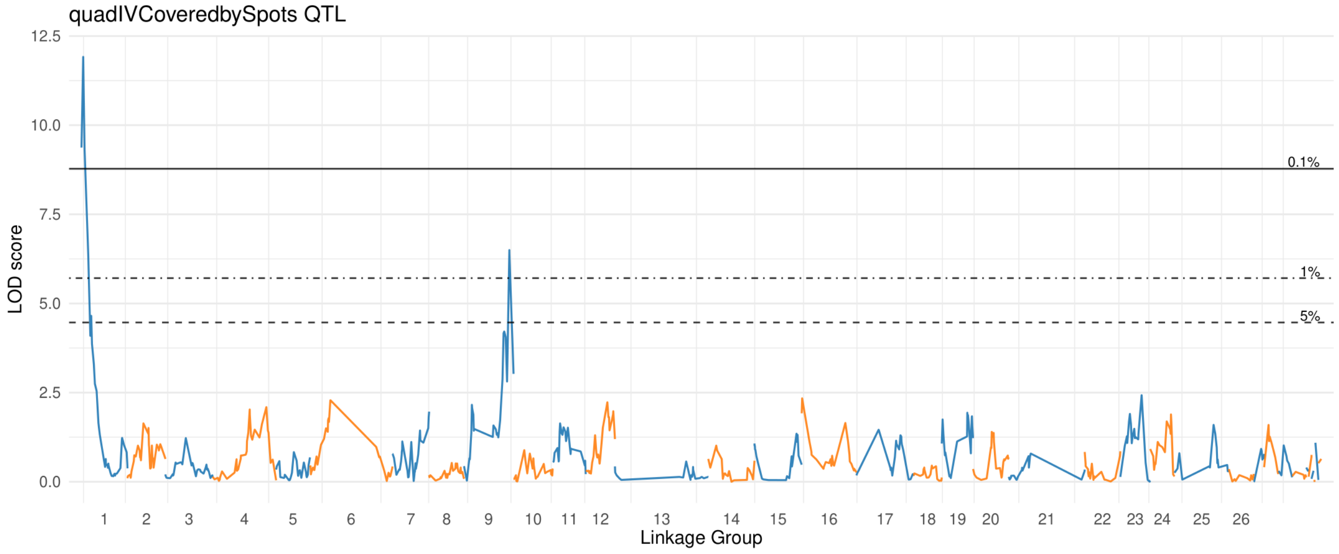

### real Edge Spotted

#### Upper Petal

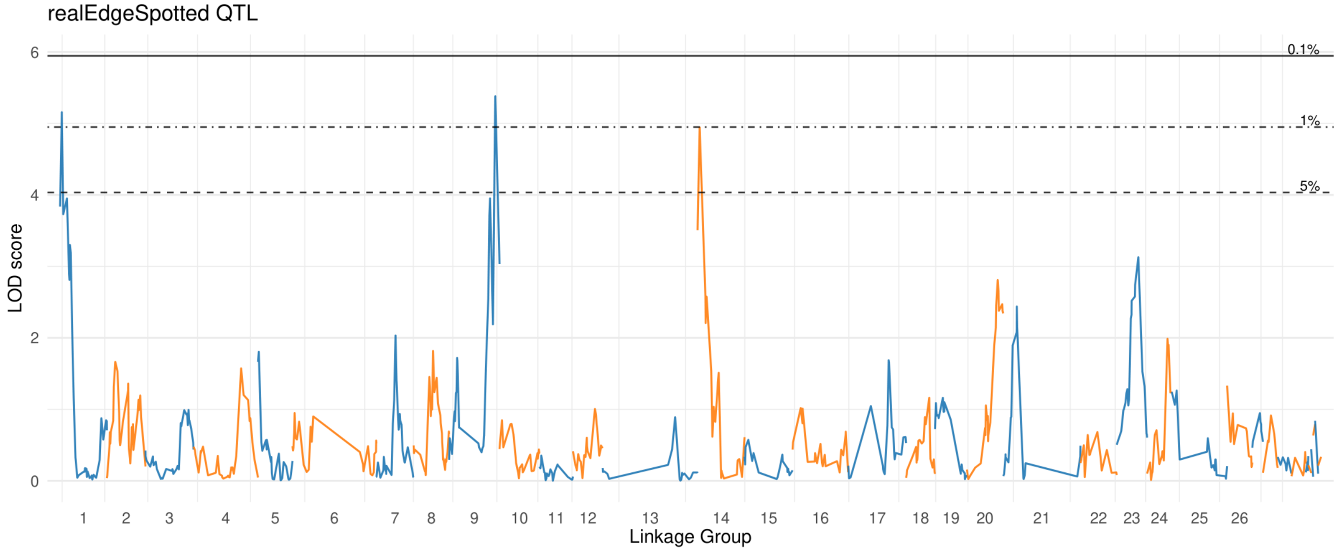


#### Lower Petal

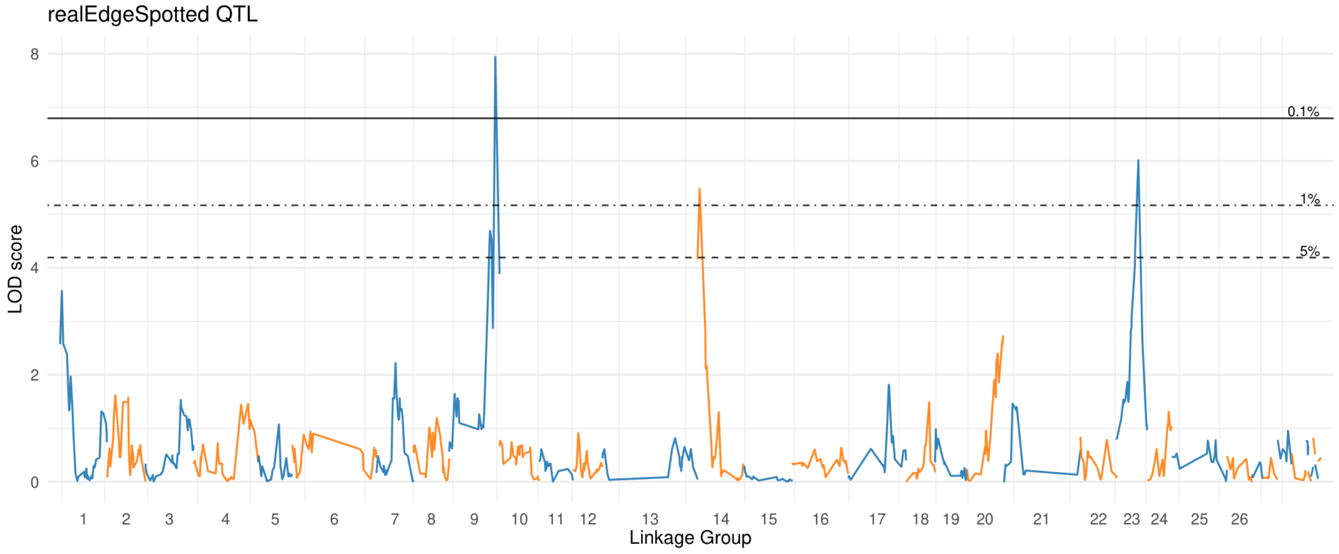

#### smallest Spot Area

##### Upper Petal

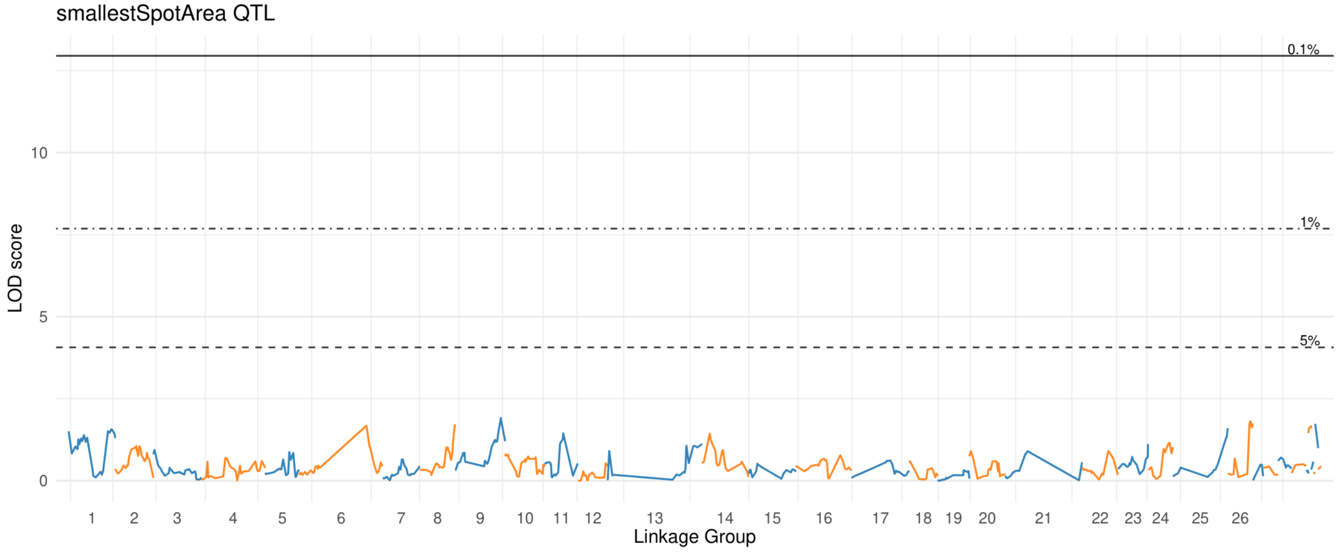


##### Lower Petal

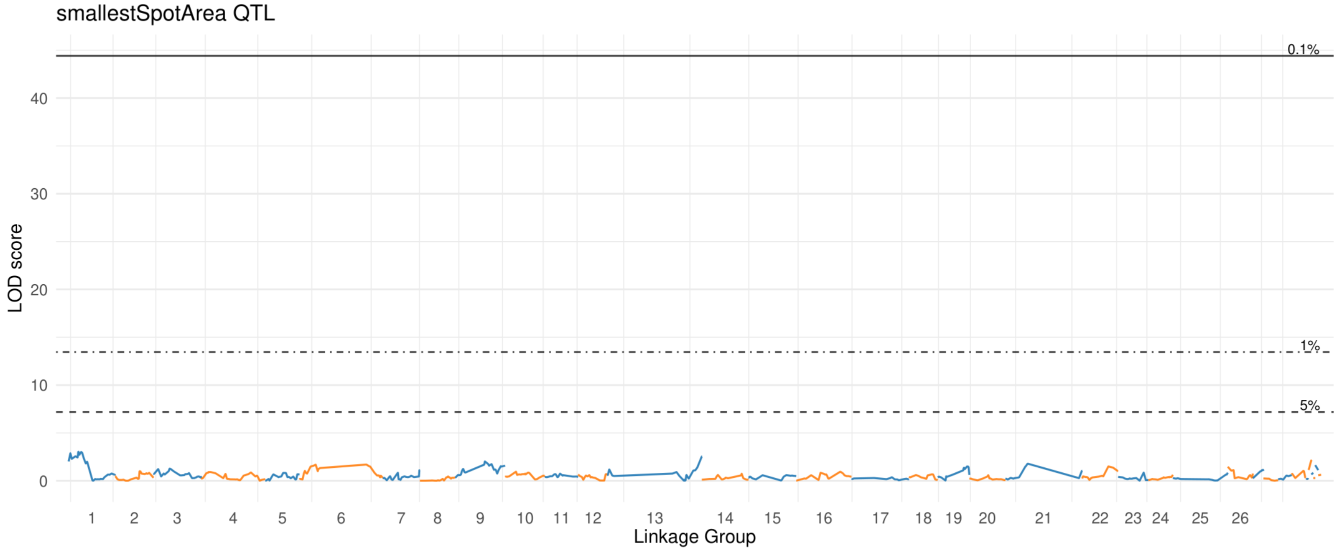

### throat Coveredby Spots

#### Upper Petal

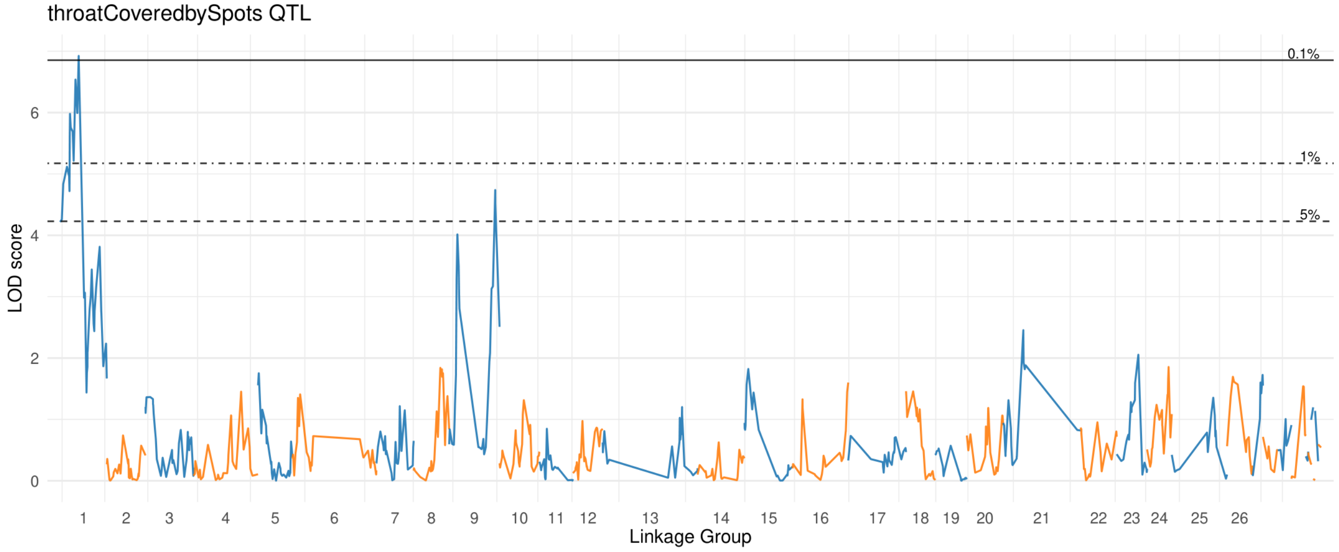


#### Lower Petal

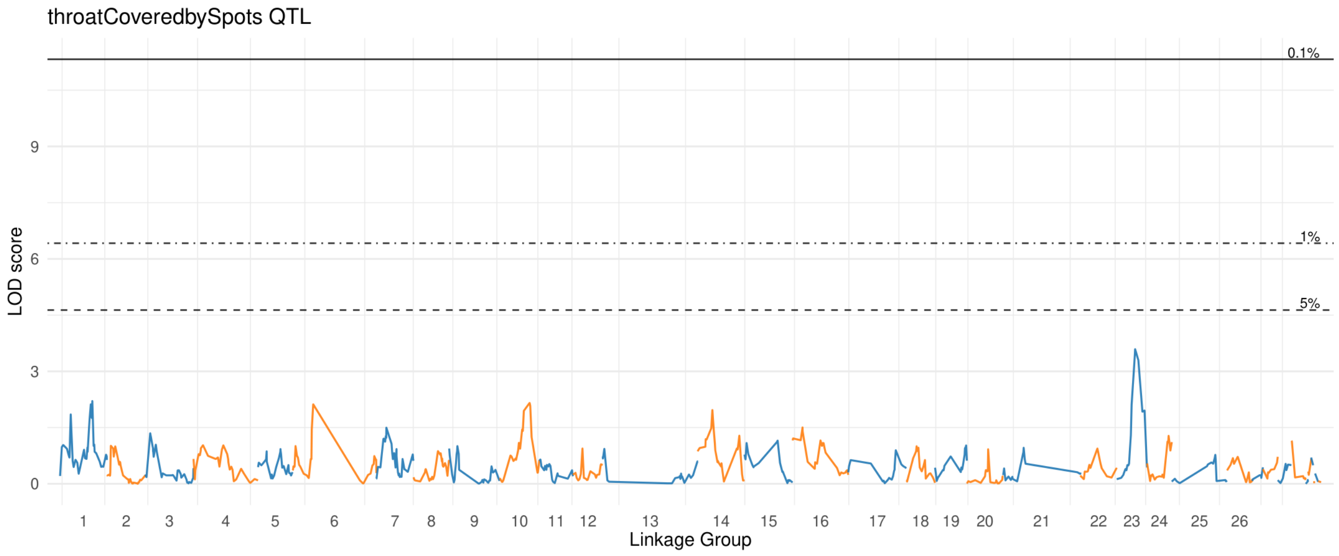
