## Supplemental File 2 for "Genetic architectures of floral pigment and patterning in hybrid monkeyflowers": chatGPTcarotenoid.txt

I have a large tab-delimited file of protein BLAST hits. Each line represents a hit from a predicted protein in my species against Arabidopsis proteins. Example format: query_id arabidopsis_hit_id arabidopsis_annotation percent_identity alignment_length mismatches gap_opens qstart qend sstart send evalue bitscore My biological question is: which predicted genes are the best candidates for involvement in carotenoid production? If the file is large, first scan all rows, group hits by query ID, and identify the most informative hits per query before interpreting the results. Do not base the interpretation on the first few rows only. I am interested in genes involved in carotenoid biosynthesis, carotenoid precursor production, carotenoid pathway regulation, and carotenoid metabolic modification or degradation. I am not asking about the cellular or tissue-level location of carotenoids, pigment storage, or pigment patterning unless those genes also have a clear role in carotenoid production. Relevant classes of genes include, but are not limited to: 1. Core carotenoid biosynthesis genes Examples: PSY, PDS, ZDS, ZISO, CRTISO, LCYB, LCYE, BCH, CYP97A, CYP97B, CYP97C, LUT1, LUT2, VDE, ZEP, NXS, NSY, and other enzymes directly involved in producing carotenoids or xanthophylls. 2. Upstream isoprenoid / plastidial MEP pathway genes that supply carotenoid precursors Examples: DXS, DXR, MCT, CMK, MDS, HDS, HDR, IDI, GGPPS, and related enzymes involved in producing IPP, DMAPP, or GGPP for carotenoid biosynthesis. 3. Carotenoid pathway regulators Examples: transcription factors, plastid development regulators, light-signaling genes, phytochrome-related regulators, PIFs, HY5, GLKs, ORANGE/OR, and other genes known or plausibly implicated in regulating carotenoid biosynthetic gene expression or carotenoid accumulation through pathway control. 4. Carotenoid modification, turnover, or cleavage genes Examples: CCDs, NCEDs, carotenoid cleavage dioxygenases, ABA-biosynthesis-related carotenoid cleavage enzymes, and genes involved in xanthophyll cycle reactions. 5. Indirect but plausible carotenoid production modifiers Examples: genes involved in chloroplast/plastid development, photosynthetic apparatus development, tetrapyrrole/chlorophyll-related plastid status signaling, redox regulation, light response, or precursor availability, but only when the annotation and BLAST quality make the connection biologically plausible. Please analyze the BLAST results and return only the strongest and most interpretable candidates, not every speculative idea. For each predicted gene/query, consolidate multiple BLAST hits if they appear to point to the same biological function. Do not treat every hit independently if they are redundant. Please produce the following output: 1. A ranked candidate table with columns: - Rank - Query ID - Best Arabidopsis hit or hits - Putative functional category Use one of: Core carotenoid biosynthesis Upstream isoprenoid / MEP precursor biosynthesis Carotenoid pathway regulation Carotenoid modification / cleavage / turnover Indirect but plausible carotenoid production modifier Probably not relevant - Why this gene is relevant - Evidence strength: High / Medium / Low - Main BLAST evidence: percent identity, alignment length, E-value, bitscore - Pressure-test / caveat: why this could be a false lead - Suggested follow-up validation 2. A short “best bets” section: Identify the top 5–15 genes that are most worth following up, with concise reasoning. 3. A “do not overinterpret” section: List genes or categories that may sound relevant but should be treated cautiously, especially if the hit is short, low identity, generic, or belongs to a large nonspecific gene family. 4. A “missing expected genes” section: Note whether obvious carotenoid pathway genes or regulators appear absent from the BLAST hits. Do not assume absence from the genome, only absence from this BLAST result file. Important analysis rules: - Prioritize biological relevance plus BLAST quality. A gene with a perfect functional annotation but weak BLAST support should not be ranked as high confidence. - Be skeptical of very short alignments, generic annotations, large gene families, and “protein of unknown function” hits. - Do not invent functions beyond what is supported by the Arabidopsis annotation and BLAST quality. - Distinguish direct carotenoid biosynthesis genes from upstream precursor genes, regulators, and indirect plastid-related modifiers. - Do not include genes only because they affect pigment localization, storage, visibility, or tissue patterning. The focus is carotenoid production. - For regulatory genes, prioritize those with known or plausible roles in carotenoid biosynthesis, plastid development, light regulation, or precursor supply. - If a gene is relevant only indirectly, say so clearly. - Use cautious language: “candidate,” “possible,” “weakly supported,” “requires validation.” - Return a concise, prioritized interpretation rather than a comprehensive list. Give me results as an excel file and named carotenoid.xlsx
