## Supplemental File 2 for "Genetic architectures of floral pigment and patterning in hybrid monkeyflowers": chatGPTspotprompt.txt

I have a large tab-delimited file of protein BLAST hits. Each line represents a hit from a predicted protein in my species against Arabidopsis proteins. Example format:

query_id   arabidopsis_hit_id   arabidopsis_annotation   percent_identity   alignment_length   mismatches   gap_opens   qstart   qend   sstart   send   evalue   bitscore

My biological question is: which predicted genes are the best candidates for involvement in anthocyanin pigmentation and anthocyanin patterning?

If the file is large, first scan all rows, group hits by query ID, and identify the most informative hits per query before interpreting the results. Do not base the interpretation on the first few rows only.

I am interested in several classes of genes, including but not limited to:

1. Core anthocyanin / flavonoid biosynthesis genes
   Examples: CHS, CHI, F3H, F3'H, FLS, DFR, ANS/LDOX, UFGT, OMTs, acyltransferases.

2. Anthocyanin regulators
   Examples: MYB, bHLH, WD40/TTG1, PAP1/MYB75, MYB90, MYB113, MYB114, GL3/EGL3/TT8, repressors such as MYBL2, SPLs, and other known flavonoid regulators.

3. Genes affecting anthocyanin localization, storage, stability, or visibility
   Examples: GSTs such as TT19, MATE/ABC transporters, vacuolar pH genes, proton pumps, ion transporters, vesicle trafficking, glutathione-related genes.

4. Genes plausibly involved in spatial patterning, epidermal patterning, or Turing-like / reaction-diffusion pattern formation
   Examples: genes involved in lateral inhibition, activator-inhibitor systems, epidermal cell fate, trichome/root hair patterning, mobile transcriptional regulators, hormone signaling, auxin/cytokinin/JA/ABA signaling, peptide signaling, receptor-like kinases, transporters, or genes with known roles in spatial prepatterning.

Please analyze the BLAST results and return only the strongest and most interpretable candidates, not every speculative idea.

For each predicted gene/query, consolidate multiple BLAST hits if they appear to point to the same biological function. Do not treat every hit independently if they are redundant.

Please produce the following output:

1. A ranked candidate table with columns:
   - Rank
   - Query ID
   - Best Arabidopsis hit or hits
   - Putative functional category
     Use one of:
       Core anthocyanin biosynthesis
       Anthocyanin regulation
       Pigment transport/storage/modification
       Pattern formation / spatial regulation
       Indirect but plausible pigmentation modifier
       Probably not relevant
   - Why this gene is relevant
   - Evidence strength: High / Medium / Low
   - Main BLAST evidence: percent identity, alignment length, E-value, bitscore
   - Pressure-test / caveat: why this could be a false lead
   - Suggested follow-up validation

2. A short “best bets” section:
   Identify the top 5–15 genes that are most worth following up, with concise reasoning.

3. A “do not overinterpret” section:
   List genes or categories that may sound relevant but should be treated cautiously, especially if the hit is short, low identity, generic, or belongs to a large nonspecific gene family.

4. A “missing expected genes” section:
   Note whether obvious anthocyanin pathway genes or regulators appear absent from the BLAST hits. Do not assume absence from the genome, only absence from this BLAST result file.

Important analysis rules:

- Prioritize biological relevance plus BLAST quality. A gene with a perfect functional annotation but weak BLAST support should not be ranked as high confidence.
- Be skeptical of very short alignments, generic annotations, large gene families, and “protein of unknown function” hits.
- Do not invent functions beyond what is supported by the Arabidopsis annotation and BLAST quality.
- For patterning genes, distinguish strong known patterning regulators from vague developmental genes.
- For Turing-like patterning, only flag genes that plausibly participate in spatial feedback, activator-inhibitor logic, mobile signaling, lateral inhibition, or pigment-domain formation. Do not label ordinary biosynthetic enzymes as Turing-pattern genes unless there is a clear regulatory or spatial reason.
- If a gene is relevant only indirectly, say so.
- Use cautious language: “candidate,” “possible,” “weakly supported,” “requires validation.”
- Return a concise, prioritized interpretation rather than a comprehensive list.
