## Supplementary figures and images for "Genetic architectures of floral pigment and patterning in hybrid monkeyflowers"

### Supplemental File 3

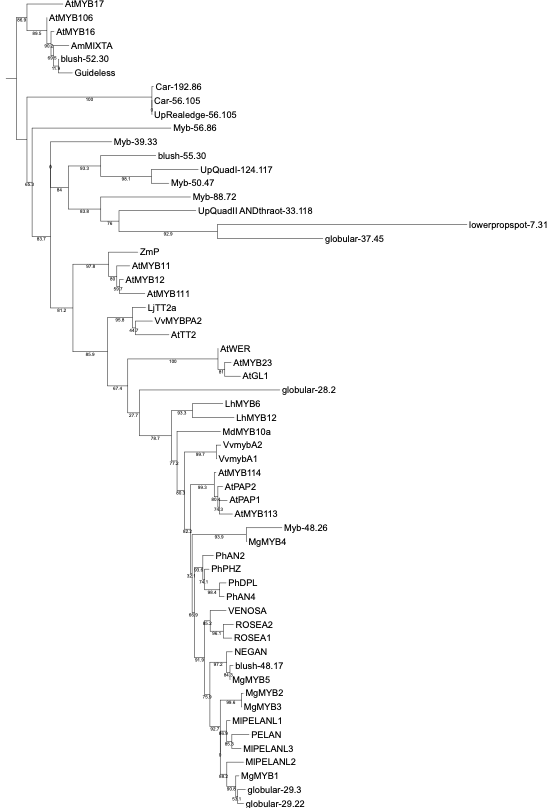

**Supplemental Figure S6** Tree of Myb sequences
