## Supplemental File 4 for "Genetic architectures of floral pigment and patterning in hybrid monkeyflowers"

**Supplemental Figures**

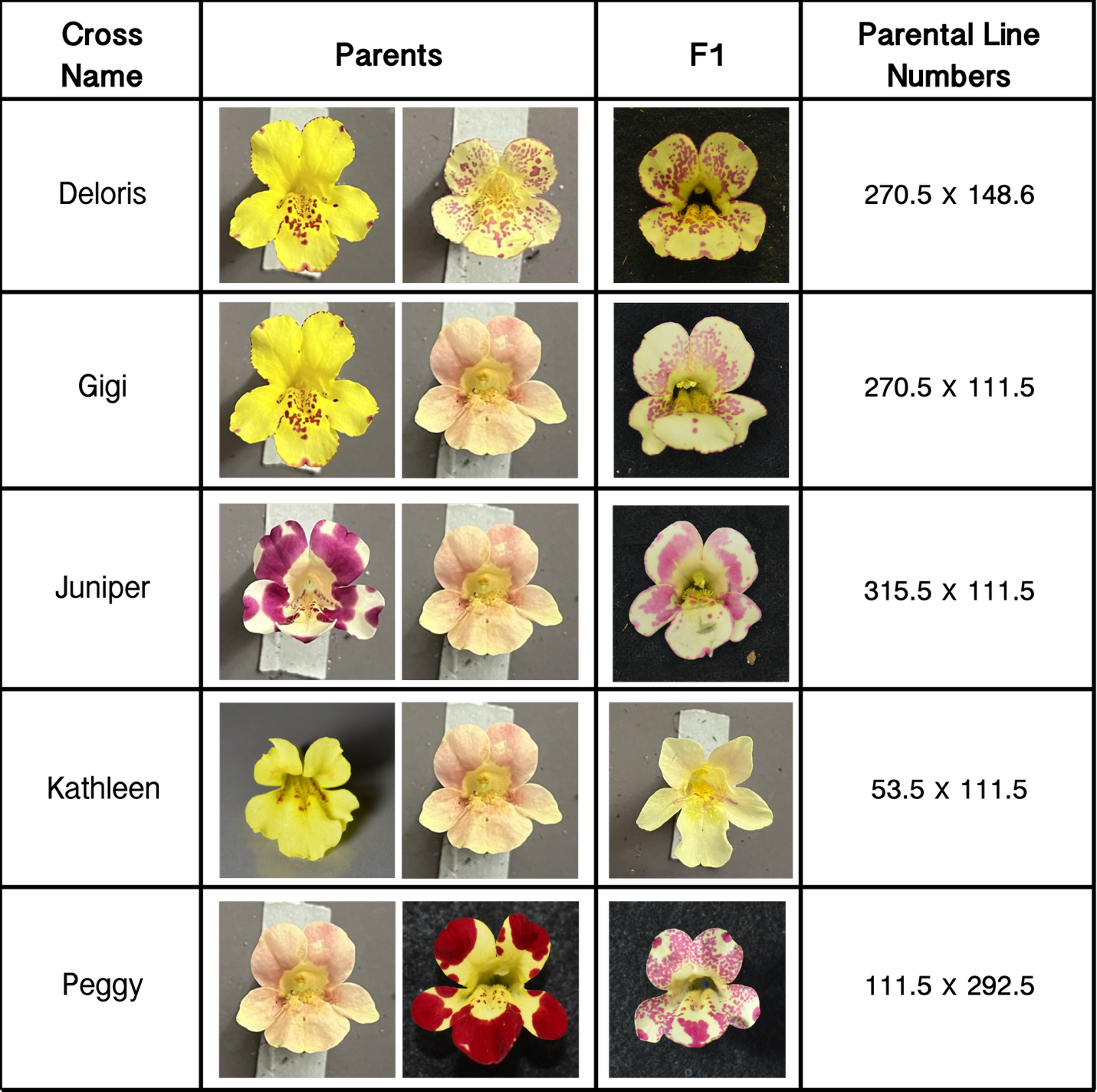

**Supplemental Figure S1** Summary of RIL crosses including cross name, photos of parental flowers, F1 flower photo, and RIL line information.

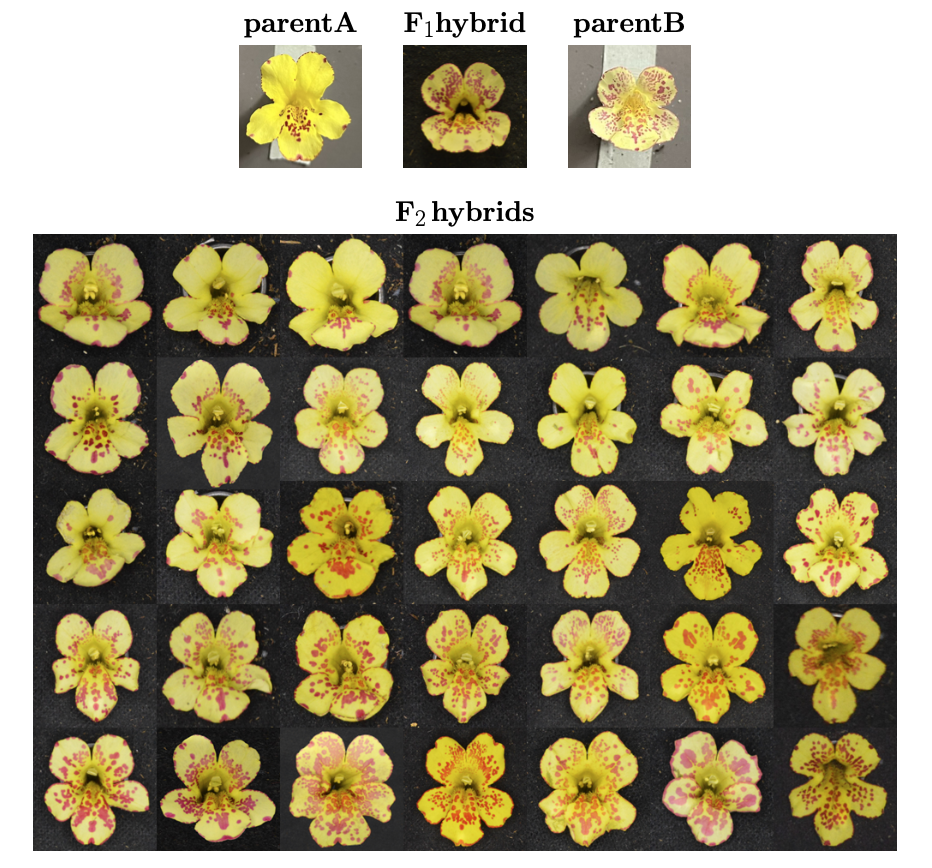
**Figure S2: Deloris cross** One individual with tip spots (parent A) is crossed with a speckled individual (parent B). Their F1 offspring bears tip spots and speckled petals. In the F2 generation, we see a distribution of phenotypes that do not easily segregate. Most F2 individuals exhibit some combination of parental traits. However, some individuals show traits not clearly seen in either parent. For example, the individual in row 5, column 6 (R5C6) exhibits large spots at in the center of its petals, while both parents show smaller spots. In R4C7, we see an individual with dense spotting near the throat, but only small tip spots on the distal side of the petals.

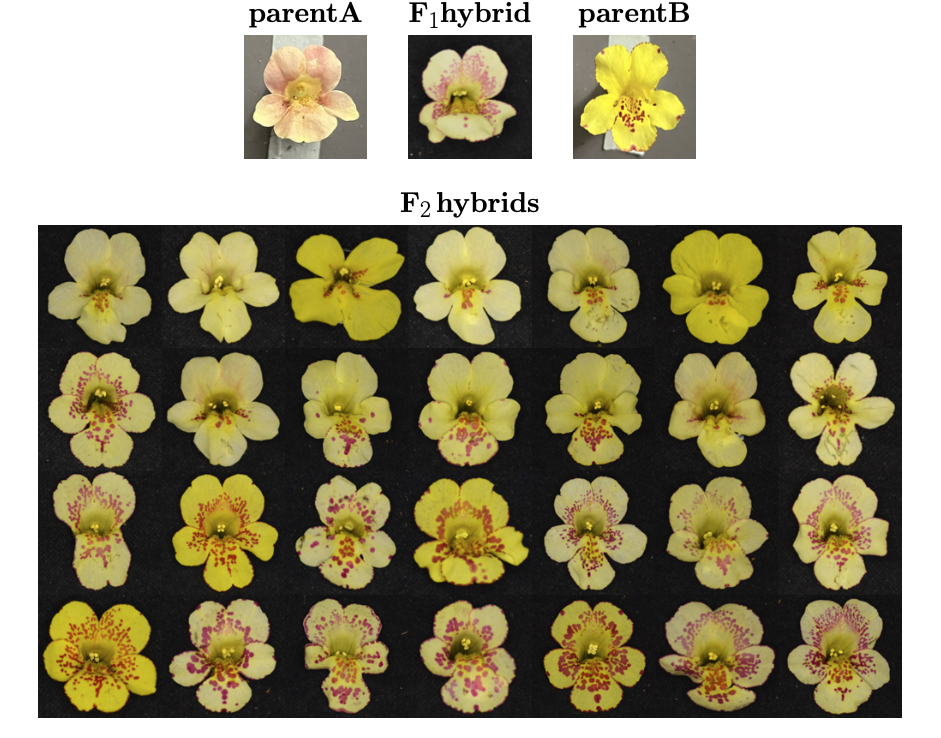

**Figure S3: Gigi cross.** One individual with high-coverage blush phenotype (parent A) is crossed with an individual with tip spots (parent B). The resulting F1 hybrid exhibits dense speckles emanating from the flower throat. In the F2 generation, the blush phenotype is recovered in a small subset of the population. Another subset of the population is devoid of anthocyanin, which should not be due to homozygous recessive activator alleles. Most F2 offspring exhibit a speckled phenotype similar to the F1 individual. The extent of speckling varies in a continuous, non-segregating fashion.

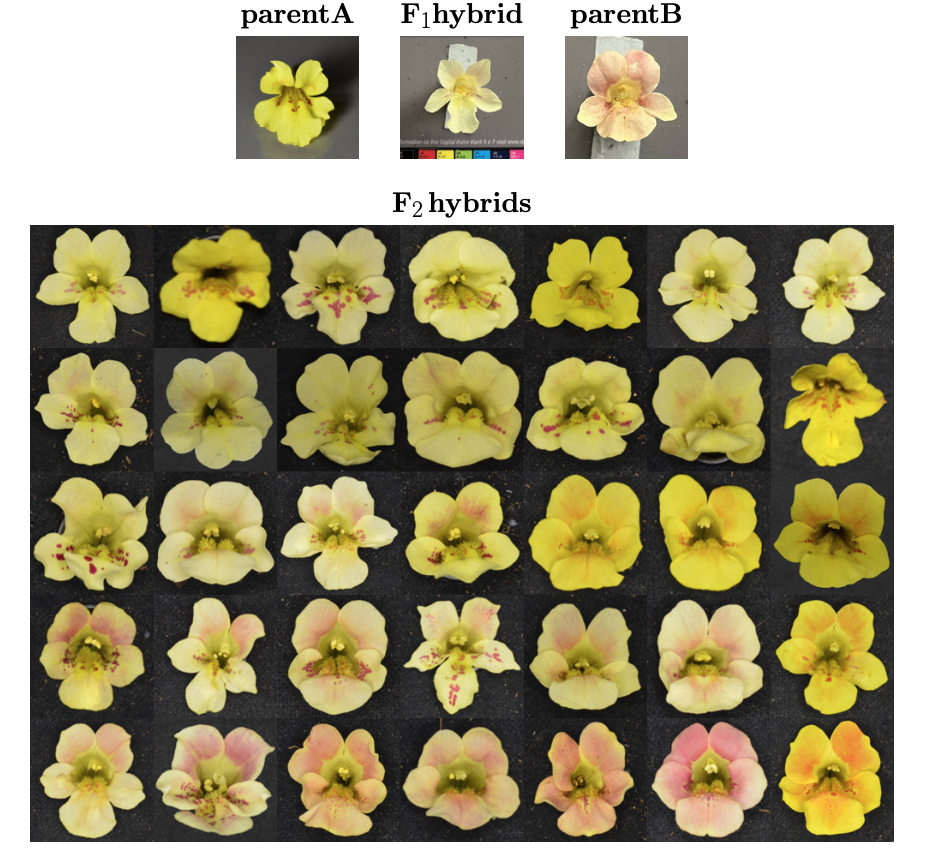

**Figure S4: Kathleen cross.** One individual with slight blush coverage (parent A) is crossed with high-coverage blush individual (parent B). The resulting F1 hybrid exhibits an intermediate level of blush coverage. In the F2 population, most individuals show a blush phenotype intermediate to the parents. Some individuals exhibit a spotted phenotype not found in either parent (R2C5 and R4C4). Another subset of the population is devoid of anthocyanin, which cannot be due to a homozygous recessive genotype for PELAN.

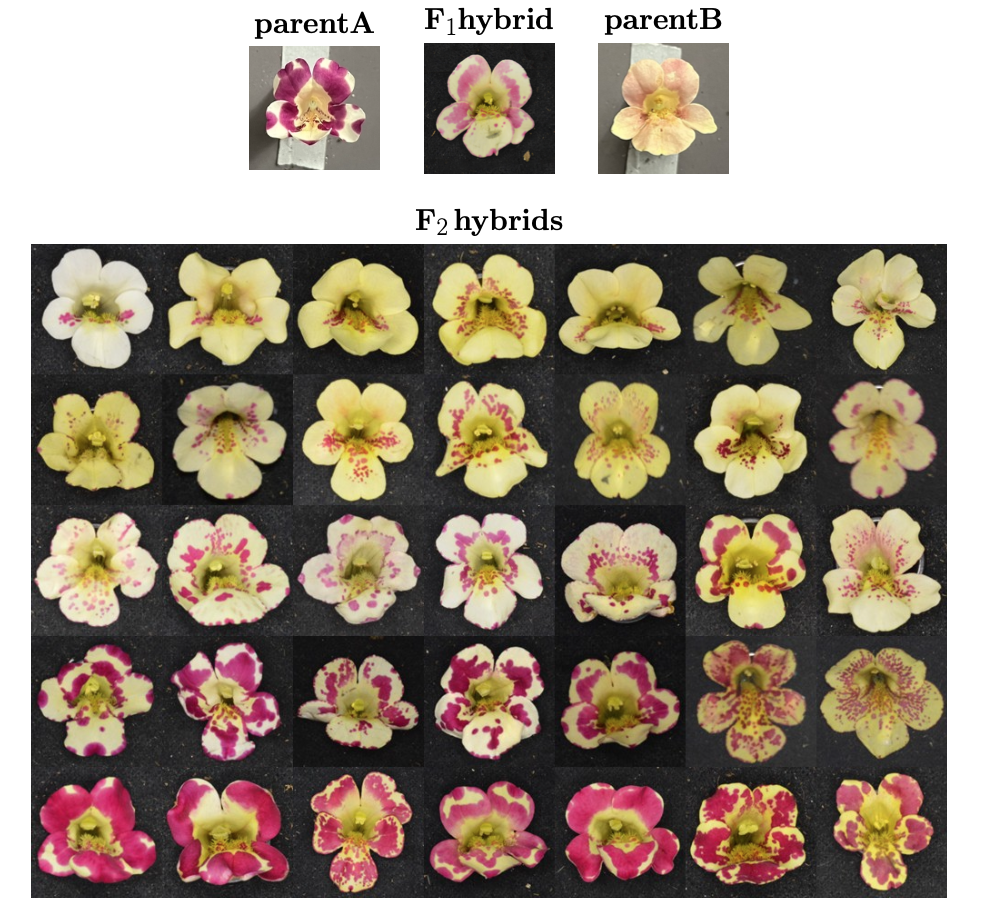

**Figure S5: Juniper cross.** A blotchy spotted individual (parent A) is crossed with a blush individual (parent B). The blotchy spots of parent A emanate from the tips of the petals and the flower throat. The resulting F1 hybrid shows blotchy spots emanating from the petal tips and flower throat, but with less coverage and lower pigment intensity than in parent A. In the F2 generation, there is a large diversity of petal phenotypes. In row one, many individuals are devoid or nearly devoid of anthocyanin. The F2 population shows a substantial number of individuals with speckled phenotypes varying in spot size, intensity, and coverage. Blush phenotype is recovered in a subset of the F2 population, but no full-blush individuals were observed. Several individuals (e.g., R3C7) show a phenotype that appears to blend spots and blush, which was not observed by this author outside of this cross.

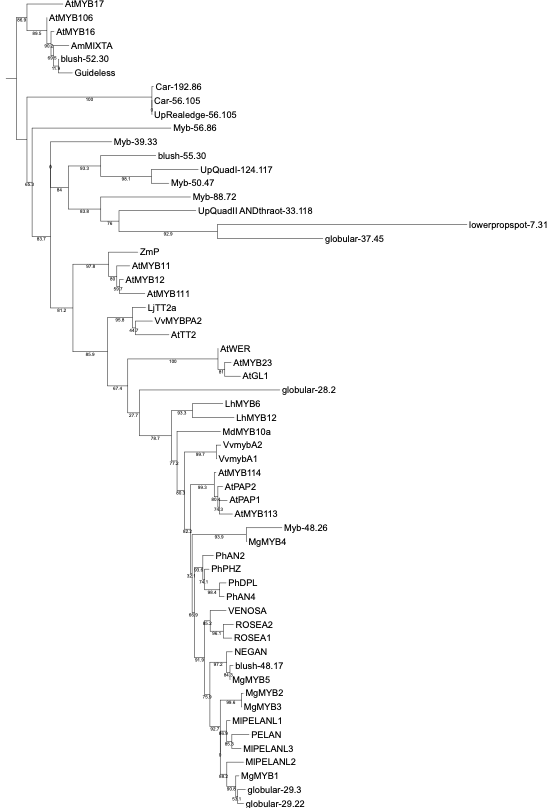

**Supplemental Figure S6** Tree of Myb sequences
